# A Novel Transcription Factor Combination for Direct Reprogramming to a Spontaneously Contracting Human Cardiomyocyte-like State

**DOI:** 10.1101/2023.03.14.532629

**Authors:** Marisol Romero-Tejeda, Hananeh Fonoudi, Carly J. Weddle, Jean-Marc DeKeyser, Brian Lenny, K. Ashley Fetterman, Tarek Magdy, Yadav Sapkota, Conrad Epting, Paul W. Burridge

## Abstract

The reprogramming of somatic cells to a spontaneously contracting cardiomyocyte-like state using defined transcription factors has proven successful in mouse fibroblasts. However, this process has been less successful in human cells, thus limiting the potential clinical applicability of this technology in regenerative medicine. We hypothesized that this issue is due to a lack of cross-species concordance between the required transcription factor combinations for mouse and human cells. To address this issue, we identified novel transcription factor candidates to induce cell conversion between human fibroblasts and cardiomyocytes, using the network-based algorithm Mogrify. We developed an automated, high-throughput method for screening transcription factor, small molecule, and growth factor combinations, utilizing acoustic liquid handling and high-content kinetic imaging cytometry. Using this high-throughput platform, we screened the effect of 4,960 unique transcription factor combinations on direct conversion of 24 patient-specific primary human cardiac fibroblast samples to cardiomyocytes. Our screen revealed the combination of *MYOCD*, *SMAD6*, and *TBX20* (MST) as the most successful direct reprogramming combination, which consistently produced up to 40% TNNT2^+^ cells in just 25 days. Addition of FGF2 and XAV939 to the MST cocktail resulted in reprogrammed cells with spontaneous contraction and cardiomyocyte-like calcium transients. Gene expression profiling of the reprogrammed cells also revealed the expression of cardiomyocyte associated genes. Together, these findings indicate that cardiac direct reprogramming in human cells can be achieved at similar levels to those attained in mouse fibroblasts. This progress represents a step forward towards the clinical application of the cardiac direct reprogramming approach.

**GRAPHICAL ABSTRACT:** 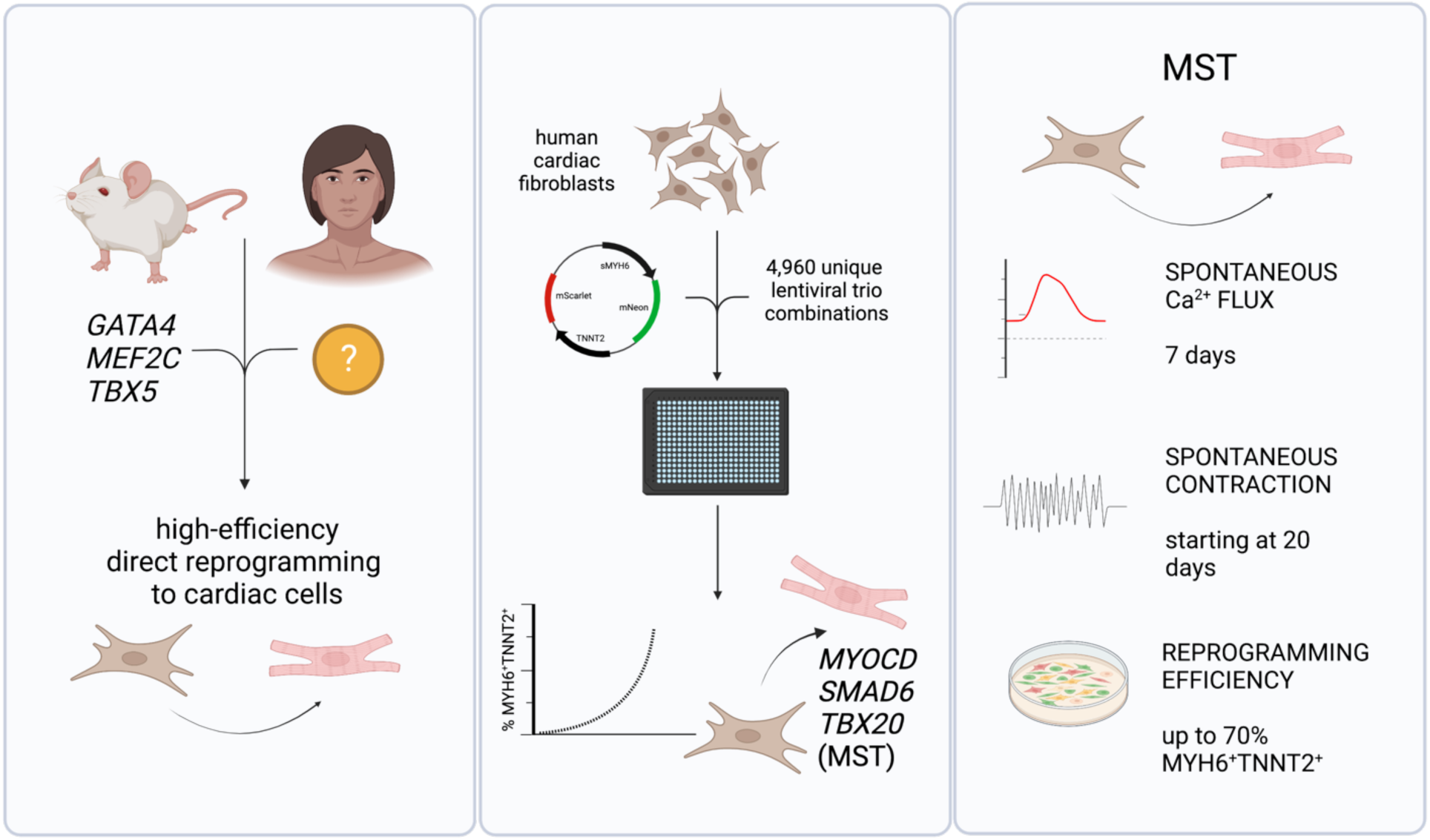

**HIGHLIGHTS:**

- Using network-based algorithm Mogrify, acoustic liquid handling, and high-content kinetic imaging cytometry we screened the effect of 4,960 unique transcription factor combinations.
- Using 24 patient-specific human fibroblast samples we identified the combination of *MYOCD*, *SMAD6*, and *TBX20* (MST) as the most successful direct reprogramming combination.
- MST cocktail results in reprogrammed cells with spontaneous contraction, cardiomyocyte-like calcium transients, and expression of cardiomyocyte associated genes.

## INTRODUCTION

The human heart is thought to lose approximately 1 billion cardiomyocytes after a myocardial infarction (MI)^1^. As human adult cardiomyocytes are only minimally proliferative (<1% per year)^2^, these lost cardiomyocytes cannot be replaced and instead a fibrotic scar is formed, resulting in reduced cardiac function. The development of approaches to replace the function of these lost cardiomyocytes is therefore key in recovery from MI and heart failure. Existing methods for cardiac regenerative medicine, such as engraftment of ‘cardiac progenitor cells,’ have largely either proven unsuccessful^3^ or, in the case of engraftment of human pluripotent stem cell-derived cardiomyocytes, will require immunosuppression and suppression of arrythmias^4^. Conversely, *in situ* direct reprogramming of cardiac fibroblasts to cardiomyocytes within the heart is potentially an ideal method to replace lost cardiomyocytes in the heart and provide functional recovery from MI or heart failure. However, despite advances in this field over the past decade, the direct reprogramming of human fibroblasts to cardiomyocytes remains inefficient compared to that of mouse.

In mouse cells, the combination of the transcription factors *Gata4*, *Mef2c*, and *Tbx5* (GMT) has been demonstrated by numerous groups to reprogram mouse fibroblasts to cardiomyocyte-like cells (iCMs)^5–10^, despite early skepticism^11,12^. The efficiency of this reprogramming has since been further enhanced by supplementing GMT with transcription factors such as *Hand2*^6,7,13,14^, *Nkx2-5*^7,13^*, Akt1*^14,15^*, Znf281*^15^*, or Myocd, Srf, Mesp1, and Smarcd3*^10^. Small molecules and microRNAs have likewise been used to augment reprogramming^8,9,13,16^ or to induce GMT expression^17^ from mouse fibroblasts.

In human cells, several groups have demonstrated that GMT alone is not sufficient for the reprogramming of human fibroblasts to human iCMs (hiCM). Studies show that addition of other transcription factors such as *MYOCD*, *HAND2*, *MESP1*, *ESSRG*, *ZFPM2*, *ASCL1*, *TBX20*, and/or *NKX2-5* are required. From these transcription factor-based cocktails, the combination of GMT plus *MYOCD* and *HAND2* (GMTM-H) has been demonstrated as the most efficient for reprogramming the human fibroblast to hiCM. GMTM-H generates 7.27-19.6% TNNT2-GFP^+^ cells after 2 weeks of reprogramming, depending on the fibroblast source^18^. The use of reprogramming enhancers, such as microRNAs *miR-1* or *miR-133*^9,18–22^ and small molecules^20^ have also been identified to improve the efficiency of human direct reprogramming, based on cardiac promoter activity and gene expression. GMT in the presence of miR-133 can generate 41.8% TNNT2^+^ cells after 2 weeks of reprogramming with antibiotic selection^22^. Furthermore, optimization of transcription factor stoichiometry using polycistronic vectors with a MEF2C-GATA4-TBX5 splicing order has further improved the reprogramming efficiency 10-fold in mouse cells^23,24^. Nonetheless, despite this progress, generation of hiCMs suffers from lower efficiency than mouse iCMs and the generation of spontaneously contracting cells has not been conclusively proven without co-culture with mouse CMs or hiPSC-CMs (**Table S1**). These difficulties suggest that there may be inter-species differences in the gene regulatory networks that control the fibroblast and/or cardiomyocyte states; therefore, alternative non-GMT-based transcription factor combinations are required for reprogramming.

Previous attempts to find transcription factor combinations to reprogram human fibroblasts to cardiomyocytes have been guided by low-throughput additive approaches using the mouse-discovered GMT as a starting combination and have used limited phenotypic measures of cardiomyocyte identity, such as expression of MYH6 or TNNT2, to measure reprogramming success. However, this ‘single gene’ methodology potentially may only optimize for transcription factor cocktails that primarily upregulate the expression of MYH6 or TNNT2 themselves rather than switch on the complete cardiomyocyte transcriptional network or generate hiCMs. Indeed, a combinatorial screen utilizing RT-qPCR to optimize for factors that upregulate simultaneous expression of 5 cardiac genes (*Myh6, Myl2, Actc1, Nkx2.5,* and *Scn5a*) in mouse embryonic fibroblasts revealed that the combination of *Tbx5*, *Mef2c*, and *Myocd* upregulated a broader range of cardiac genes compared to GMT^25^. Understandably, prior studies have selected candidate genes based solely on their high expression in human adult cardiomyocytes and their role during cardiac development. However, existing network-based algorithms that consider the transcription factor network already in place in the starting cell type, such as Mogrify^26^, could be used to identify candidate factors for direct reprogramming that specifically restructure the human fibroblast gene regulatory network; thus facilitating more informed transcription factor screens for hiCM generation.

Until now, high-throughput methods have not been applied to screen and identify reprogramming factors in primary human cells. A major limitation for performing high-throughput screens using human fibroblasts is cell availability. Different cell sources have been used in prior studies, including neonatal human foreskin fibroblasts^19^, human cardiac fibroblasts^18^, dermal fibroblasts^21^ or embryonic stem cell-derived fibroblasts^9,20^. However, due to the limited proliferative capacity of primary fibroblasts and the decrease in reprogramming efficiency^20^ over multiple passages, only immortalized fibroblasts have been used to perform high-throughput chemical screens for reprogramming enhancers^9^ in mouse fibroblasts. Since concordance between immortalized and primary cells has not been well studied, it is unclear whether the optimized condition can be directly translated to primary cells.

Another major barrier for performing high-throughput screening in human cells is the transfection strategy. Retroviruses are used to overexpress factors in most human fibroblast cardiac direct reprogramming protocols, as early work demonstrated these infect cells at >90% efficiency rather than the <20% achieved with lentiviruses^18^. The amphitropic retrovirus produced by PLAT-A cells^9,19,20^ is not able to withstand centrifugation or freeze/thaw cycles without up to a 100-fold reduction in viral titer; therefore, it must be made fresh prior to each experiment. Incorporating viral production into the reprogramming pipeline makes the addition of automation to the transcription factor screening process highly onerous. A major step forward was the use of more stable VSV-G pseudotyped doxycycline-inducible lentiviruses^21^ in HEK293T cells, which are compatible with freeze/thaw cycles, express factors at the same level as non-inducible constructs, and have been successfully used to generate cardiac-like cells from human fibroblasts^22^.

In this study, we developed a novel high throughput reprogramming strategy using low passage primary fibroblasts derived from 24 pediatric patients undergoing cardiac surgery. Cardiac, rather than dermal, fibroblasts were chosen as a starting cell type due to their central role in decreased heart function following MI, making these cells an attractive target for direct reprogramming *in vivo*. We have combined a lentiviral approach for reproducible gene introduction, a dual TNNT2-mScarlet MYH6-mNeonGreen reporter strategy, automated 384-well virus, small molecule, and growth factor dispensing, and high-content kinetic imaging cytometry to assess for both calcium cycling and cardiac reporter activity. Using this methodology, we found that variations of the traditional GMT-containing transcription factor combinations are not suitable for human direct cardiac reprogramming. Therefore, taking an unbiased approach, we used the Mogrify algorithm to identify new candidate transcription factors. Utilizing our optimized high-throughput screening methodology to assess reprogramming success, we found the combination of *MYOCD*, *SMAD6*, and *TBX20* (MST) as the most efficient combination. In just 6 days, MST produces 33% MYH6^+^ and 17% TNNT2^+^ cells on average and can produce upwards to 90% MYH6 ^+^ and 40% TNNT2^+^ cells. With the addition of a growth factor FGF2 and small molecule XAV939, these cells exhibited cardiomyocyte-like gene and protein expression, spontaneous calcium transients starting at 7 days, and spontaneous contraction at just 25 days. Our efficient reprogramming strategy can open new avenues for therapeutic application of hiCMs.

## METHODS

### Fibroblast isolation

Deidentified human heart samples were obtained from 24 patients aged 4 days – 21 years undergoing cardiac surgery with informed consent in accordance with Lurie Children’s Hospital Institutional Review Board (**Table S2**). Cardiac tissues were collected in RPMI 1640, transferred to a cell culture dish, and minced into small pieces (<1 mm^2^) using a sterile scalpel. Tissue pieces were then suspended in IMDM (Hyclone) supplemented with 20% FBS (VWR) and transferred to one well of a 6-well cell culture plate (Greiner) coated with a 1:800 dilution of growth factor-reduced Matrigel (Corning)^27^. All cells were kept at 37 °C and 5% CO_2_. Media was changed every 2 days beginning on day 5 and primary cardiac fibroblasts (PCFs) migrated out of the explants within 3-4 weeks. Migrated fibroblasts were passaged once they reached 90% confluence by rinsing with DPBS, then digesting in TrypLE Express (Gibco) for 5 minutes, and suspending with IMDM+20% FBS. Fibroblasts were filtered through a 100 µm cell strainer (Falcon) and transferred to Matrigel-coated 15 cm^2^ cell culture plates (Greiner). Cells were grown to 90% confluence and subsequently passaged (1:3 ratio) at least 2 more times before use in direct reprogramming experiments.

### Cardiac differentiation from human induced pluripotent stem cell-derived cardiomyocytes

Human induced pluripotent stem cell-derived cardiomyocytes (hiPSC-CMs) were used as positive controls. Differentiation of human induced pluripotent stem cells into cardiomyocytes was performed according to previously described protocol with slight modifications^28,29^. Briefly, hiPSCs were split at a 1:15 ratio using 0.5 mM EDTA and grown in B8 medium for 4 days reaching ∼80% confluence. At the start of differentiation (day 0), B8 medium was changed to R6C, consisting of RPMI 1640 (Corning, 10-040-CM), supplemented with 6 µM of glycogen synthase kinase 3-inhibitor CHIR99021 (LC Labs, C-6556). On day 1, medium was changed to RPMI 1640 basal medium alone, and on day 2 medium was changed to RBA-C59, consisting of RPMI 1640 supplemented with 2 mg/mL fatty acid-free bovine serum albumin (GenDEPOT, A0100), 200 µg/mL L-ascorbic acid 2-phosphate (Wako, 321-44823) and 2 µM Wnt-C59 (Biorbyt, orb181132). Medium was then changed on day 4 and then every other day with RBAI consisting of RPMI 1640 supplemented with 500 µg/mL fatty acid-free bovine serum albumin, 200 µg/mL L-ascorbic acid 2-phosphate, and 1 µg/mL *E. coli*-derived recombinant human insulin (Gibco, A11382IJ). Contracting cells were noted from day 8, differentiated cardiomyocytes were treated with 100 µg/mL of neomycin from day 8 to day 12. On day 16 of differentiation, cardiomyocytes were dissociated using DPBS for 20 min at 37 °C followed by 1:200 Liberase TH (Roche, 5401151001) diluted in DPBS for 20 min at 37 °C, centrifuged at 300 × g for 5 min, and filtered through a 100 µm cell strainer (Falcon). Cells were then plated in RBAI+10% Cosmic Calf Serum (Hyclone) for 2 days on 1:800 Matrigel-coated plates for each assay, media was then switched back to RBAI which was changed every 2-3 days and cells were assayed on day 30.

### Flow cytometry

For flow cytometry analysis, fibroblasts were dissociated from cell culture dishes using TrypLE Express (Gibco) for 5 minutes and hiPSC-CMs were dissociated using 1:200 Liberase TH (Roche) in PBS for 20 min. 1 × 10^6^ cells were transferred to round bottom FACS tubes (Falcon) and fixed with 4% PFA in DPBS (100 µL) for 20 min. For analysis of intracellular markers, cells were washed twice using DPBS and permeabilized in 100 µL of DPBS with 0.5% BSA and 0.5% saponin for 15 min. Once fixed and permeabilized, cells were stained for 2 h at room temperature and washed prior to data collection. For analysis of extracellular markers, fixed cells were washed twice and stained for 30 min on ice prior to data collection. Antibodies used are listed in **Table S3.** All data were collected using a CytoFLEX flow cytometer (Beckman Coulter) and analyzed using CytExpert 2.2 software (Beckman Coulter) using isotype controls to set positive gates.

### Plasmids

All plasmids used in this study have been deposited in Addgene. To generate doxycycline-inducible lentiviral plasmids for overexpression of candidate factors, ORF cDNA was inserted into the FU-tetO-Gateway (Addgene 43914) backbone using Gateway cloning. First, attB sites were added to factor ORF cDNA via PCR reaction. We amplified the N-terminal region of each gene in our candidate pool by designing forward primers containing the attB1 sequence, Shine-Dalgarno sequence, and Kozak sequence upstream of the transcriptional start site, as well as overlap to the cDNA ORF. We designed reverse primers to amplify the C-terminal of each gene and to include the attB2 site. Primers used for PCR reactions are listed in **Table S4.** PCR reactions were performed by combining 12.5 µL Q5 high-fidelity 2× master mix (NEB), 2.5 µL each of 10 µM forward and reverse primers, and 1 µL of 1 ng/µL of DNA template (**Table S5**). Cycling conditions were as follows: initialization at 98 °C for 30s, followed by 35 cycles of denaturation at 98 °C for 10 seconds, annealing at 60 °C for 10 seconds, and elongation at 72 °C for 1s per KB of template, ending with a final elongation at 72 °C for 5 minutes and a hold at 4 °C. Resulting fragments were purified and recombined with Gateway pDONR221 Vector (Invitrogen) using Gateway BP Clonase II enzyme mix (Invitrogen) to generate entry clones. Purification of attB-containing PCR fragments was performed using PEG 8000/MgCl2, following the Gateway Clonase II enzyme mix protocol. The BP reaction was performed by combining 150 ng of attB-PCR product, 150 ng of pDONR221 vector, and 2 µL of BP Clonase II in TE buffer to a total volume of 8 µL. DNA was allowed to recombine overnight at 25 °C before adding 1 µL of Proteinase K and incubating at 37 °C for 10 min. We then transformed One Shot TOP10 chemically competent *E. col*i cells (Invitrogen) with 1 µL of BP product following the manufacturer’s recommended protocol. Transformed bacteria were plated on LB agar plates with Kanamycin (50 µg/mL) and surviving entry clones were selected. We purified plasmid DNA using the ZymoPURE Plasmid Miniprep Kit (Zymo Research) following the manufacturer’s protocol. Gateway LR Clonase II Enzyme mix (Invitrogen) was used to catalyze recombination of entry clones with the FU-tetO-Gateway backbone (Addgene plasmid # 43914) using 150 ng of pDONR221-cDNA plasmid, 150 ng of FU-tetO-Gateway plasmid, and 2 µL of LR Clonase II in TE buffer to a total volume of 8 µL. DNA was allowed to recombine, and expression clones were selected using ampicillin (100 μg/mL), following the same protocol described above. Expression plasmids were verified by Sanger sequencing using the T7 forward and GW_R reverse primers (**Table S4**). All expression plasmids were propagated in Stbl2 or NEB Stable bacteria to reduce the occurrence of recombination due to long terminal repeats. Plasmid maps are found in **Figure S1**.

### Lentiviral production

To generate lentivirus, Lenti-X 293T packaging cells (Takara, 632180) (passage 2 – 3) were thawed and allowed to recover in DMEM with 10% FBS (Avantor Seradigm, VWR) and 4 mM L-alanyl-L-glutamine (DMEM complete) for 4 days before transection with lentiviral plasmids. After recovery, cells were dissociated using TrypLE Express (Gibco), seeded into uncoated 10 cm^2^ cell culture plates at 3.8 million cells per plate in 15 mL of DMEM complete and incubated for 20 h (37 °C, 5% CO_2_). The following day, media was replaced with 10 mL of DMEM complete with 25 µM cloroquine diphosphate and incubated for 5 h. To transfect cells, we prepared a mixture of 1.3 pmol psPAX2 (Addgene 12260), 0.72 pmol pMD2.G (Addgene 12259), and 1.64 pmol of transfer plasmid (tet-O-cDNA) in 500 µL of OptiPro SFM (Gibco). Polyethyenimine (PEI) (Polysciences, 23966-1) was used as a transfection reagent. In a separate tube, PEI (1 mg/mL) was further diluted with OptiPro SFM at a DNA to PEI ratio of 1:2 to a final volume of 500 µL. The diluted PEI solution was then added dropwise to the diluted DNA. The mixture was incubated at room temperature for 10 min before being added dropwise to the Lenti-X 293T cells in culture. A media change with 15 mL of DMEM complete was performed the following day. Cells were kept in culture and lentiviral supernatant was collected at 48 h post-transfection and stored at 4 °C until the second collection of supernatant at 72 h post-transfection. The 30 mL of lentiviral supernatant was pooled and concentrated to 1/100^th^ of its original volume (300 µL) in DPBS using PEG-iT (System Biosciences), following the manufacturer’s protocol. Lentiviral titration was performed by RT-qPCR using the Lenti-X qRT-PCR Titration Kit (Takara) and the concentration of lentivirus used for this paper was found to be above 1 × 10^7^ copies / mL. Concentrated lentivirus was aliquoted and stored at -80 °C until use.

### Lentiviral transduction

All PCFs used for direct reprogramming were propagated in culture (37 °C, 5% CO_2_) using 15 cm^2^ cell culture plates coated in 1:800 growth factor-reduced Matrigel (Corning)^27^ and IMDM (Hyclone) supplemented with 20% FBS (VWR). On reprogramming day -1, PCFs were dissociated using TrypLE Express and plated at 600,000 cells per well in Matrigel-coated, non-treated 6-well plates in DMEM+10% FBS (GenDEPOT Opti-Gold) media. To achieve high transduction efficiency, cells were allowed to attach to the plate surface for 2-3 hours and were subsequently transduced using TransDux Max (SBI). For lentiviral transduction, media in each well was aspirated and replaced with 1.5 mL of DMEM+10% FBS media containing TransDux Max (1x), 16 µL of concentrated FUdeltaGW-rtTA lentivirus (7.2 × 10^7^ copies / mL) and 16 µL of concentrated pFU-GW-sMYH6-mScarlet-TNNT2-mNeon (1.84 × 10^7^ copies / mL). Cells were incubated overnight. On reprogramming day 0, cells were dissociated using TrypLE Express, pooled, and replated for reprogramming in 25 µL of DMEM 10% FBS media containing TransDux Max (1x) at 3,000 cells per well in a Matrigel-coated µClear flat bottom black 384-well plates (Greiner) (cell numbers and media volumes were scaled according to surface area for other plate formats). Cells were allowed to attach to the plate surface for 2-3 h before transduction. Plates were briefly centrifuged and 50 nL of each concentrated lentivirus for transcription factor overexpression was dispensed into wells using Echo Acoustic Liquid Handling (LabCyte). Plates were incubated overnight. On reprogramming day 1, a media change was performed using 25 µL of DMEM 10% FBS containing 0.25 µg/mL doxycycline (Sigma-Aldrich). Media was replaced every two days. For reprogramming with small molecules, 5 μM XAV939 (APExBIO) was added to the media starting on reprogramming day 3. In experiments using reprogramming media, DMEM supplemented with 5 mg/mL fatty acid-free bovine serum albumin (BSA), 40 µg/mL ascorbic acid 2-phosphate, 20 µg/mL insulin, 5 µg/mL transferrin, and 7 µg/L sodium selenite was used starting on reprogramming day 1.

### Reporter Fluorescence Quantification

We quantified the fluorescence of cells transduced with pFU-GW-sMYH6-mScarlet-TNNT2-mNeon (Addgene 170712) and ORF cDNA-expressing transcription factors on reprogramming day 6. Media in μClear bottom black 384-well plates (Greiner) was replaced with 25 µL of FluoroBrite DMEM (Thermo Fisher) supplemented with 4 mM L-alanyl-L-glutamine and NucBlue nucleus staining reagent (1 drop per mL) (Invitrogen). Plates were incubated for 15 min (37 °C, 5% CO_2_) prior to capturing one image per well using high-content image cytometry (KIC, Vala Sciences). For image analysis, we used an algorithm developed by Vala Sciences to identify nuclear bounds and quantify their FITC and TRITC average pixel intensity (API). We used negative control wells, containing fibroblasts transduced with pFU-GW-sMYH6-mScarlet-TNNT2-mNeon and rTTA alone, to set a negative API threshold. Cells with API above the negative threshold in both the FITC and TRITC channel were counted as positive. Percent positive cells was calculated relative to the total number of nuclei in the well. All experiments were performed with at least three technical replicates and at least three separate patient samples.

### Calcium Imaging

We assayed calcium flux in cells on days 7-30 of reprogramming. For calcium imaging, media in flat bottom black 384-well plates (Greiner) was replaced with 25 µL of HBSS (Gibco) supplemented with 0.02 M HEPES (Fisher Scientific), 2.5 mM probenecid (Sigma), 0.04% pluronic F-127 (Sigma), and 10 µM Cal-520 (AAT Bioquest). Following a 1 h incubation (37 °C, 5% CO_2_), media was aspirated and replaced with 25 µL of FluoroBrite DMEM (Gibco) supplemented with 4 mM L-alanyl-L-glutamine and NucBlue Live ReadyProbes Reagent (1 drop per mL) (Invitrogen). Plates were incubated for an additional 10 min (37 °C, 5% CO_2_) prior to data collection using high-content image cytometry (KIC, Vala Sciences). Images were captured at in the FITC channel for 60 s (50 FPS, 10 ms exposure). We then used an algorithm developed by Vala Sciences to quantify FITC pixel intensity within nuclear and cell borders and to analyze trace parameters.

### Stimulation

Reprogrammed cells were differentiated in 384-well plates and stained with Cal-520 and NucBlue Live ReadyProbes Reagent as described above. Following initial data collection, cells were stimulated with 2 µM isoproterenol (Sigma) and incubated for 10 min (37 °C, 5% CO_2_) prior to data collection.

### Quantitative Real-time PCR

Cell lysates for RT-qPCR were collected in 300 µL TRIzol (Invitrogen) and their total RNA was isolated using Direct-zol RNA Microprep kit (Zymo Research), following the manufacturer’s protocol. Reverse transcription was performed from 0.5 µg – 1 µg of RNA using Maxima H Minus master mix (Thermo), following the manufacturer’s protocol. cDNA was then diluted 1:10 for 1 µg starting RNA or 1:5 for 0.5 µg starting RNA. RT-qPCR was performed using Taqman gene expression assays (Applied Biosystems) (**Table S6**) and Taqman Fast Advanced Master Mix (Applied Biosystems), following the manufacturer’s protocol. All PCR reactions were prepared in triplicate in a 384-well format. Data was collected using the QuantStudio 5 Real-Time PCR system. Relative quantification of gene expression was calculated using the 2^-ΛΛCt^ method^30^, normalized to the reference 18S and fibroblast control sample, as specified in the figure legends.

### Immunostaining

Reprogrammed fibroblasts were dissociated at day 30 and plated at 10,000 cells/well of Matrigel-treated Nunc Lab-Tek II 8-chamber slides. After 3 days, cells were fixed with 4% PFA (Electron Microscope Sciences, 15713S) in DPBS for 20 min at RT followed by permeabilization with 0.3% Triton X-100 (Fisher Bioreagents, BP151-100) in DPBS for 10 min at RT. Cells were then blocked with 1 mg/mL BSA in DPBS for 60 min at RT, and stained with rabbit polyclonal IgG TNNT2 (1:200, abcam, ab45932) and mouse monoclonal IgG1 a-Actinin (1:1000, Sigma-Aldrich, A7811) primary antibodies in 1 mg/mL BSA plus 0.1% Tween (Fisher, BP337-100) for 2 h at RT. Cells were then washed for 5 min with DPBS three times, then stained with secondary antibodies Alexa Fluor 488 Goat anti Rabbit IgG (1:500, Invitrogen, A11012) and Alexa Fluor 594 Goat anti Mouse IgG1(1:500, Invitrogen, A21125) in 1 mg/mL BSA plus 0.1% Tween (Fisher, BP337-100) for 2 h at RT. Cells were washed three times for 10 min with 0.1% Tween in DPBS and mounted with ProLong Diamond Antifade Mountant with DAPI (Invitrogen, P36962). Slides were imaged with a Ti-E inverted fluorescence microscope (Nikon Instruments) using NIS-Elements software.

### RNA-seq

Three samples of un-transduced cardiac fibroblasts, MST transduced cardiac fibroblasts, and hiPSC-derived cardiomyocytes at d30 were prepared. RNA was extracted using 150 µL/well of TRIzol Reagent (Thermo Fisher, 15596026). RNA was then purified using the Direct-zol RNA MicroPrep kit (Zymo, R2062) including on-column DNase digestion to remove genomic DNA. Samples were quantified using a Thermo Scientific NanoDrop 8000 and passed QC, and were then shipped in dry ice for library preparation and sequencing

Paired-end (100 bp) RNA-sequencing on hiPSCs was performed using the DNBSEQ sequencing platform at Beijing Genomics Institute. Total RNA was extracted and oligo dT beads were used to enrich mRNA with poly A tail using DNBSEQ Eukaryotic strand-specific mRNA library protocol. mRNA molecules were fragmented into small pieces and the fragmented mRNA was synthesized into first strand cDNA using random primers. The second strand cDNA was synthesized using dUTP instead of dTTP. The synthesized cDNA was subjected to end-repair and 3’ adenylated and adaptors were ligated to the ends of these 3’ adenylated cDNA fragments. The U-labelled second-strand template was digested with Uracil-DNA-Glycosylase (UDG) and PCR amplification was performed. Following library quality control and circularization, the library was amplified to make DNA nanoball (DNB) and sequenced on DNBSEQ platform.

Using SOAPnuke^31^, raw fastq files were processed to trim/remove adaptors, low-quality reads, and N reads. Clean sequencing reads were mapped to the GRCh38 reference genome using STAR^32^ and counted using RSEM^33^. Genes with less than 10 counts across all 9 samples were excluded from subsequent analysis. Following variance stabilizing transformation in DESeq2 R package^34^, principal component analysis was performed to visualize the clustering of samples. Differential expression analysis was performed using the DESeq2 R package^34^ accounting for the matched pair design. Results were shown by a volcano plot generated using the ggplot2 R package^35^. Genes with adjusted *P*-value <0.05 corrected for multiple testing using the Benjamini Hochberg method were considered as differentially expressed. Using the most strongly differentially expressed genes between hiPSC-CM and PCF, and between MST-reprogrammed hiCM and PCF, hierarchical clustering based on Euclidean distance was performed by the pheatmap R package^36^. Ingenuity Pathway Analysis (IPA) was performed to identify canonical pathways based on the differentially expressed genes (FDR <0.005)^37^. This was used to identify differences in pathways related to biological processes that were relevant between our samples.

To identify relevant gene sets, Gene Set Enrichment Analysis^38^ was performed. Outputs from DESeq2 analysis for hiPSC-CM vs PCF and MST-reprogrammed hiCM vs PCF comparisons were used as inputs for GSEA analysis. Human curated gene sets (C2) and cell type signature gene sets (C8) were the target gene sets. Results with FDR < 0.05 were considered as statistically significant.

### Statistics

24 primary samples were used to generate data for this study. Specific samples used for each experiment are detailed in **Table S2** and sample size is indicated in each figure legend. All statistical analysis was performed using GraphPad Prism 9 software. Data are shown as mean ± standard deviation. Student’s t-test was used for pairwise comparisons in most instances, as indicated in figure legends. However, because gene expression data from RT-qPCR analysis was not normally distributed, a non-parametric Mann-Whitney test was performed. Comparisons among 3 or more groups were performed using One-way ANOVA with subsequent pairwise testing via t-test with Tukey’s adjustment for multiple comparisons or Fisher’s LSD test.

## RESULTS

### Platform Optimization for transcription factor screening

We first developed a lentiviral transduction and reporter platform to allow for high-throughput screening of candidate transcription factors for direct reprogramming in primary human cardiac fibroblasts (PCFs) from pediatric patients (**Figure 1A**). These PCFs expressed typical fibroblast markers CD90 and vimentin and did not express endothelial cell marker CD31 by flow cytometry (**Figure S2A**). PCFs expressed fibroblast markers and did not express cardiomyocyte markers by RT-qPCR. hiPSC-derived cardiomyocytes were used as a positive control (**Figure S2B**). To facilitate the screening, PCFs were transduced with a dual-fluorescence reporter (pFU-GW-sMYH6-mScarlet-TNNT2-mNeon), validated using human induced pluripotent stem cell-derived cardiomyocytes (hiPSC-CMs) (**Figure S3A**). PCFs were then transduced in multiplex using only 50 nL of concentrated doxycycline-inducible lentivirus (1 x 10^8-10^ copies/mL), delivered in 384-well format via liquid handling, resulting in >90% transduction efficiency (**Figure S3B**). Cell seeding density (3,000 cells per well), MOI (0.17), basal media (DMEM+10% FBS), and doxycycline dose (0.25 µg/mL) were optimized for our system as described in **Figure S3C-S3F**. Reporter activity was measured using kinetic image cytometry on reprogramming day 6, thus facilitating high-throughput screens.

**Figure 1.**
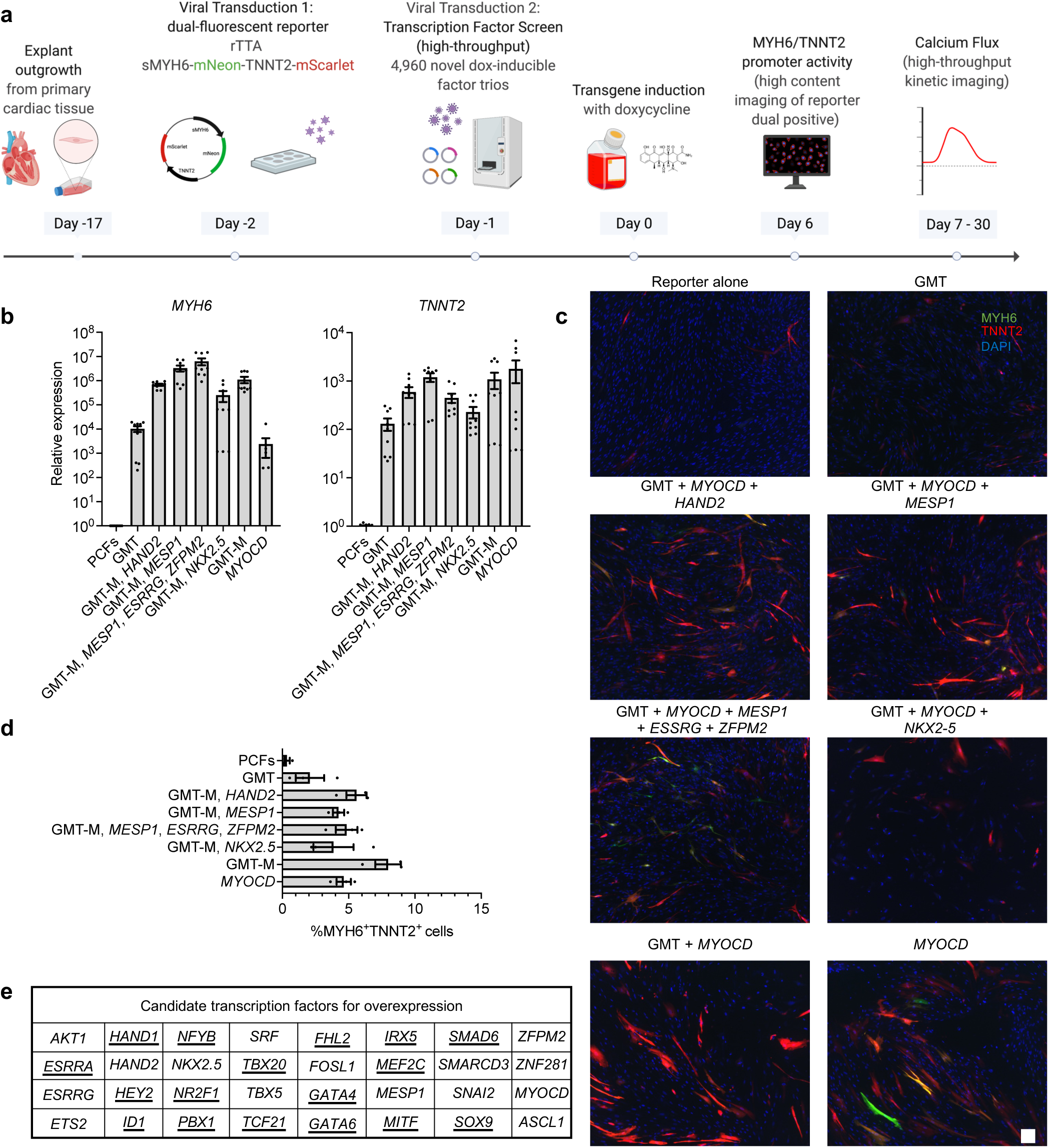
Establishment of a high throughput platform to assess the effect of transcription factor combinations on cardiac direct reprogramming. (**a**) Lentiviral transduction and reporter platform: human cardiac fibroblasts are obtained from discarded primary human cardiac tissue by explant outgrowth. Cells are collected and transduced overnight with rTTA and a MYH6/TNNT2 dual cardiac reporter. The following day, cells are collected and replated. Lentivirus combinations are delivered in multiplex using an acoustic liquid handler. Media containing doxycycline is changed every two days beginning on day two. Fluorescence from cardiac reporters is measured using a high-content kinetic imager 6 days after doxycycline addition. (**b**) Relative expression of *MYH6* and *TNNT2* measured by RT-qPCR in primary fibroblasts transduced only with existing cardiac reprogramming factor cocktails at reprogramming day 6. Expression normalized to PCF control. (*n* = 3 biological replicates, 3 technical replicates each). (**c**) Representative images of primary cardiac fibroblasts transduced with sMYH6-mNeonGreen-TNNT2-mScarlet and reprogrammed using published factor combinations. Scale bar = 100 μM. Images acquired using high content image cytometry at reprogramming day 6 (KIC, VALA Sciences) (**d**) Quantification of MYH6^+^TNNT2^+^ cells as measured by pFU-GW-sMYH6-mScarlet-TNNT2-mNeon reporter fluorescence on reprogramming day 6 (*n* = 3 biological replicates). All cells were subject to reporter overexpression (**e**) Candidate transcription factors for overexpression. Factors predicted by Mogrify underlined. Primary cardiac fibroblasts (PCF), *GATA4, MEF2C, TBX5* (GMT), GMT *+ MYOCD* (GMT-M). Error bars: mean ± SEM.

To validate our workflow and the ability of our lentiviral system to induce differentiation towards the cardiac fate in the absence of a transgenic reporter, we transduced PCFs with previously published transcription-factor based cardiac reprogramming cocktails (**Table S1**). As expected, we observed upregulation of endogenous *MYH6* and *TNNT2* transcripts by RT-qPCR within 6 days of reprogramming with each of the tested published factor combinations, (**Figure 1B**). Likewise, the tested combinations induced activation of our dual MYH6/TNNT2 reporter; however, simultaneous cardiac-specific MYH6 and TNNT2 activity was detected in only 2-8% of transduced cells (**Figure 1C-1D). *MYH6* activity was detected in 3-9% of cells and *TNNT2* activity was detected in 4-12% of cells (Figure S4A-S4C**). No reporter activity was observed in control cells (PCFs) transduced with rTTA and reporter alone. Interestingly, overexpression of *MYOCD* alone led to upregulation of endogenous TNNT2 and, to a lesser extent, MYH6, as measured by RT-qPCR as well as activation of our dual fluorescent reporter (**Figure 1B-D**). This finding is consistent with reports of partial activation of the cardiac gene network, including aMHC and cTnT, by overexpression of *MYOCD* alone in human foreskin fibroblasts^39^ and in mouse fibroblasts^40^.

Nonetheless, despite cardiac promoter activation in cells obtained using our optimized platform, none of the published transcription factor combinations tested was sufficient to generate spontaneously beating cells within a 4-week timeframe.

### Identification of novel transcription factor combination *MYOCD*, *SMAD6*, *TBX20* for direct cardiac reprogramming

To increase reprogramming efficiency and generate functional reprogrammed cardiomyocytes, we next used the network-based algorithm Mogrify^26^ to identify a novel set of transcription factors whose overexpression is predicted to result in the cardiac reprogramming of PCFs (**Figure S5**). Additionally, transcription factors which were previously reported to reprogram fibroblasts into cardiomyocytes, as well as factors that have been reported to increase cardiac direct reprogramming efficiency were also included in our candidate list (**Figure 1F, Table S5**). We generated doxycycline-inducible lentiviral constructs for overexpression of the 32 candidate factors and used our platform to assay the reprogramming efficiency of all factor trios in PCFs expanded from each of 3 distinct primary samples. Our initial screen of a total of 4,960 distinct combinations identified several factor trios with higher reprogramming efficiency compared to GMT-containing cocktails or *MYOCD* overexpression alone (**Figure 2A, Table S7**). To ensure reproducibility of the identified combinations, we validated our top 50 hits by transducing PCFs from 13 primary samples in triplicate to identify the top 3 combinations resulting from our screen: (1) *MYOCD*, *SMAD6*, *TBX20* (MST); (2) *MYOCD*, *SMAD6*, *PBX1* (MSP); and (3) *MYOCD*, *SMAD6*, *SNAI2* (MSS) (**Figure 2B**). While reprogramming success among experiments and samples is highly variable, each of these combinations generated MYH6^+^TNNT2^+^ cells at higher efficiency (up to 70%, 13.3% on average) compared to GMT-based cocktails (up to 20%, 1-5% on average) and to *MYOCD* overexpression alone (up to 21%, 6% on average). (**Figure 2C**) Although MST, MSP, and MSS all produce similar percentages of TNNT2^+^ cells (∼20%), MST produces significantly more MYH6^+^ cells (∼30%) from which up to 80% are MYH6^+^TNNT2^+^ cells, as measured by reporter expression (**Figure 2C**). The potential of the MST combination to reprogram PCFs with high efficiency is illustrated in **Fig. 2D**. Likewise, MST-derived cells express higher levels of TNNT2 protein compared to MSP and MSS and *MYOCD* overexpression alone (**Figure S6**) and the combination gives rise to approximately 40% TNNT2^+^ by flow cytometry at reprogramming day 15 (**Figure 2E**).

**Figure 2.**
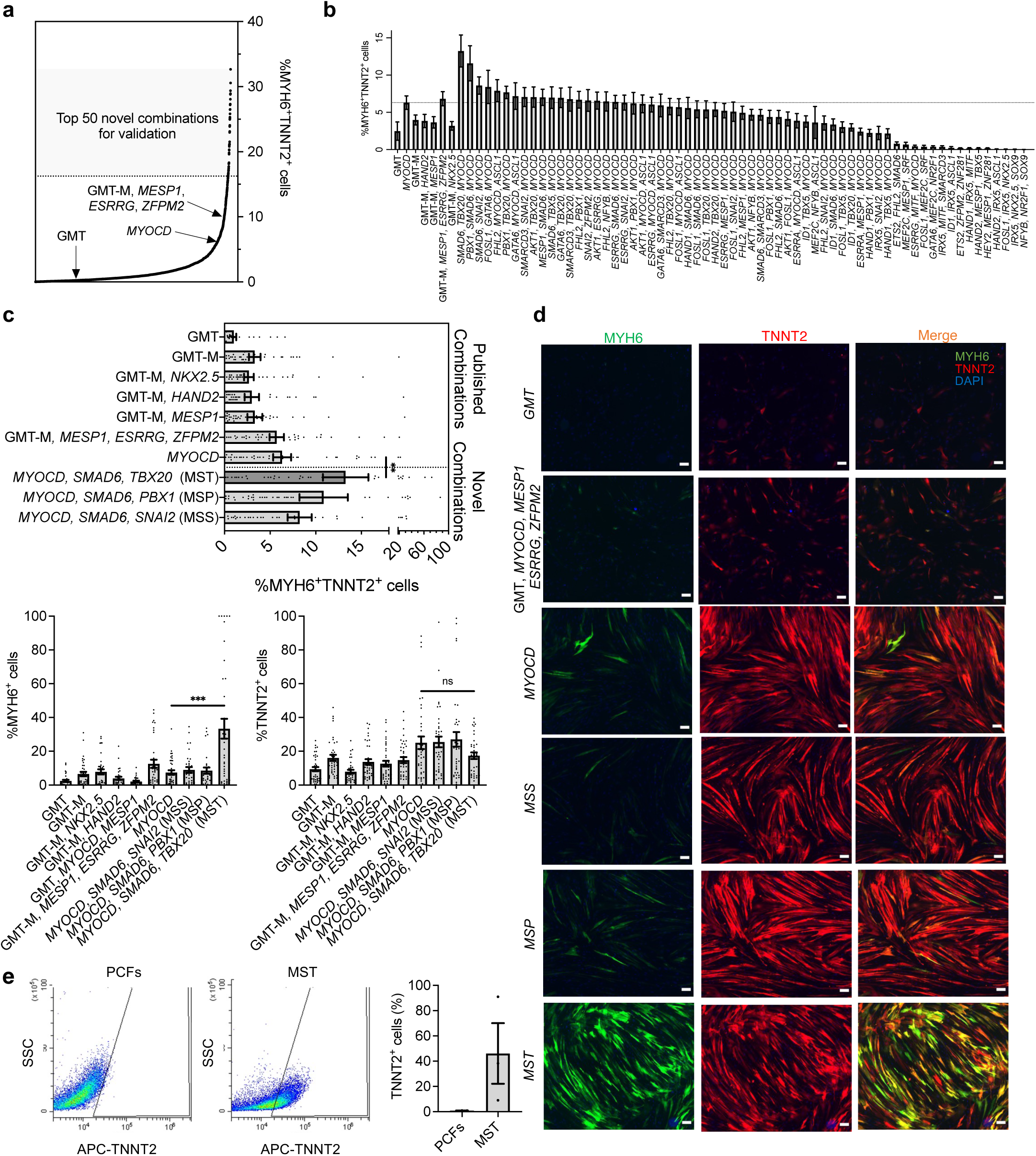
*MYOCD, SMAD6, TBX20* (MST) overexpression generates MYH6^+^TNNT2^+^ cells with higher efficiency compared to existing GMT-based transcription factor combinations. (**a**) Primary screen of transcription factor trios (*n* = 3 biological replicates). PCFs were transduced with rTTA and sMYH6-mNeon-TNNT2-mScarlet lentivirus and were subsequently transduced with e ach 3-factor combination from our lentiviral library. Cells were cultured in DMEM 10% FBS containing 0.25μg/mL doxycycline and fluorescence was quantified on reprogramming day 6. Distribution of means plotted. Top 50 factors, *MYOCD*, and GMT-M, *MESP1*, *ESRRG*, *ZFPM2* (best-performing published combination) indicated. (**b**) Validation of top 50 factors from lentiviral screen (*n* = 42 wells analyzed from 13 biological replicates) (**c**) Top 3 transcription factor trio combinations (*n* = 38 wells analyzed per condition from 13 biological replicates): %MYH6^+^TNNT2^+^ cells (top) %MYH6^+^ cells (bottom left) and %TNNT2^+^ cells (bottom right), measured by pFU-GW-sMYH6-mNeon-TNNT2-mScarlet fluorescence on reprogramming day 6. *P* values were calculated by paired t-test. (**d**) Images comparing *MYH6* and *TNNT2* reporter expression in novel combinations MST, MST, and MSS with *MYOCD* and published factor cocktails. Scale bar = 100 μM. Images acquired using high content image cytometry at reprogramming day 6 (KIC, Vala Sciences) (**e**) Flow cytometry analysis of TNNT2 expression in MST-transduced cells; reprogramming day 15. Representative data of untransduced and MST-transduced samples (left), quantification (right) (*n* = 3 biological replicates). ns *P* > 0.05, ** *P* ≤ 0.01, *** *P* ≤ 0.001. Error bars: mean ± SEM.

### Optimization of Reprogramming Media and Small Molecule Screen

Culture conditions have been reported to influence reprogramming efficiency and reprogrammed cell function based on cardiac gene expression. We optimized our reprogramming media to provide a more defined serum-free environment in which to assess the effect of different small molecules on cardiac reprogramming. We began by supplementing DMEM with 5 mg/mL bovine serum albumin (BSA), 200 µg/mL ascorbic acid, 10 µg/mL insulin, and 7 µg/L transferrin (D-BAITS). To further optimize this media formulation for direct reprogramming, we performed a dose optimization for each component. Our findings suggest that using 5 mg/mL BSA, 40 µg/mL ascorbic acid, 20 µg/mL insulin, 5 µg/mL transferrin, and 7 µg/L sodium selenite starting at reprogramming day 0 leads to an increased reprogramming efficiency in MST-transduced cells (**Figure 3A**). However, compared to DMEM+10% FBS, the optimized D-BAITS media decreased MST reprogramming efficiency measured by reporter activity by 15%, resulting in an average of 20.2% ± 5.6% MYH6^+^TNNT2^+^ reprogrammed cells, compared to 35.8% ± 12.2% in the presence of FBS (**Figure 3B**). To assess the endogenous expression of cardiac genes in D-BAITS media compared to FBS-supplemented media, we performed RT-qPCR at reprogramming day 6 on MST transduced cells with no reporter overexpression. Our data demonstrate that genes related to cardiac structure (*MYH6*, *TNNT2*, *MYL7*) in both conditions were upregulated. However, MST-induced cells derived in D-BAITS media showed significantly higher levels of endogenous sodium and calcium channel-related genes (*SCN5A*, *ATPA2*) compared to those derived cells in DMEM+10% FBS *(P* = 0.0004), suggesting improved cardiac function. Notably, *RYR2,* which encodes the ryanodine receptor essential for cardiomyocyte calcium handling was not upregulated in either condition, suggesting incomplete reprogramming (**Figure 3C**).

**Figure 3.**
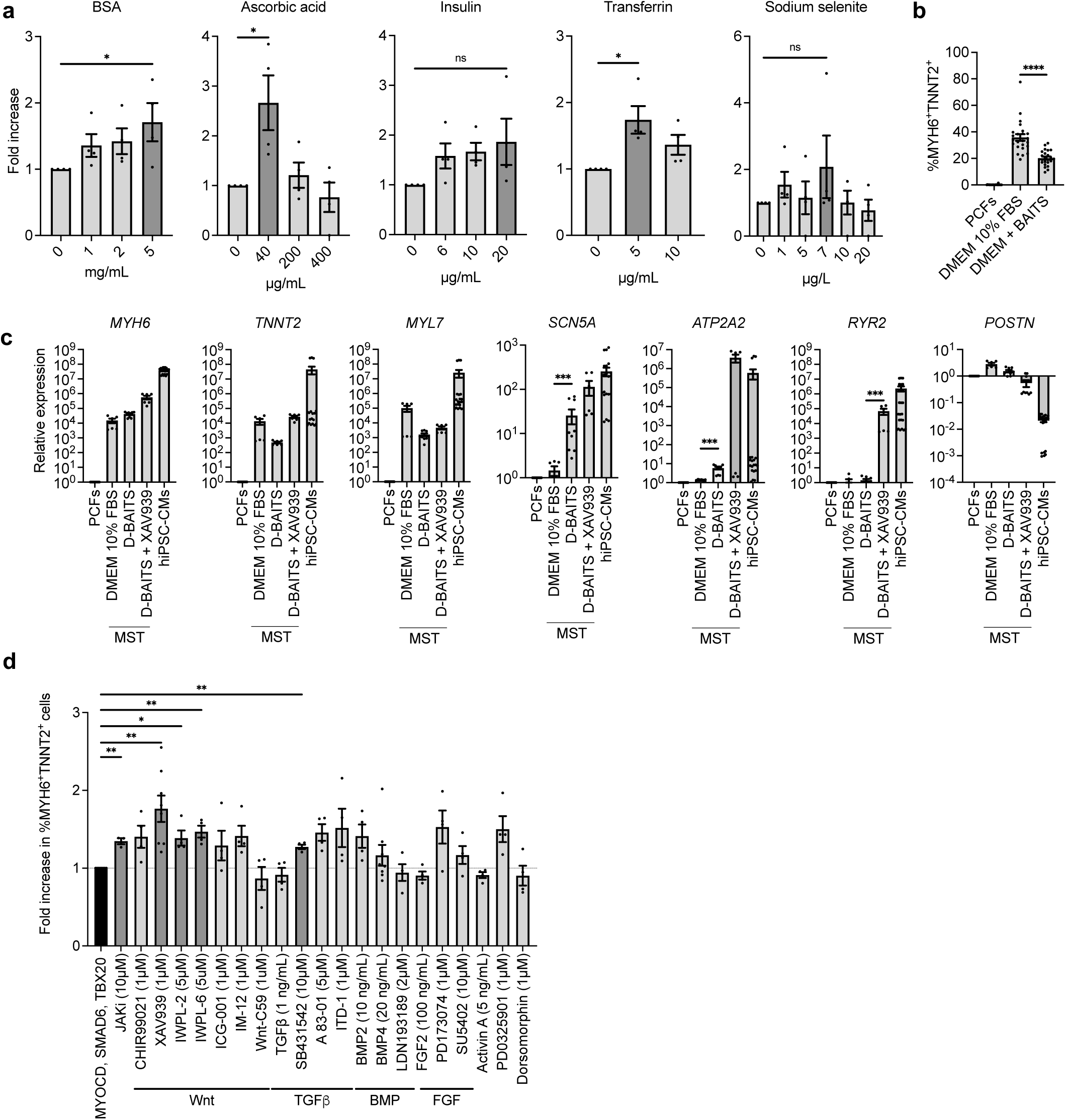
Wnt Inhibition using XAV939 enhances reprogramming efficiency of MST in defined media. **(a)** Reprogramming media component dose optimization. Fold increase in %MYH6^+^TNNT2^+^ cells, measured by pFU-GW-sMYH6-mNeon-TNNT2-mScarlet fluorescence on reprogramming day 6. Cells were transduced with MST and cultured in DMEM media containing BSA, ascorbic acid, insulin, transferrin, and sodium selenite starting 24h post-transduction. (*n* = 83 wells analyzed for each condition from 3-4 biological replicates; mean of biological replicates plotted). *P* values were calculated by t-test. **(b)** Percent mNeon (MYH6) mScarlet (TNNT2) double positive cells following culture in either DMEM+10% FBS or D-BAITS media following transduction with MST. Flourescence quantified on reprogramming day 6. (*n* = 24 technical replicates). *P* values were calculated by t-test, **(c)** RT-qPCR analysis for cardiac and fibroblast gene expression (*n* = 2-3 biological replicates), reprogramming day 6. *P* values were calculated by Mann-Whitney test. **(d)** Small molecule screen. Fold increase in percent MYH6^+^TNNT2^+^ cells on reprogramming day 6. Cells were transduced with MST and cultured in DMEM media containing 5 mg/mL BSA, 40 μg/mL ascorbic acid, 20 μg/mL insulin, 5 μg/mL transferrin, and 7 μg/L sodium selenite. (*n* = 40 wells analyzed for each condition from 3-4 biological replicates; mean of biological replicates plotted). *P* values were calculated by ANOVA followed by Fisher’s LSD test. ns *P* > 0.05, * *P* ≤ 0.05, ** *P* ≤ 0.01, *** *P* ≤ 0.001, **** *P* ≤ 0.0001. Error bars: mean ± SEM.

We then performed a small molecule screen to identify pathways that can be modulated to further increase the yield of MYH6^+^TNNT2^+^ cells and improve cardiac function. Small molecules and growth factors were added to cells 3 days after MST transduction, and reporter fluorescence was quantified 6 days after transcription factor induction with doxycycline-containing media. We used PCFs from 4 patient samples to screen 21 modulators of major signaling pathways (Wnt, TGFβ, BMP, and FGF) at a range of concentrations (**Figure S7**). Our findings demonstrated a consistent fold increase in MYH6^+^TNNT2^+^ cells following Wnt pathway inhibition with XAV939, IWP-L2, or IWP-L6. Whereas no significant change was observed following Wnt activation with CHIR99021, nor with modulation of BMP or FGF pathways. A slight, but significant increase in MYH6^+^TNNT2^+^ cells following treatment with the TGFβ inhibitor, SB431542 and JAKi was observed (**Figure 3D**). However, because reprogramming efficiency was most consistently and highly increased by Wnt pathway inhibition, we further characterized cells derived using MST and D-BAITS media supplemented with XAV939 (1 µM). Our data demonstrate that the addition of XAV939 to MST-derived cells significantly increased their expression of endogenous *RYR2* compared to use of MST alone (*P* = 0.0004) (**Figure 3C**). Addition of XAV939 to MST + D-BAITS media also increased expression of all other cardiac genes analyzed, especially that of *ATPA2* and, to a lesser extent, *SCN5A.* Co-expression of endogenous α-Actinin and TNNT2 at reprogramming day 30 was also observed. While levels of protein expression varied in the reprogrammed population, we observed striation patterns in both high- and low-expressing α-Actinin^+^TNNT2^+^ cells (**Figure S8A**). MST reprogrammed cells also displayed elevated expression of sarcomere-related genes *MYH6, ACTC1, TNNT2, TPM1, MYBPC3,* and *TTN* compared to PCFs (**Figure S8B**). In addition to upregulation of cardiac genes, we noted downregulation of the fibroblast marker gene *POSTN* in these conditions (**Figure 3C**).

### Functional assessment of MST-derived hiCMs

To determine the degree of reprogramming induced by MST and to compare the effect of media and small molecules on the reprogrammed cells’ function, we assessed the cells for spontaneous calcium flux. Reprogrammed cells were incubated with Cal-520, a fluorogenic calcium-sensitive dye prior to imaging. Fluorescence was measured for 60 seconds using high-content kinetic imaging at reprogramming day 7, 15, 22, and 30. Although calcium transients were detectable as early as differentiation day 7, they were more pronounced by day 22 (**Video S1**). At that time point, calcium transients were extremely rare (<0.1%) and slow (∼40 s) in cells cultured using DMEM+10% FBS (**Figure 4A-4B**) but were more commonly seen in cells cultured using D-BAITS media, with an average calcium transient duration 90% (CTD_90_) of 22.4s ± 1.7 s (**Figure 4A-4C**). Although the addition of XAV939 to MST-derived cells in D-BAITS media lowered the total number of cells cycling calcium by 44.7% compared to D-BAITS alone, their flux occurred more rapidly, with a CTD_90_ of 19.4 s ± 2.3 s compared to 22.4 s ± 1.7 s without XAV939 (**Figure 4A** **ii, 4B-C**). Previous studies using serum-free media and GMT in mouse suggest that FGF signaling improves functionality of directly reprogrammed cells by resulting in a 100-fold increase in beating cells^41^. In our system, addition of FGF2 (100 ng/mL) to D-BAITS resulted in a decrease in CTD_90_ time (14.2 s ± 0.9 s compared to 22.4s ± 1.7s without FGF2) and a 38.3% increase in the total number of cycling cells (**Figure 4B**). Simultaneous treatment with XAV939 and FGF2 did not significantly change the CTD_90_ (12.3 s ± 0.75s) compared to FGF2 alone; however, reproducibility, based on a lower SEM (0.74 s compared to 0.90 s), was increased without a decrease in the total number of cycling cells.

**Figure 4.**
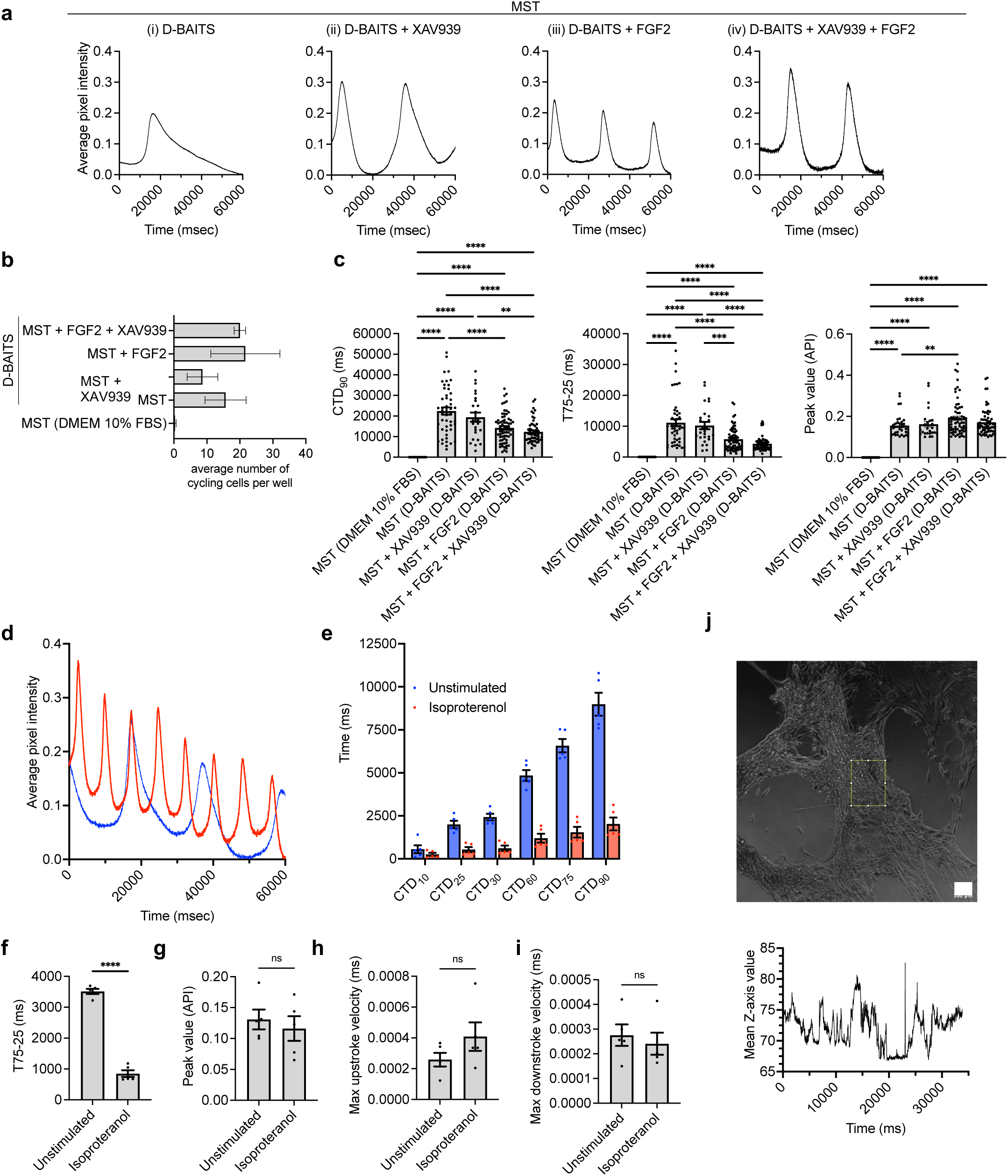
FGF2 enhances calcium cycling in MST-derived cells and produces contractile cells that respond to pharmacological stimulation. (**a**) Spontaneous Ca^2+^ oscillations in reprogrammed cells on reprogramming day 22; representative traces: (i) DMEM + BAITS, (ii) DMEM + BAITS + XAV939, (iii) DMEM + BAITS + FGF2, (iv) DMEM + BAITS + XAV939 + FGF2 (**b**) Aggregate number of cycling cells per well of a 384-well plate detected by analysis software (**c**) CTD90 (left), T75-25 (center), peak value (right) per condition (*n* = 3 wells per condition). *P* values calculated using one-way ANOVA with subsequent pairwise testing via t-test with Tukey’s adjustment for multiple comparisons. (**d**) representative traces of unstimulated (blue) and isoproterenol stimulated (red) cells. Effects of 2 μM isoproterenol stimulation on (**e**) calcium-transient duration of reprogrammed cells (f) T75-25 (**g**) peak value (**h**) maximum upstroke velocity (**i**) maximum downstroke velocity (*n* = 5 cells per condition). *P* values calculated using t-test. (**j**) Spontaneous contraction in MST-derived cells; reprogramming day 25. Still from video (top), quantification of movement along Z-axis within ROI (yellow) (bottom). ns *P* > 0.05, ** *P* ≤ 0.01, *** *P* ≤ 0.001, **** *P* ≤ 0.0001. Error bars: mean ± SEM.

To investigate whether MST-derived cells respond to pharmacological stimulation, we analyzed calcium flux in response to the β-adrenergic agonist isoproterenol (2 μM), as previously described^42^. Treatment with isoproterenol resulted in faster calcium cycling with a CTD_90_ of 2.0 s ± 0.4s compared to 8.9 s ± 0.7s for untreated cells, as would be expected with cardiomyocytes (**Figure 4D-E**). However, peak amplitude as well as maximum upstroke and downstroke velocity were unchanged, suggesting that further optimization of Ca2^+^ signaling machinery is necessary to produce cells that fully emulate cardiac function. (**Figure 4F-I**). We monitored spontaneous cell contraction and observed cell movement as early as day 12 of differentiation, with contraction becoming more evident by day 29 (**Figure 4J, Video S2**).

To determine the extent of reprogramming in MST-derived cells, we performed RNA sequencing and compared the transcriptome of three samples of MST-reprogrammed hiCMs to that of matched cardiac fibroblasts using hiPSC-CMs as a control. PCA of gene expression data showed distinct clustering of individuals into the three sample groups (PCFs, MST-reprogrammed hiCMs, and hiPSC-CMs) based on PC1 (67% of variance) and PC2 (24% of variance) (**Figure 5A**). The top 10 upregulated genes contributing to PC1, based on absolute value of the loadings, are all mitochondrial related (including *MT-CO1*, *MT-RNR2*, *MT-ND4*, etc.). These are followed by cardiac *ACTC1*, *MYL7*, and *MYH6*. For PC2, the top upregulated genes are smooth muscle related *ACTA2*, *MYH11*, *FLNA*, and calcium related *MYLK.* As expected, top downregulated genes for both PC1 and PC2 are fibroblast related (*FN1*, *COL1A1*, and *COL1A2*, and *EEF1A1)* (**File S1**). This independent clustering was further corroborated by hierarchical cluster analysis (**Figure 5B**). Together, these data suggest that MST-reprogrammed hiCMs are in an intermediate state between cardiac fibroblasts and hiPSC-CMs. Nonetheless, gene-set enrichment analysis (GSEA) revealed an upregulation of cardiac gene sets in MST-derived cells compared to PCFs. Enriched gene sets include cardiac conduction, calcium regulation in cardiac cells, and cardiac muscle contraction (**Figure 5C**). Likewise, Ingenuity Pathway Analysis (IPA) identified several significant cardiac-related pathways in MST-derived cells, including cardiac hypertrophy signaling (*P* = 1.48e-10), dilated cardiomyopathy signaling (*P* = 1.12e-08), calcium signaling (*P* = 8.13e-06), cardiac β-adrenergic signaling (*P* = 4.57e-04), and factors promoting cardiogenesis in vertebrates (*P* = 0.02) (**File S2**).

**Figure 5.**
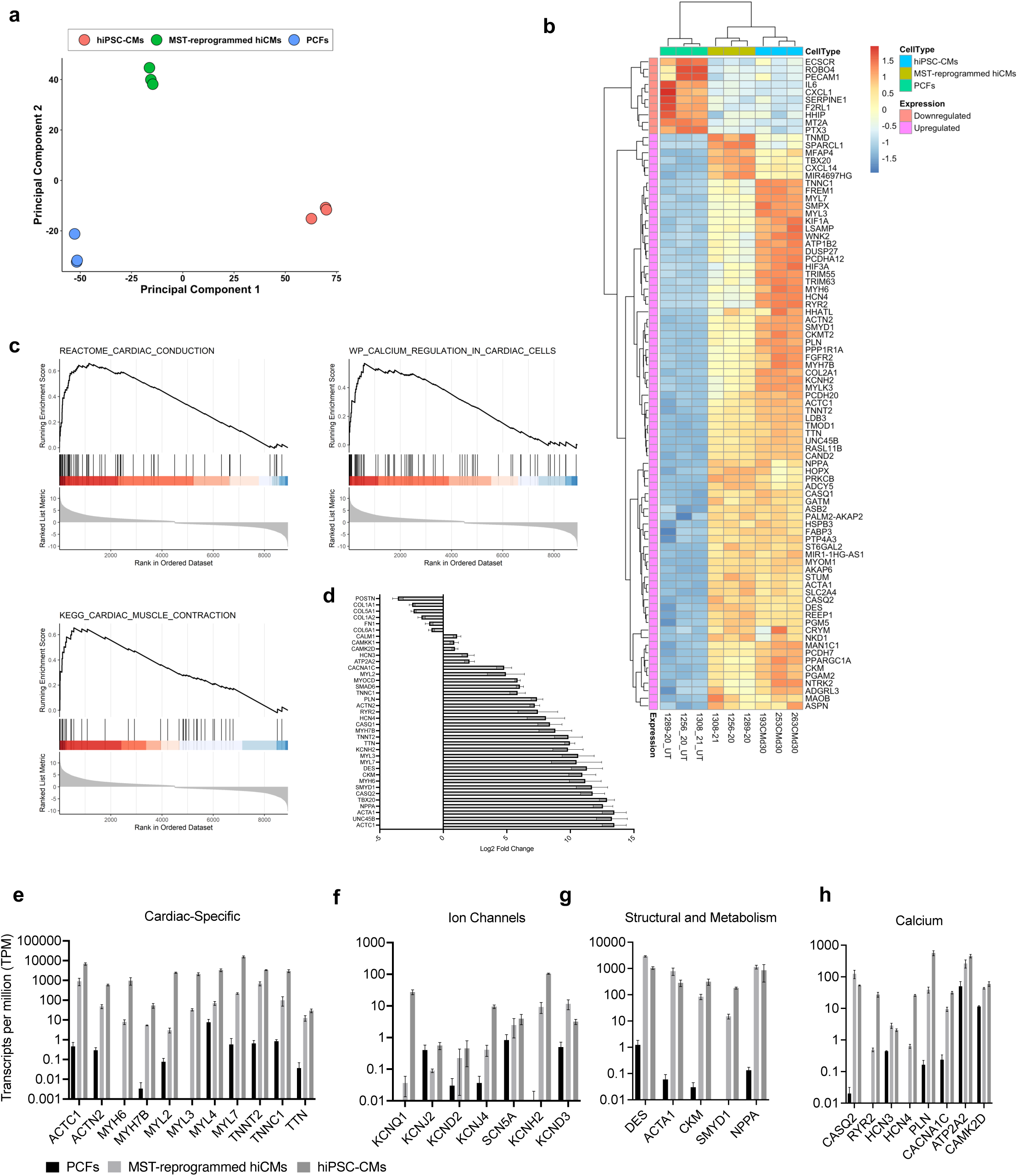
RNA-seq analysis reveals cardiac transcriptome changes in MST-reprogrammed cells. **(a)** Principal component analysis of MST-reprogrammed hiCMs, PCFs, and hiPSC-CMs (n =3 biological replicates per condition). **(b)** Hierarchical clustering of 86 variable genes significantly upregulated or downregulated in both hiPSC-CMs and MST-derived hiCMs (*n* = 3, P-adjusted value < 0.00001 and abs(log2foldchange) > 5). **(c)** Enrichment plots of the indicated gene sets: cardiac conduction (top left), calcium regulation in cardiac cells (top right), cardiac muscle contraction (bottom)**. (d)** Comparative expression of cardiac genes in MST-reprogrammed hiCMs vs. primary cardiac fibroblasts. Transcripts per million for expression of **(e)** cardiac-specific genes, **(f)** ion channels, **(g)** structural and metabolism-related genes **(h)** calcium-related genes in primary cardiac fibroblasts (PCFs) (black), MST-reprogrammed hiCMs (light gray), and hiPSC-CMs (dark gray). Error bars: mean ± SEM.

Analysis of the 9,293 differentially expressed genes (FDR < 0.05) in MST-derived cells over fibroblasts also suggests a global shift to a cardiac-like state (**Figure 5D, S9A**). As expected, MST-derived hiCMs expressed high levels of *MYOCD*, *SMAD6*, and *TBX20*, even higher than levels found in hiPSC-CMs (**Figure S9B**). Cardiac fibroblast markers *IL6*, *MMP1* (collagenase 1), *SERPINE1*, *POSTN*, *P4HB*, *COL1A1*, *COL1A2*, and *FN1* were all downregulated after MST reprogramming to levels similar to hiPSC-CMs (**Figure S9C**). Compared to cardiac fibroblasts, MST-derived hiCMs showed significant upregulation of cardiac-specific genes, including *ACTC1, ACTN2, MYH6, MYH7B, MYL2, MYL3, MYL4, MYL7, TNNT2, TNNC1,* and *TTN* (**Figure 5E**). We also noted expression of all cardiac ion channel genes: *KCNQ1, KCNJ2, KCND2, KCNJ4, SCN5A, KCNH2,* and *KCND3* (**Figure 5F**). Upregulated expression of genes encoding cardiac structural proteins (*DES, ACTA1, UNC45B*) and proteins involved in cardiac cell metabolism and signaling (*CKM SMYD1, NPPA*) were also observed in MST-derived hiCMs, (**Figure 5G**). Finally, we noted increased expression of several cardiac calcium-handling related genes (*CASQ2, RYR2, HCN3, HCN4, PLN, CACNA1C, ATP2A2, CAMK2D*) in MST-derived hiCMs relative to PCFs (**Figure 5H**).

While this data demonstrates a transcriptomic shift towards a cardiac state, the overall expression of cardiac-related genes in MST-reprogrammed hiCMs remains lower than that of hiPSC-CMs, suggesting that further optimization, perhaps by inclusion of growth factors, additional reprogramming factors, or small molecules is necessary for complete reprogramming.

## DISCUSSION

We have optimized a lentiviral transduction and cardiac reporter platform to efficiently screen combinations of transcription factors for their ability to produce cells with active cardiac MYH6 and TNNT2 promoters and to assay for calcium flux. Using this platform, we identified a novel combination of transcription factors: *MYOCD, SMAD6, TBX20* (MST) that reprograms human PCFs into cardiomyocyte-like cells based on gene expression and sarcomere structure. MST, in conjunction with FGF2, XAV939, and our dose-optimized media containing BSA, ascorbic acid, insulin, transferrin, and sodium selenite, allows for the derivation of cells with functional cardiac characteristics. We, like others, have found that reprogramming efficiency and calcium flux are enhanced by addition of the Wnt pathway inhibitor, XAV939. In our system, the number of calcium cycling cells is further increased with the addition of FGF2. Together, these conditions result in the reprogramming of human PCFs to cardiac-like cells that are responsive to pharmacological stimulation and spontaneously contract in culture.

We have used MST to successfully reprogram 24 distinct primary samples from patients (aged 0.01 - 21 years) undergoing cardiac biopsy for diverse diagnoses, as well as one healthy tissue (**Figure S2**). To our knowledge, this is the first study to demonstrate direct cardiac reprogramming using cells from numerous patients and diverse cardiac diagnoses. Expectedly, we have found that reprogramming efficiency is variable from patient to patient, with some lines not converting, and have included all such instances in our data. While genetic background may play a role in reprogramming efficiency, we speculate that the inherent diversity in starting fibroblast population and activation state may also contribute to differences in reprogramming outcomes. Further studies will be necessary to determine whether MST-reprogrammed cells retain disease-associated phenotypes and whether reprogramming efficiency differs when using aging tissues. Likewise, we have found that MST-derived cells express both atrial and ventricular-associated genes (**Figure S9D**). Future optimizations focusing on subtype specification may be beneficial for disease modeling. Although hiCM-derived cells express several genes associated with a mature electrophysiological phenotype, including *KCNJ2*, *KCNQ1*, *SCN5A*, *CACNA1C*, *RYR2*, and *ATP2A2*, expression of genes associated with myofibril isoform switching (*MYH7*, *MYL2*, *TNNI3*) is low relative to fetal-associated *MYH6*, *MYL7*, and *TNNI1 (***Figure S9E**); highlighting an area for further improvement of this reprogramming protocol.

*MYOCD* and T-box factors have been utilized in existing cardiac direct reprogramming cocktails. TBX20 is known to play a role in cardiac development through regulation of GATA4 and NKX2-5 transcription^43^. It is also implicated in maintaining function of the cardiac conduction system^44^. While this manuscript was under review, Tang et. al independently identified TBX20 as a cardiac reprogramming factor^45^. In their study, addition of TBX20 to GMT + miR133 resulted in cells with more fully activated cardiac conduction systems. The reprogrammed cells exhibited increased mitochondrial function and spontaneous calcium flux after 1 month; however, contraction was observed only in co-culture with hiPSC-CMs, limiting the therapeutic applications of the approach.

While overexpression of *MYOCD* alone in PCFs led to activation of our cardiac reporter and to some cardiac gene induction, no spontaneous contraction nor calcium flux was observed in the resulting cells. Partial activation of the cardiac gene network, including aMHC and cTnT, by overexpression of *MYOCD* alone in human foreskin fibroblasts has been previously reported^39^. However, as in our study, the authors conclude that additional factors are necessary to derive cells that are fully committed to the cardiac fate. Similarly, in mouse fibroblasts, addition of *MYOCD* to GMT contributes to improved cardiac gene expression but does not enhance cardiac function^40^.

Further studies will be necessary to determine the mechanism of MST-based direct reprogramming; however, we note that the factors *MYOCD* and *SMAD6* were present in each of the top 3 combinations identified in our screen. This may suggest a shared reprogramming mechanism driving activation of our aMHC-TNNT2 transgenic reporter. SMAD6 is known to inhibit TGFβ signaling through inhibition of receptor-associated SMADs (SMAD2/3) and TGFβ inhibition using SB431542 has been shown to improve the efficiency of GMT-based reprogramming^9^. Therefore, we speculate that SMAD6 allows for direct reprogramming through a similar mechanism. In our system, the addition of TGFβ1 did not affect MST-reprogramming efficiency; however, further TGFβ inhibition using SB431542, A83-01, or ITD-1 did result in a marginal increase in MYH6^+^TNNT2^+^ cells.

Directly reprogrammed cardiomyocytes have the potential to advance precision medicine by providing an expedient source of patient-matched cells that can be used *in vitro* for drug screening or *in vivo* as therapy for heart failure. This study presents MST as a novel transcription factor combination that is suited for direct reprogramming of human cardiac fibroblasts. Further studies can determine whether efficiency of this system can be improved by using a polycistronic construct and by optimizing the timing of small molecule delivery. Due to the observed functional characteristics of MST-derived cells, future investigational and therapeutic approaches may benefit from the use of this combination.

## Supporting information

Video_S1_Calcium

Video_S2_Contraction

File_S1_PCA_Loading

File_S2_IPA

## ACKNOWLEDGEMENTS

The authors would like to thank all patient donors for their contributions to this work. Graphical abstract and Figure 1a were created with BioRender.com.

## ACCESSION CODES

Gene Expression Omnibus: RNA-seq data have been deposited with accession code GSE218091.

## SUPPLEMENTAL MATERIAL

Tables S1-S7

Figures S1-S9

Files S1-S2

Videos S1-S2

References 1-45

## ABBREVIATIONS

MST: MYOCD, SMAD6, TBX20
MI: myocardial infarction
GMT: *GATA4*, *MEF2C*, *TBX5*
GMT-M: GATA4, MEF2C, TBX5, MYOCD
iCMs: cardiomyocyte-like cells, induced cardiomyocytes
hiCMs: human cardiomyocyte-like cells, human induced cardiomyocytes
GMTM-H: GATA4, MEF2C, TBX5, MYOCD, HAND2
PCFs: primary cardiac fibroblasts
hiPSC-CMs: human induced pluripotent stem cell-derived cardiomyocytes
BSA: bovine serum albumin
MOI: multiplicity of infection
MSP: MYOCD, SMAD6, PBX1
MSS: MYOCD, SMAD6, SNAI2
D-BAITS: DMEM, bovine serum albumin, ascorbic acid, insulin, transferrin, sodium selenite
CTD_90_: calcium transient duration 90%

---

**Figure S1.**
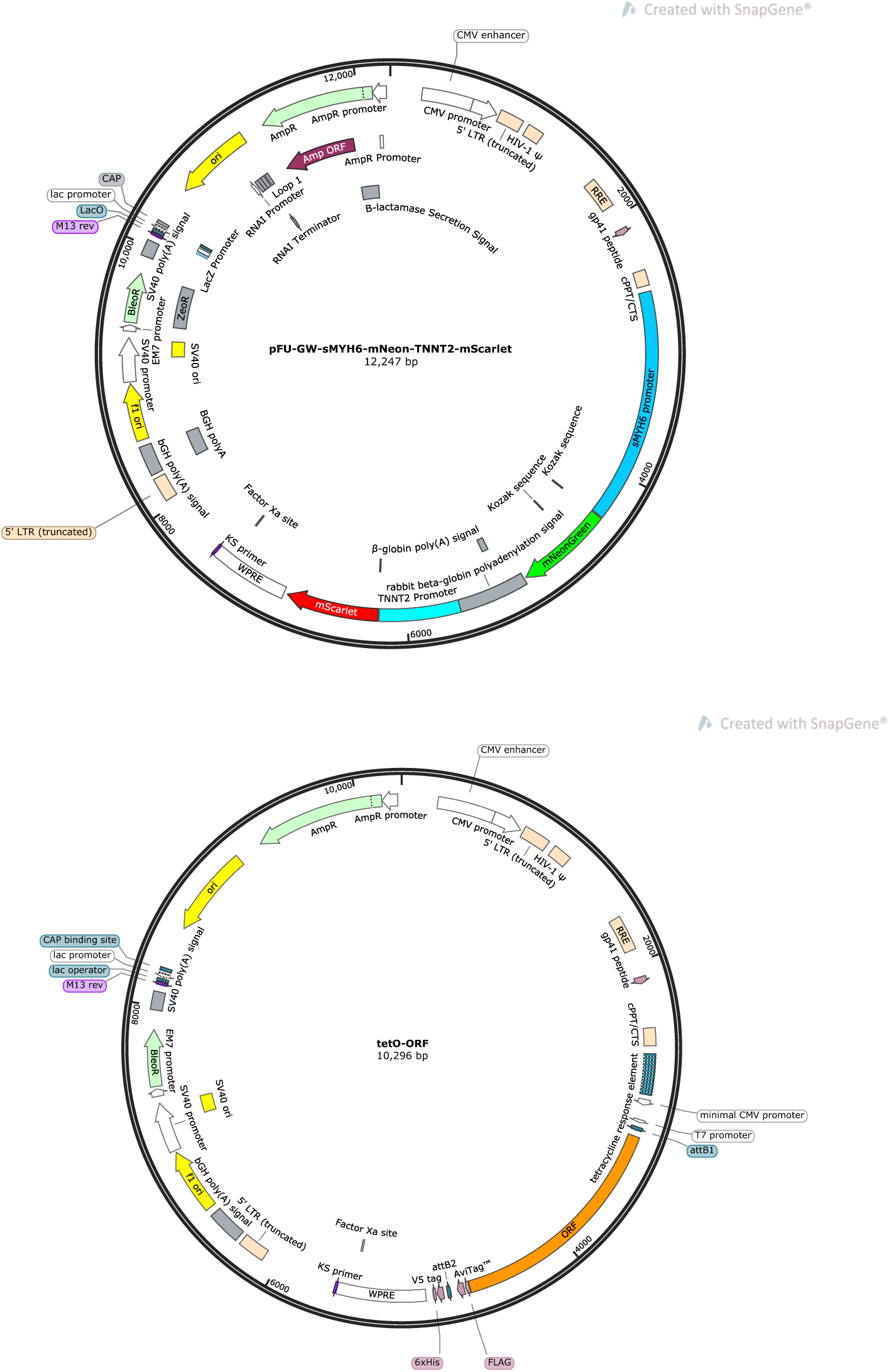
Plasmid maps. (A) Dual-fluorescent reporter plasmid pFU-GW-sMYH6-mNeon-TNNT2-mScarlet (Addgene #170712). (B) cDNA expression plasmid tetO-ORF.

**Figure S2.**
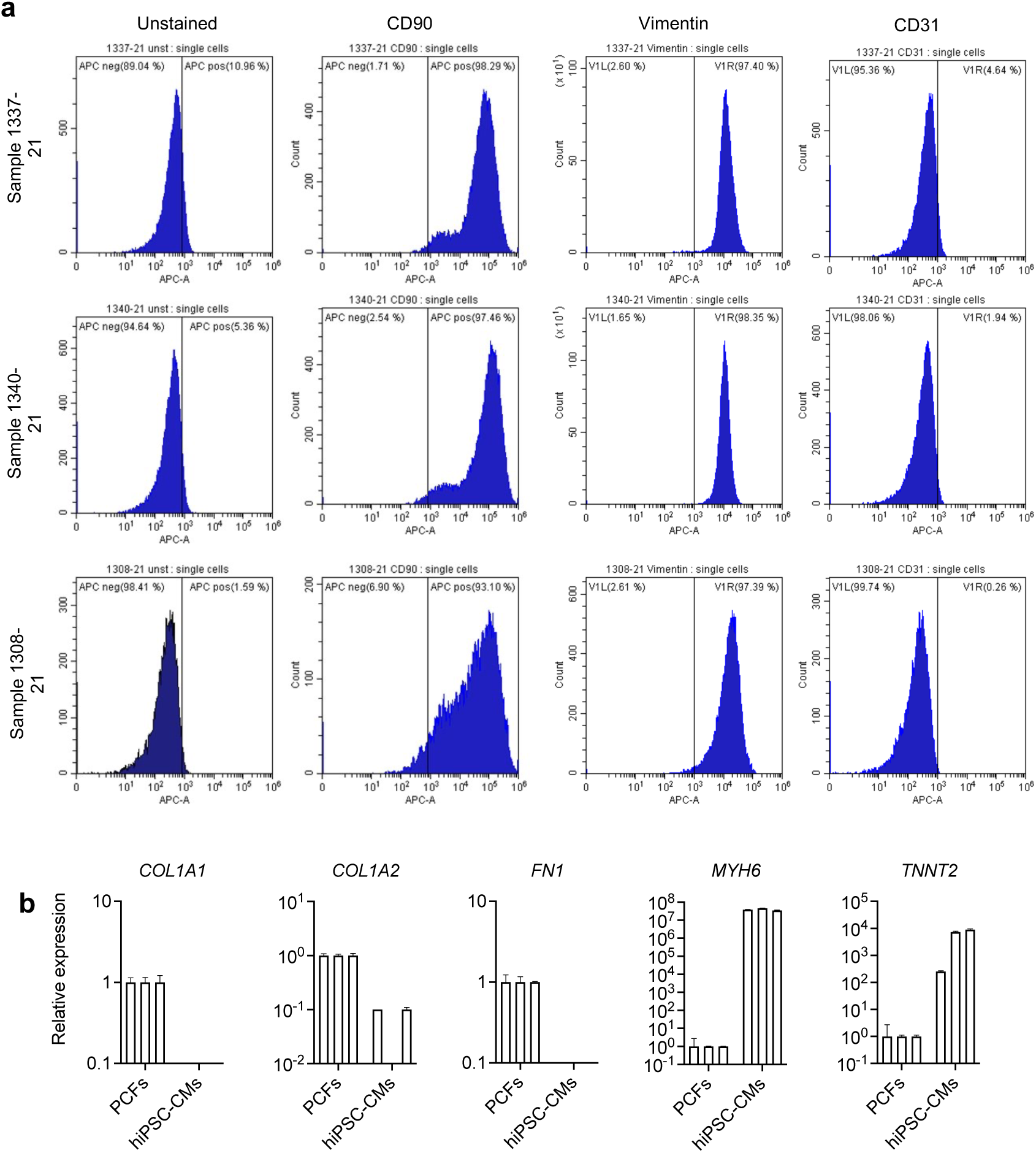
Outgrowth PCFs cells express fibroblast markers. (a) Cells stain positive for fibroblast markers CD90 and vimentin and negative for endothelial cell marker CD31 by flow cytometry (*n* = 3 samples). (b) Cells express fibroblast markers COL1A1, COL1A2, and FN1 and do not express cardiac markers MYH6 nor TNNT2 by qRT-PCR (*n* = 3 samples) .

**Figure S3.**
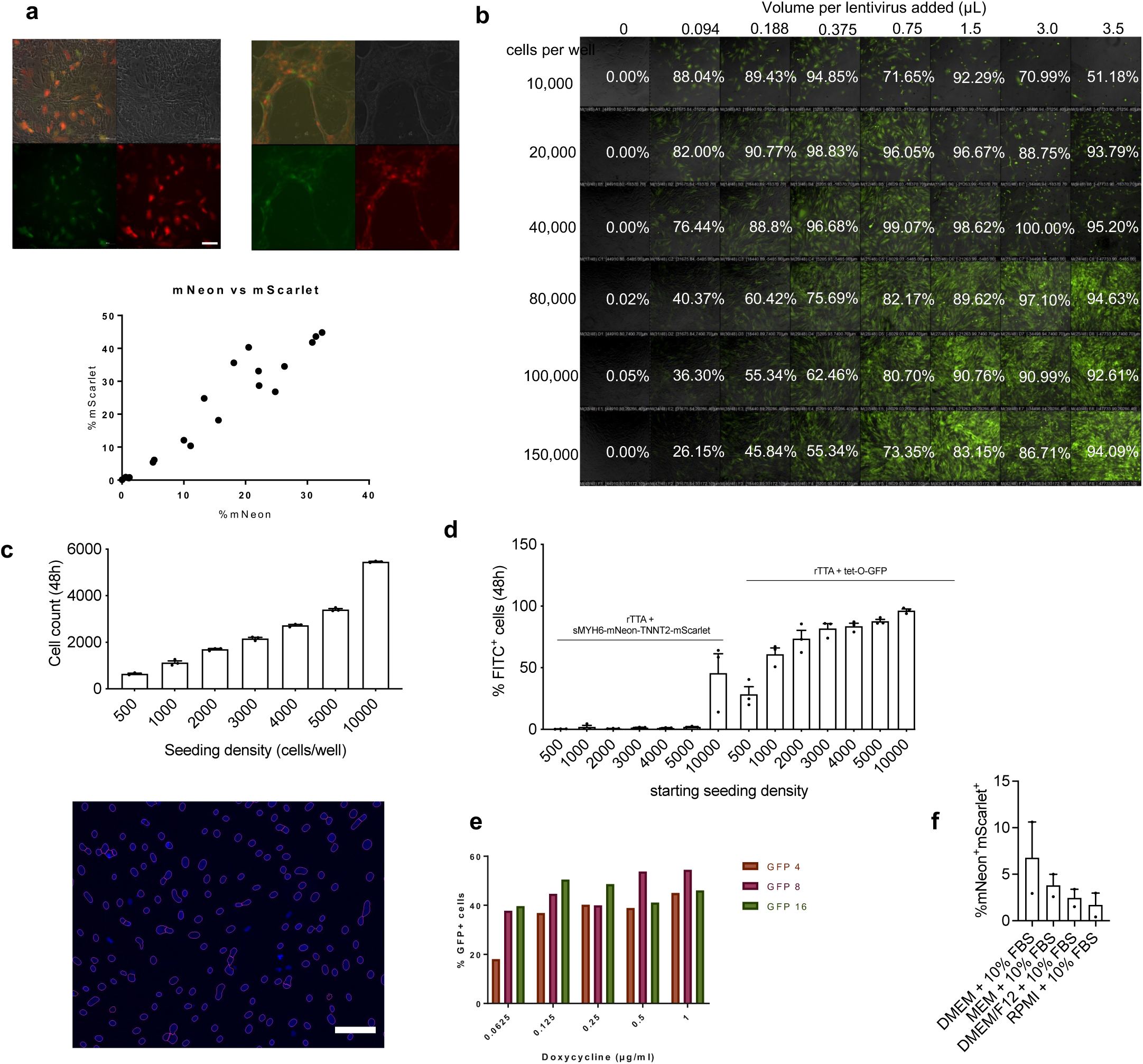
Lentiviral Transduction Optimization. **(a)** Validation of pFU-GW-sMYH6-mNeon-TNNT2-mScarlet reporter. hiPSC-CMs (d32) transduced with dual reporter (left) exhibited mNeon and mScarlet fluorescence 72 h post-transduction, and transduced hiPSCs differentiated into hiCMs exhibited mNeon and mScarlet fluorescence starting at differentiation d17 (right). mNeon and mScarlet fluorescence linearly correlate (bottom) (Scale bar = 100 μM) **(b)** Lentiviral transduction efficiency of primary cardiac fibroblasts with pFU-tet-O-GFP and rTTA. Cells were passaged to a Matrigel-coated 96-well plate at various seeding densities and transduced using Transdux Max with a range of lentiviral volumes dispensed by liquid handler. Cells were imaged and analyzed by flow cytometry at 72 h post-transduction. Percent positive GFP cells in white text. For direct reprogramming, MOI was scaled accordingly and 50nL of concentrated lentivirus was used to infect 3,000 cells per well in a 384-well plate, resulting in a transduction efficiency of >90%. **(c)** Validation of VALA gating algorithm. Increased seeding density corresponds with higher cell count (left). Representative gating image (right). Cells for transduction were seeded at 3,000 cells per well in a 384-well plate. Average pixel intensity within the nuclear gate is used to quantify percent positive cells using our fluorescent reporter (Scale bar = 100 μM). **(d)** Seeding density optimization. Cells were plated in 384-well plates and transduced with tet-O-GFP to assess transduction efficiency and with sMYH6-mNeon alone as a negative control. Percent positive cells were quantified at 48h post-transduction. Seeding cells at 3,000 cells per well resulted in >90% transduction efficiency and no fluorescence in reporter-transduced cells. (n=3 biological replicates) **(e)** Doxycycline dose optimization. 0.125 μg/mL doxycycline is sufficient to induce GFP expression at 72 h post-transduction and no toxicity is observed at concentrations up to 1 μg/mL. We used 0.25 μg/mL doxycycline for direct reprogramming. **(f)** Basal media selection. Cells were transduced with pFU-GW-sMYH6-mNeon-TNNT2-mScarlet, rTTA, and tet-O-MYOCD. Fluorescence was quantified on reprogramming day 6 (*n* = 2 biological replicates). DMEM+10% FBS was used going forward for direct reprogramming.

**Figure S4.**
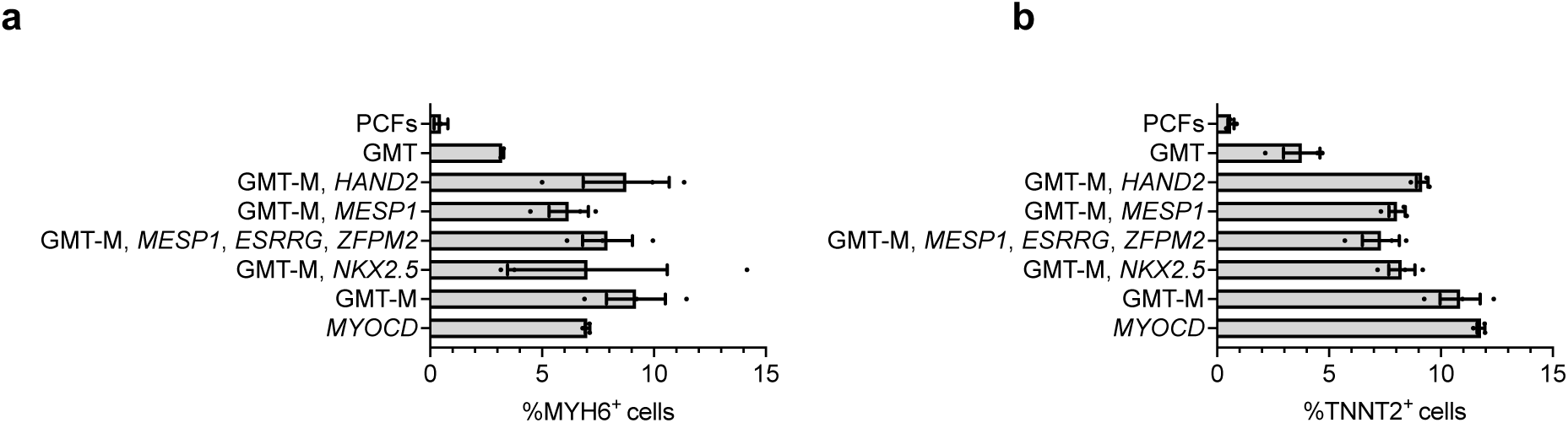
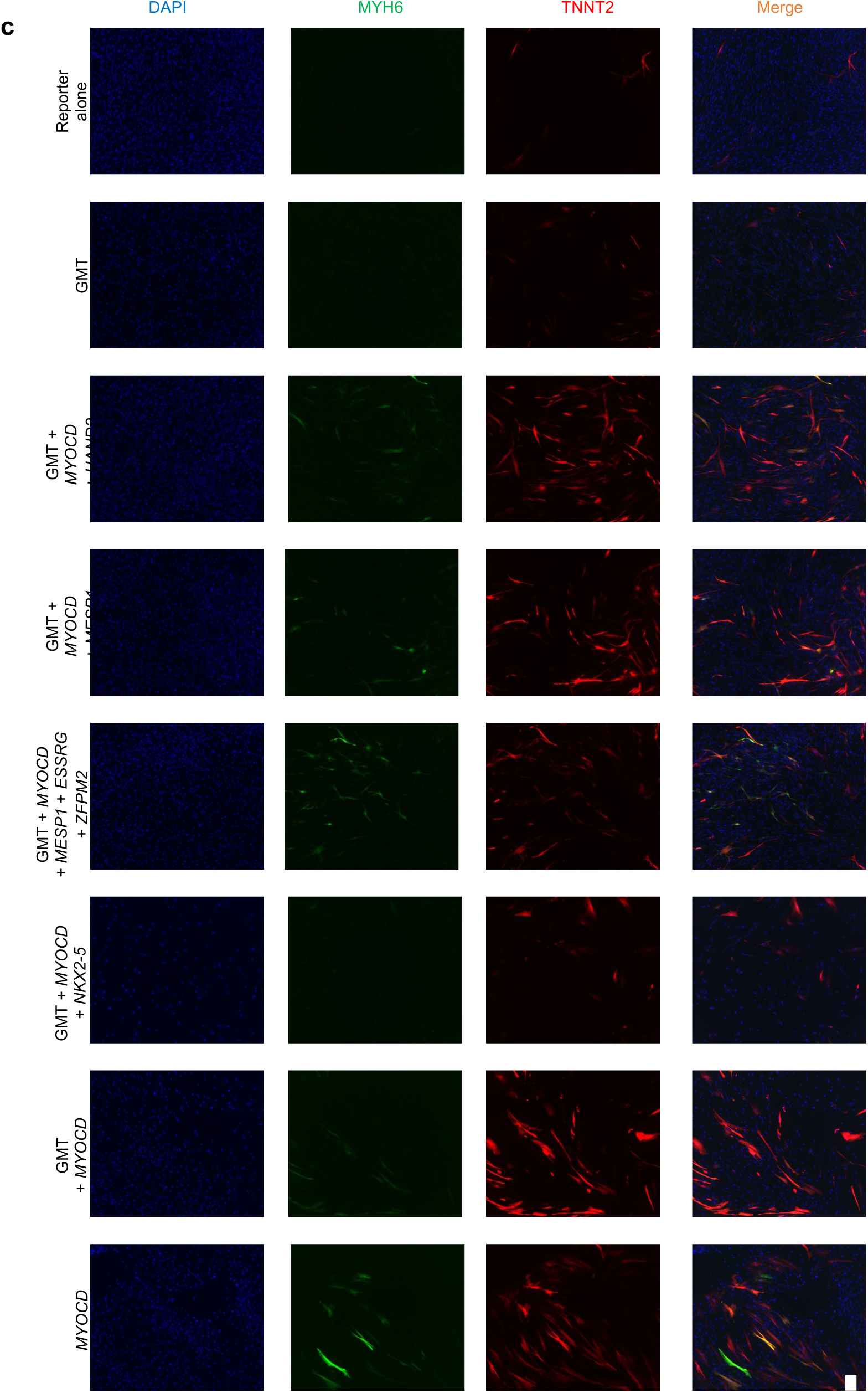
Published transcription factor combinations activate MYH6 and TNNT2 promoters at low levels. Quantification of **(a)** MYH6^+^ and **(b)** TNNT2^+^ cells as measured by pFU-GW-sMYH6-mScarlet-TNNT2-mNeon reporter fluorescence on reprogramming day 6 (*n* = 3 biological replicates). All cells were subject to reporter overexpression. Quantification of **(c)** Representative images of primary cardiac fibroblasts transduced with sMYH6-mNeonGreen-TNNT2-mScarlet and reprogrammed using published factor combinations. Scale bar = 100 μM. Images acquired using high content image cytometry at reprogramming day 6 (KIC, VALA Sciences).

**Figure S5.**
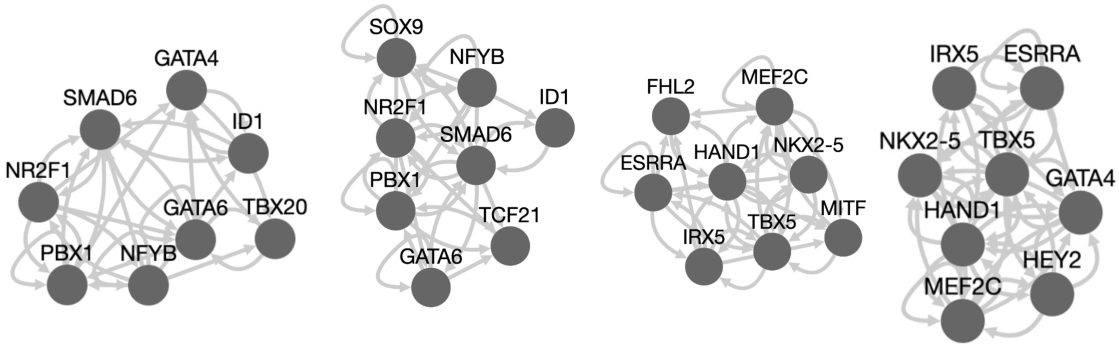
Reprogramming transcription factor networks predicted by Mogrify. Left to right: fibroblast to cardiac myocyte, cardiac fibroblast to cardiac myocyte, cardiac fibroblast to adult heart, and fibroblast to adult heart.

**Figure S6.**
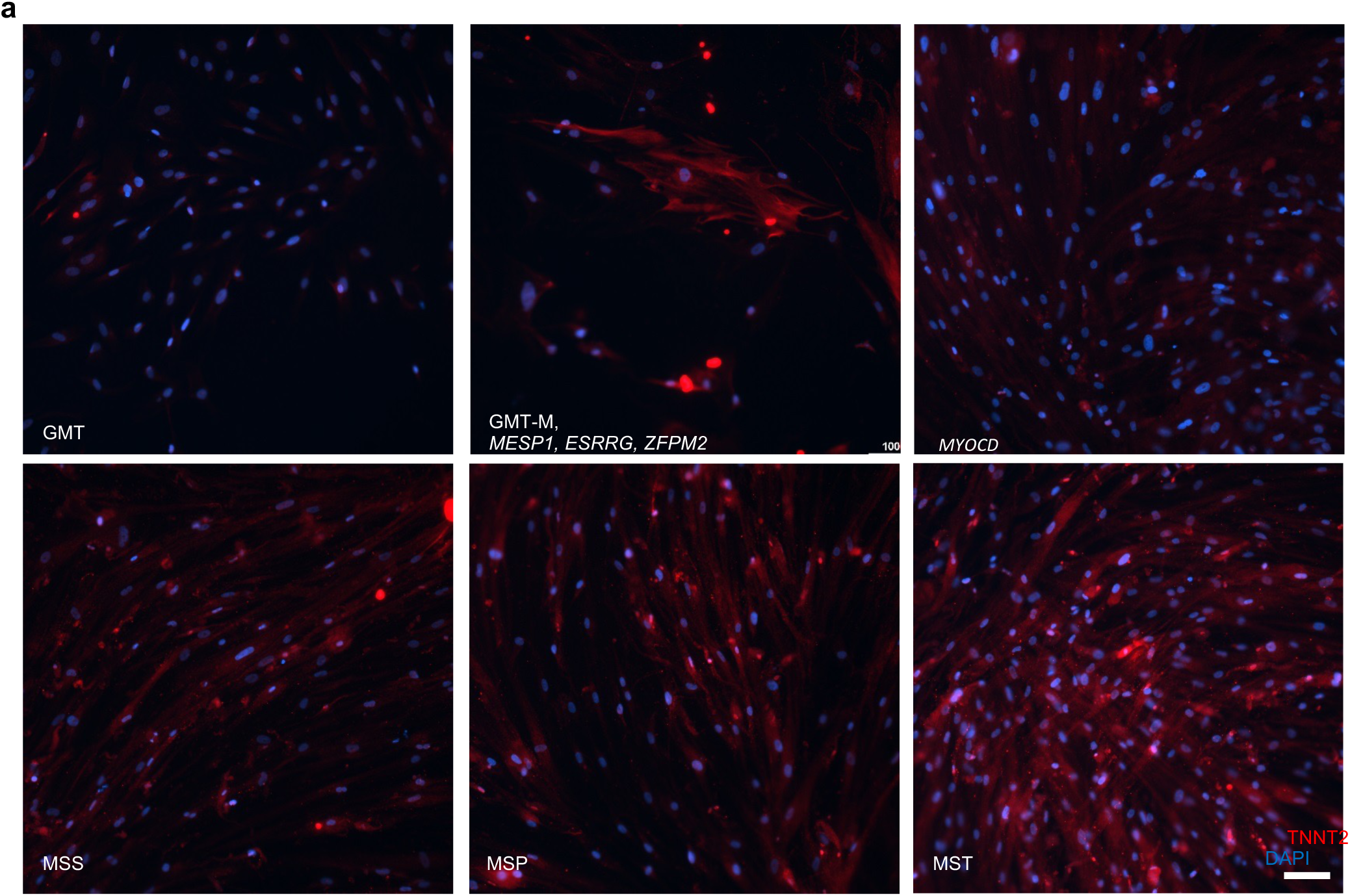
TNNT2 immunostaining of top 3 novel combinations; reprogramming day 60.

**Figure S7.**
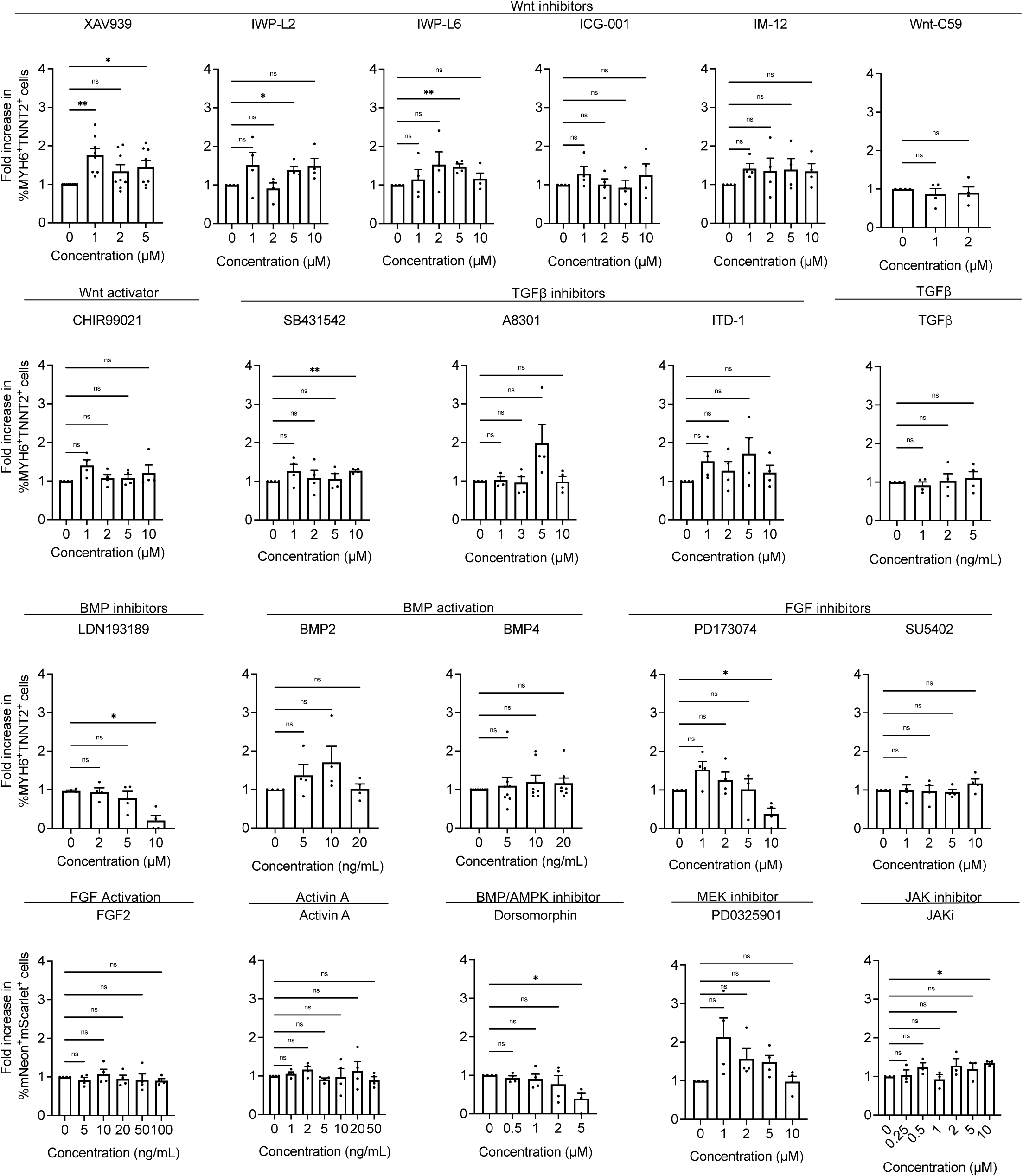
Small Molecule Screen. Cells were transduced with MST and exposed to small molecules starting on reprogramming day 3. Fluorescence was quantified on reprogramming day 6. (*n* = 40 wells analyzed for each condition from 3-4 biological replicates; mean of biological replicates plotted). *P* values were calculated by ANOVA followed by Fisher’s LSD test. ns *P* > 0.05, * *P* ≤ 0.05, ** *P* ≤ 0.01, *** *P* ≤ 0.001, **** *P* ≤ 0.0001.

**Figure S8.**
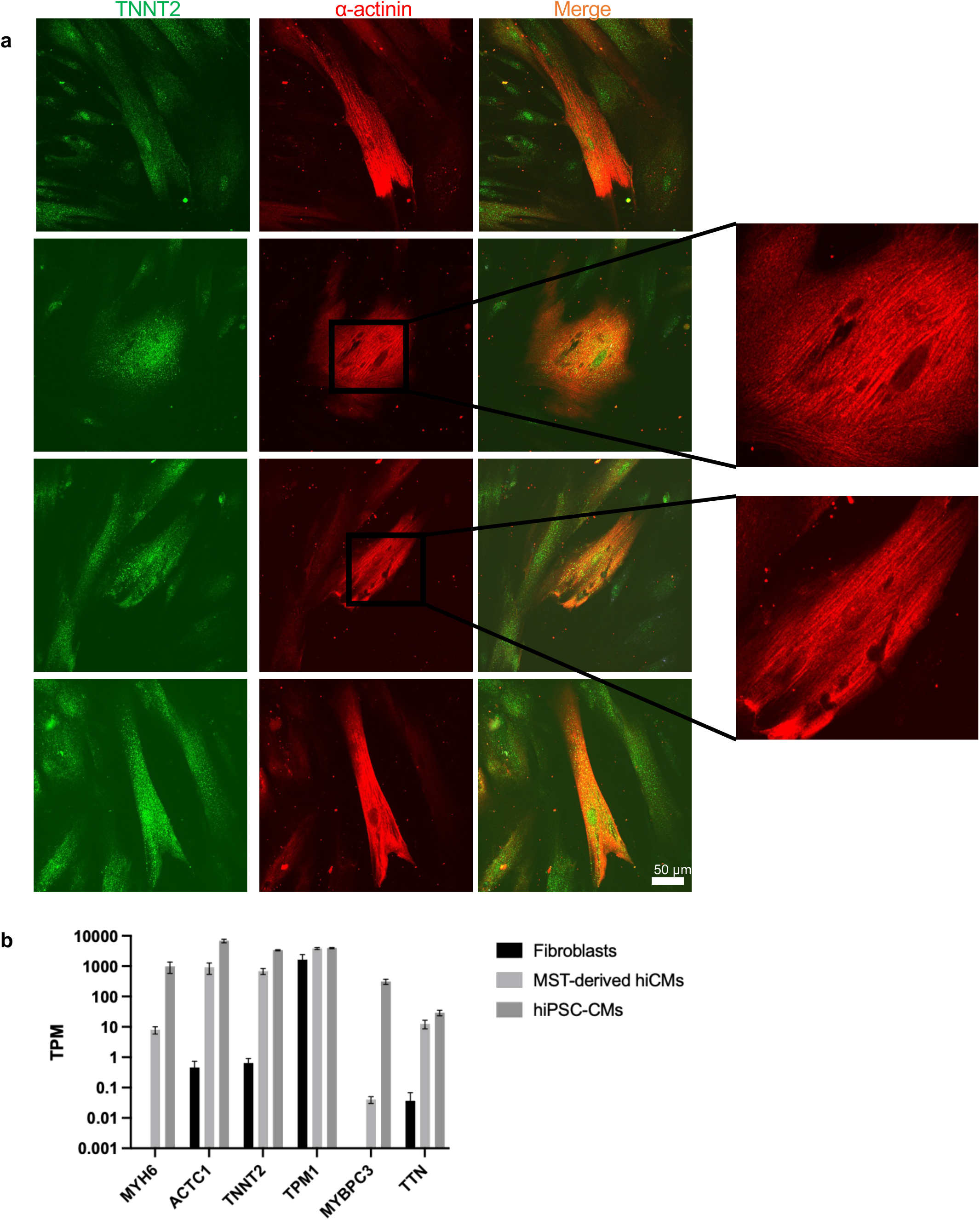
MST-derived cells express sarcomeric genes and proteins. **(a)** TNNT2 and α-actinin immunostaining on reprogramming day 30. D-BAITS (DMEM, BSA, Ascorbic acid, Insulin, Transferrin, Sodium selenite). MST (*MYOCD*, *SMAD6*, *TBX20*). **(b)** Comparative expression of cardiac sarcomeric genes in MST-reprogrammed hiCMs vs. primary cardiac fibroblasts. Transcripts per million for expression. Error bars: mean ± SEM.

**Figure S9.**
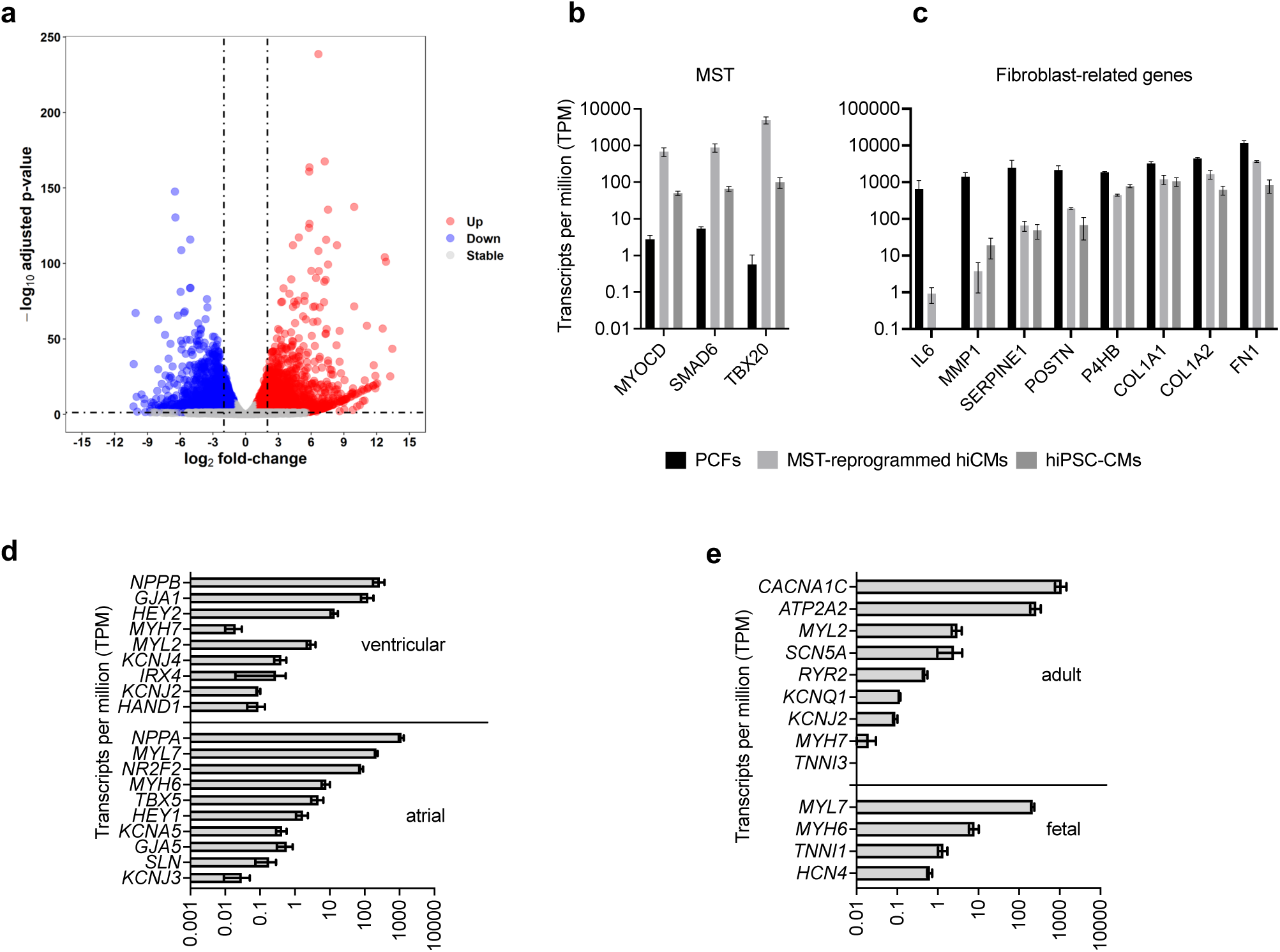
Transcriptional reprogramming of MST-derived hiCMs toward a cardiac state. **(a)** Volcano plot displaying 9,293 differentially expressed genes in MST-differentiated hiCMs relative to untransduced PCFs (FDR < 0.05). Comparative expression of cardiac genes in MST-reprogrammed hiCMs vs. primary cardiac fibroblasts. Transcripts per million for expression of **(b)** *MYOCD*, *SMAD6*, *TBX20* **(c)** fibroblast-related genes **(d)** atrial and ventricular genes **(e)** cardiac adult and fetal genes.

**Table S1.** Published transcription factor combinations for direct differentiation of human fibroblasts into cardiomyocytes. CF: cardiac fibroblast, TTF: tail tip fibroblasts, DF: dermal fibroblasts, FF: foreskin fibroblasts, hESC-FB: human embryonic stem cell-derived fibroblasts, hiPSC-CMs: human iPSC derived cardiomyocytes. FC: flow cytometry, IF: immunofluorescence. * published after completion of our screen.

| Publication | Starting human cell type | Virus | Transcription factor cocktail | Additional enhancers | Initial screening reporter/readout | Cardiac markers | Spontaneous calcium flux | Spontaneous contraction in human cells |
| --- | --- | --- | --- | --- | --- | --- | --- | --- |
| Nam/Olson PNAS, 2013 | Adult CF<br>Adult DF<br>Neonatal FF | Retrovirus<br>amphotropic<br>fresh | GATA4, TBX5, MYOCD, HAND2 or GATA4, MEF2C, TBX5, MYOCD, HAND2 (in absence of microRNAs) | miR-1, miR-133 | TNNT2 expression (FC) | Neonatal FF 2 weeks: TNNT2<sup>+</sup> (25.7% with microRNAs, 19.6% transcription factors alone)<br>Adult CF 2 weeks: TNNT2<sup>+</sup> (10.4% with microRNAs, 7.27% transcription factors alone)<br>Adult DF 2 weeks: TNNT2<sup>+</sup> (4.43% with microRNAs) | 8 weeks: 'low percentage' | 11 weeks: 'small subset' |
| Wada/Ieda PNAS, 2013 | Adult CF, pediatric CF | Retrovirus<br>VSVG<br>fresh | GATA4, MEF2C, TBX5, MYOCD, MESP1 | None | Cardiac gene induction (RT-qPCR) | 4 weeks: α-actinin<sup>+</sup> (5.5%)<br>4 weeks: TNNT2<sup>+</sup> (5.9%) | 4 weeks: ~1%, "low frequency" | Contraction in co-culture with mouse CMs |
| Fu/Srivastava Stem Cell Rep, 2013 | hESC-FB, adult DF, CF | Retrovirus<br>VSVG<br>fresh | GATA4, MEF2C, TBX5, MYOCD, MESP1, ESRRG, ZFPM2 | TGFβ1 (20 ng/mL) (in absence of MYOCD and ZFPM2) | MYH6 promoter reporter (FC)<br>TNNT2 expression (FC) | hESC-FB 2 weeks: αMHC-mCherry<sup>+</sup> (18.1%), TNNT2<sup>+</sup> (10.7%), αMHC-mCherry:cTNT<sup>++</sup> (13.0%)<br>DF 2 weeks: TNNT2<sup>+</sup> (3.71%)<br>CF 2 weeks: TNNT2<sup>+</sup> (1.77%) | 4 weeks: only with stimulation (20% of mCherry<sup>+</sup>) | Not described |
| Christoforou/Leong Sci Rep, 2017 | DF | Lentivirus<br>VSVG<br>fresh | GATA4, MEF2C, TBX5, MYOCD, NKX2-5 | miR-1, miR-133a, JAK1i | ACTN2 expression (IF) | 2 weeks: TNNT2<sup>+</sup> (0.21% transcription factors alone, 3.8% with micro-RNAs, 3.8% with JAK1i) | 7 days, but subsided by day 12-14 | 4 weeks: not detected |
| Mohamed/Srivastava Circulation, 2017 | Immortalized CF | Retrovirus<br>amphotropic<br>fresh | GATA4, MEF2C, TBX5, MYOCD | SB431542 (2.6 μM)<br>XAV939 (5 μM) | α-MHC-GFP reporter<br>TNT-GCaMP (FC) | 3 weeks: TNNT2-GFP<sup>+</sup> (13.4% with small molecules) | 10 days: spontaneous transients, homogenous transients within 3 weeks | Not described |
| Zhou/Qian Cell Stem Cell, 2019 | Adult CF | Retrovirus<br>VSVG<br>frozen | GATA4, MEF2C, TBX5 (polycistronic) | miR-133 | TNNT2 expression (FC) | 2 weeks: TNNT2<sup>+</sup> (2.83%) (41.8% with antibiotic selection) | 20 days: spontaneous transients | Contraction in co-culture with mouse neonatal CMs |
| Garry/Olson Nat Cell Biol., 2021* | Adult CF | Retrovirus<br>amphotropic<br>fresh | GATA4, MEF2C, TBX5, MYOCD, HAND2, PHF7 | None | TNNT2 and α-Actinin expression (FC) | 3 weeks: TNNT2<sup>+</sup> (~3.5%)<br>α-Actinin<sup>+</sup> (~10.0 %)<br>TNNT2+α-Actinin<sup>+</sup> (~3.0%) | Not described for human cells | Not described for human cells |
| Tang/Zhou Circulation, 2022* | hESC-FB, CF, DF | Retrovirus<br>VSVG<br>fresh | GATA4, MEF2C, TBX5, TBX20 | miR-133 | α-MHC and α-Actinin expression (FC) | 2 weeks: α-MHC<sup>+</sup> (30.3%)<br>α-Actinin<sup>+</sup> (23.8%) | 1 month: spontaneous transients in co-culture with hiPSC-CMs | Contraction in co-culture with hiPSC-CMs |
| Wang/Qian Cell Stem Cell, 2022* | hESC-FB | Lentivirus<br>Retrovirus | MEF2C, ASCL1 | None | cTnT and α-Actinin expression (FC) | 2 weeks: cTnT+α-Actinin<sup>+</sup> (6.53%) | Not described for human cells | Not described for human cells |
| This work | Pediatric CF | Lentivirus<br>VSVG<br>frozen | MYOCD, SMAD6, TBX20 | XAV939 (1 μM)<br>FGF2 (100 ng/ mL) | MYH6/TNN T2 promoter reporter and Ca<sup>2+</sup> flux (high content kinetic imaging) | 6 days: MYH6-mNeon<sup>+</sup> (33%)<br>TNNT2-mScarlet<sup>+</sup> (17%)<br>MYH6-mNeon-TNNT2-mScarlet<sup>++</sup> (13%)<br>without antibiotic selection | 7 days: spontaneous transients, homogenous transients within 3 weeks | Spontaneous contraction starting at 20 days |

**Table S2.**
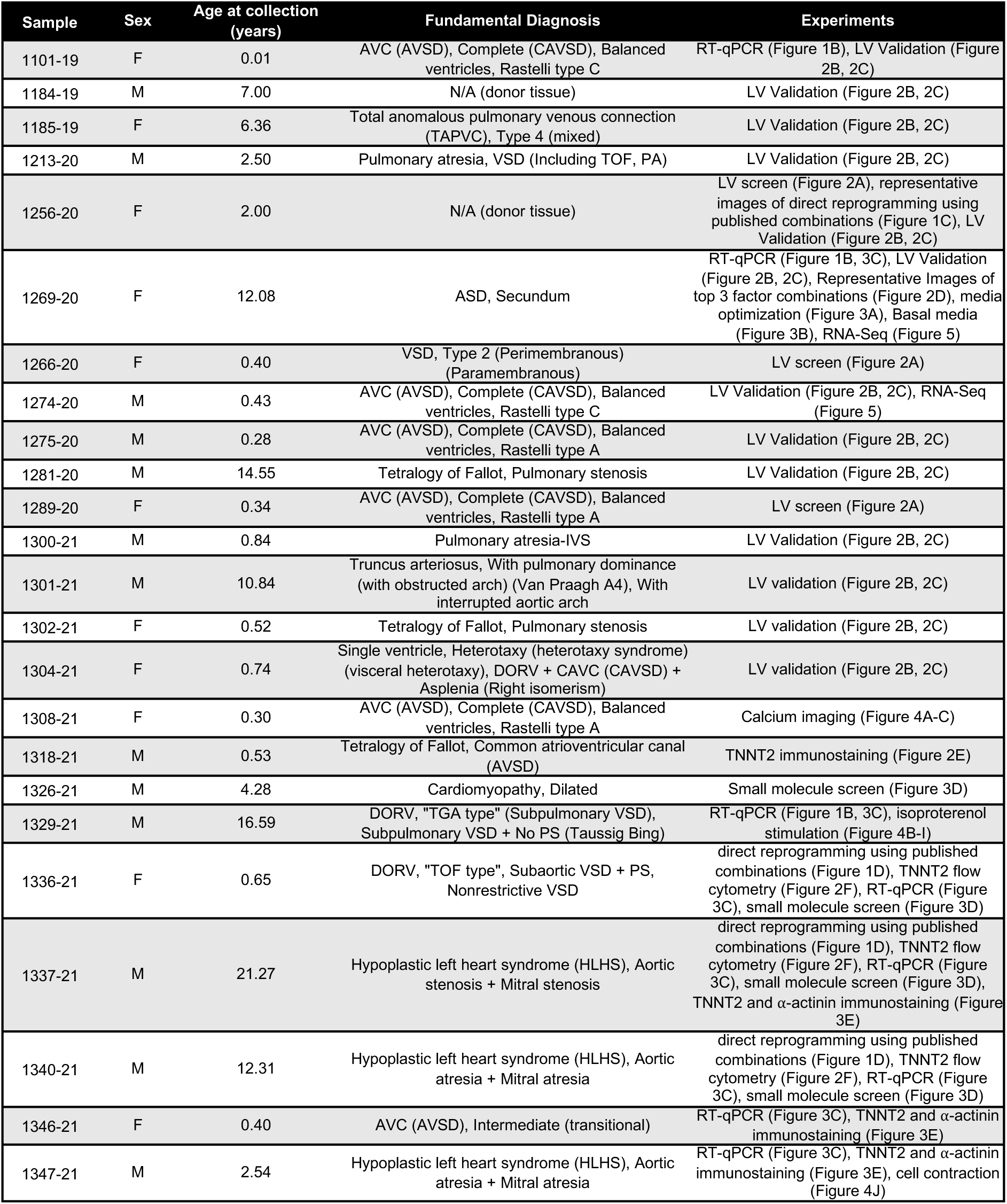
Patient sample information

**Table S3.** Antibodies for flow cytometry (FC) and immunocytochemistry (ICC).

| Marker | Antibody | Source | Application |
| --- | --- | --- | --- |
| TNNT2 | Alexa Fluor 647 Mouse anti-cardiac troponin T | BD Biosciences #565744 | 1:33 (FC) |
| TNNT2 | Rabbit anti-cardiac troponin T | Abcam ab45932 | 1:200 (ICC) |
| α-actinin | Mouse anti-a-actinin | Sigma A7811 | 1:500 (ICC) |
| CD31 | Alexa Fluor 647 Mouse anti-Human CD31 | BD Pharmingen #558094 | 1:100 (FC) |
| CD90 | APC Mouse Anti-Human CD90 | BD Pharmingen #559869 | 1:50 (FC) |
| Vimentin | Mouse anti-vimentin | BD Pharmingen #550513 | 1:100 (FC) |

**Table S4.**
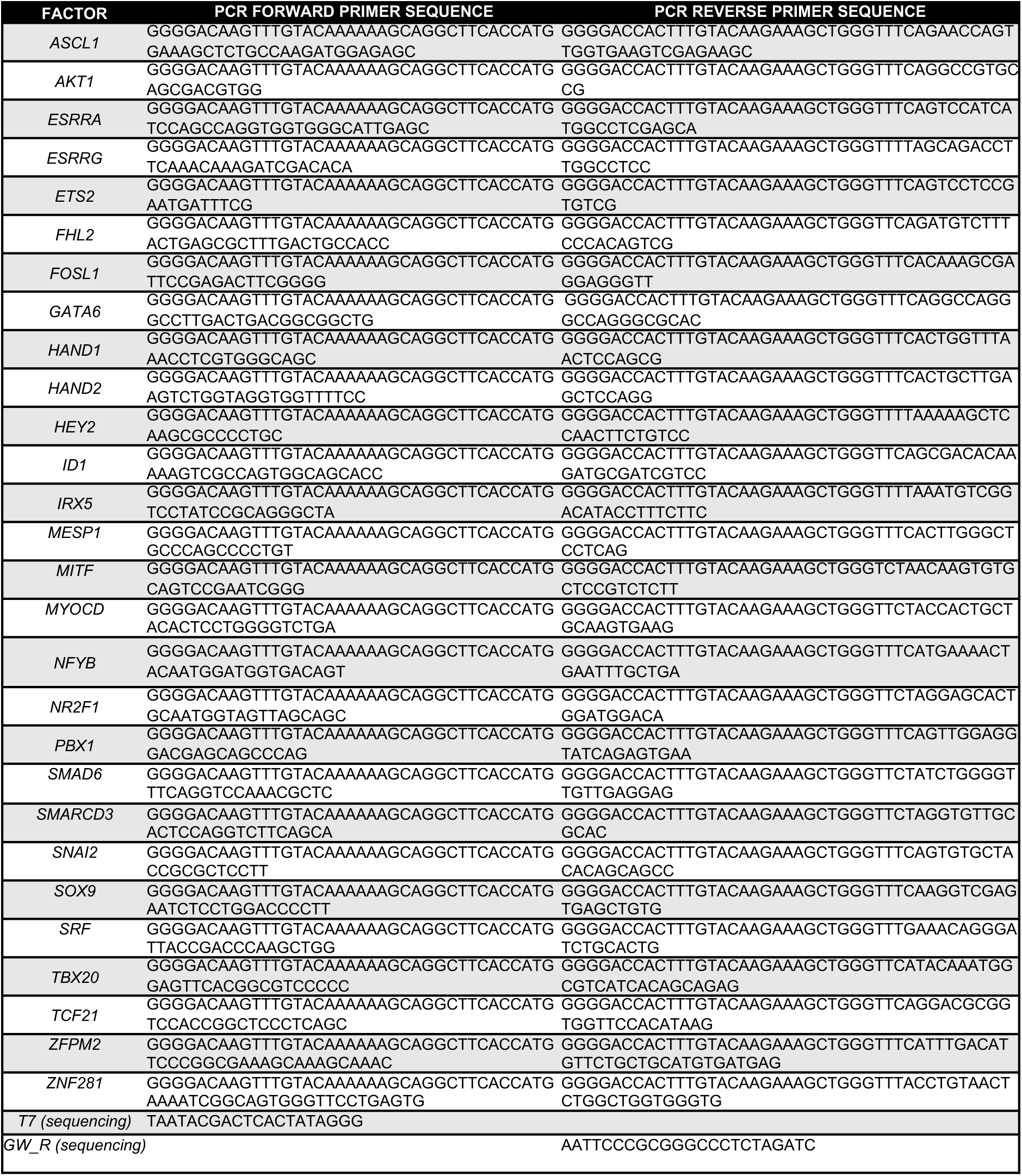
Primers used for Gateway cloning and Sanger sequencing

**Table S5.** Addgene deposit details of candidate factors for reprogramming

| Factor | Plasmid Addgene # | Source Sequence | Inclusion Rationale |
| --- | --- | --- | --- |
| ASCL1 | 184382 | ORF amplified from tetO-ALN (Addgene Plasmid #43918) | Used in direct reprogramming |
| AKT1 | 170682 | ORF amplified from pWZL Neo Myr Flag AKT1 (Addgene Plasmid #20422) | Reported to enhance reprogramming of mouse fibroblasts to functional cardiomyocytes (Zhou/Olson 2015) |
| ESRRA | 170683 | ORF amplified from MGC Human ESRRA Sequence-Verified cDNA (Dharmacon Cloneld:6527166) | Mogrify prediction (fibroblast to adult heart)<br>Mogrify prediction (cardiac fibroblast to adult heart) |
| ESRRG | 170684 | ORF amplified from pMX-hESRRG (a gift from Deepak Srivastava) | Used in published cocktail combinations |
| ETS2 | 170685 | ORF amplified from FLAG-Ets2 (Addgene Plasmid #28128) | Reported role in direct differentiation of human fibroblast to cardiac progenitors (Islas/Schwartz 2012) |
| FHL2 | 170686 | ORF amplified from pcDNA3.1-C-(k)DYK-FHL2 (Genscript Clone ID #OHu26596) | Mogrify prediction (cardiac fibroblast to adult heart) |
| FOSL1 | 170687 | ORF amplified from p6599 MSCV-IP N-HAonly FOSL1 (Addgene Plasmid #34897) | Mogrify prediction (whole blood to cardiac myocyte) |
| GATA4 | 46030 | Plasmid was a gift from John Gearhart (Addgene Plasmid #46030) | Used in published cocktail combinations<br>Mogrify prediction (fibroblast to cardiac myocyte)<br>Mogrify prediction (fibroblast to adult heart) |
| GATA6 | 170688 | ORF amplified from pEN_TT 3XFLAG-wtGATA6-3XAU1 (Addgene Plasmid #72612) | Mogrify prediction (fibroblast to cardiac myocyte)<br>Mogrify prediction (cardiac fibroblast to cardiac myocyte) |
| HAND1 | 170689 | ORF amplified from MGC Human HAND1 Sequence-Verified cDNA (Dharmacon Cloneld:3162118) | Mogrify prediction (fibroblast to adult heart)<br>Mogrify prediction (cardiac fibroblast to adult heart) |
| HAND2 | 170690 | ORF amplified from MGC Fully Sequenced Human HAND2 cDNA Dharmacon #40003700 | Used in published cocktail combinations |
| HEY2 | 170691 | ORF amplified from MGC Human HEY2 Sequence-Verified cDNA (Dharmacon Cloneld:3945225) | Mogrify prediction (fibroblast to adult heart) |
| ID1 | 170692 | ORF amplified from pcDNA3 hId1 (Addgene Plasmid #16061) | Mogrify prediction (fibroblast to cardiac myocyte)<br>Mogrify prediction (cardiac fibroblast to cardiac myocyte) |
| IRX5 | 170693 | ORF amplified from pcDNA3.1-C-(k)DYK-IRX5 (Genscript Clone ID #OHu18967) | Mogrify prediction (fibroblast to adult heart)<br>Mogrify prediction (cardiac fibroblast to adult heart) |
| MEF2C | 46031 | Plasmid was a gift from John Gearhart (Addgene Plasmid #46031) | used in published cocktail combinations<br>Mogrify prediction (fibroblast to adult heart)<br>Mogrify prediction (cardiac fibroblast to adult heart) |
| MESP1 | 170694 | ORF amplified from pSAM2_mCherry_Mesp1 (Addgene #72687) | Used in published cocktail combinations |
| MITF | 170695 | ORF amplified from pEGFP-N1-MITF-A (Addgene Plasmid #38132) | Mogrify prediction (cardiac fibroblast to adult heart) |
| MYOCD | 170696 | ORF amplified from MGC Human MYOCD sequence-verified cDNA (Dharmacon, clone ID 9051897) | Used in published cocktail combinations |
| NFYB | 170697 | ORF amplified from MGC Human NFYB sequence-verified cDNA (Dharmacon, clone ID 3680937) | Mogrify prediction (fibroblast to cardiac myocyte)<br>Mogrify prediction (cardiac fibroblast to cardiac myocyte) |
| NKX2-5 | 46029 | Plasmid was a gift from John Gearhart (Addgene Plasmid #46029) | used in published cocktail combinations<br>Mogrify prediction (fibroblast to adult heart)<br>Mogrify prediction (cardiac fibroblast to adult heart) |
| NR2F1 | 170698 | ORF amplified from MGC Human NR2F1 sequence-verified cDNA (Dharmacon, clone ID 3912075) | Mogrify prediction (fibroblast to cardiac myocyte)<br>Mogrify prediction (cardiac fibroblast to cardiac myocyte) |
| PBX1 | 170699 | ORF amplified from PBX1A-pCMV1 (Addgene #21029) | Mogrify prediction (fibroblast to cardiac myocyte)<br>Mogrify prediction (cardiac fibroblast to cardiac myocyte) |
| SMAD6 | 170700 | ORF amplified from CS2 Smad6 (Addgene #14960) | Mogrify prediction (fibroblast to cardiac myocyte)<br>Mogrify prediction (cardiac fibroblast to cardiac myocyte) |
| SMARCD3 | 170701 | ORF amplified from pBS-hBAF60c (Addgene #21036) | Reported to enhance reprogramming of mouse fibroblasts to functional cardiomyocytes (Christoforou/Leong 2013) |
| SNAI2 | 170702 | ORF amplified from MGC Human SNAI2 Sequence-Verified cDNA (Dharmacon, clone ID 3908245) | Mogrify prediction (PBMC to adult heart) |
| SOX9 | 170703 | ORF amplified from MGC Human SOX9 Sequence-Verified cDNA (Dharmacon, clone ID 6200521) | Mogrify prediction (cardiac fibroblast to cardiac myocyte) |
| SRF | 170704 | ORF amplified from pEntr-SRFfbio (Addgene #32971) | Reported to enhance reprogramming of mouse fibroblasts to functional cardiomyocytes (Christoforou/Leong 2013) |
| TBX20 | 170705 | ORF amplified from pcDNA3.1-C-(k)DYK-TBX20 (Genscript Clone ID #OHu10999) | Mogrify prediction (fibroblast to cardiac myocyte) |
| TBX5 | 46032 | Plasmid was a gift from John Gearhart (Addgene Plasmid #46032) | used in published cocktail combinations<br>Mogrify prediction (fibroblast to adult heart)<br>Mogrify prediction (cardiac fibroblast to adult heart) |
| TCF21 | 170706 | ORF amplified from pcDNA3.1-C-(k)DYK-TCF21 (Genscript Clone ID #OHu24607) | Mogrify prediction (cardiac fibroblast to cardiac myocyte) |
| ZFPM2 | 170707 | ORF amplified from pMX-hZFPM2, a gift from Deepak Srivastava | Used in published cocktail combinations |
| ZNF281 | 170708 | ORF amplified from MGC Human ZNF281 Sequence-Verified cDNA (Dharmacon Cloneld:6527166) | Reported to stimulate cardiac reprogramming of adult mouse fibroblasts (Zhou/Olson 2017) |

**Table S6.**
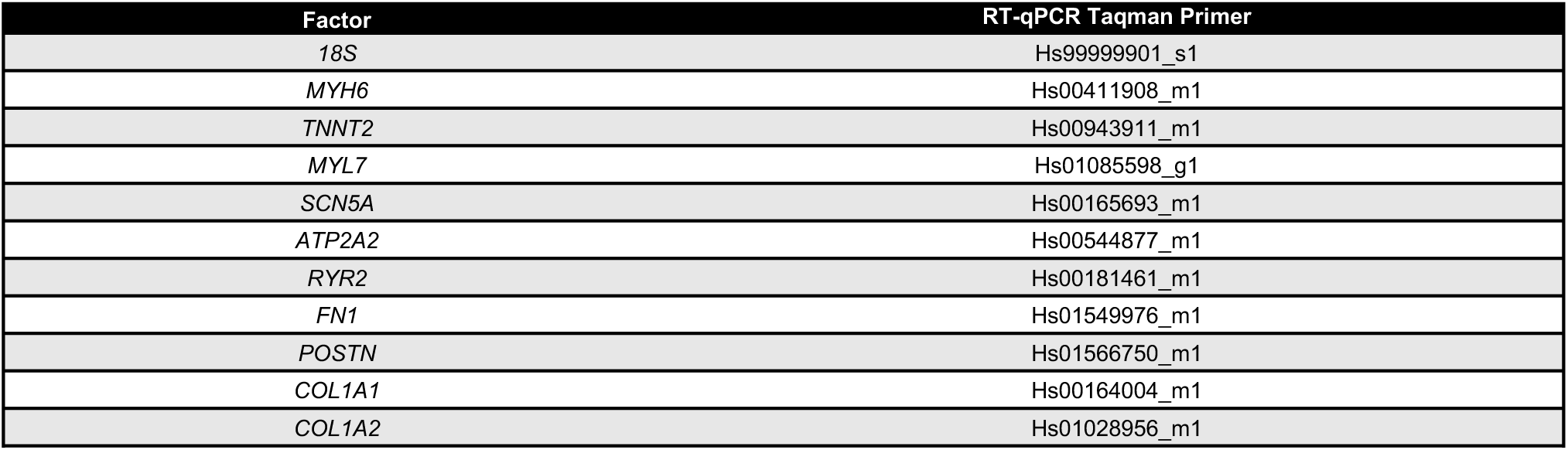
RT-qPCR primers

**Table S7.**
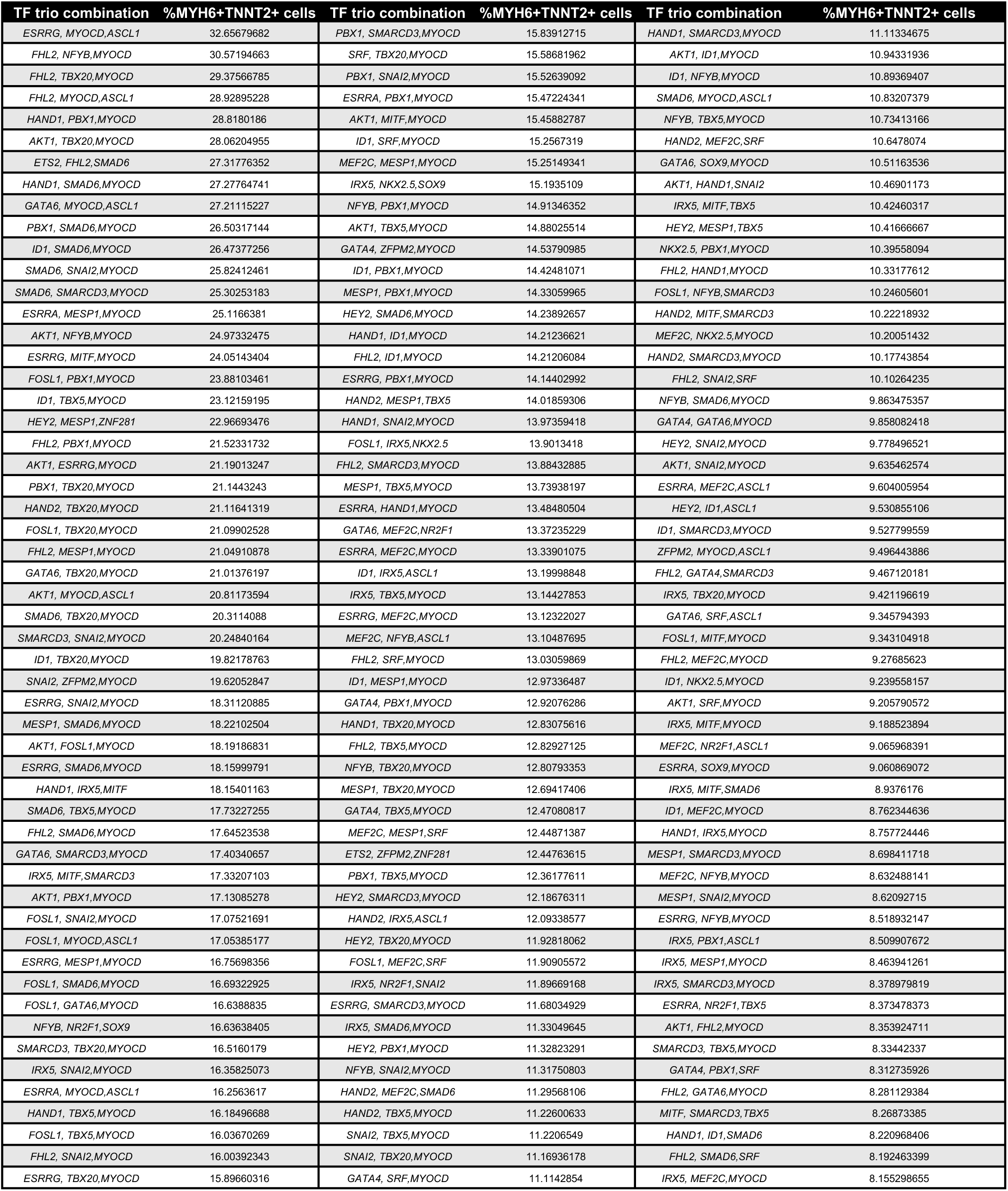
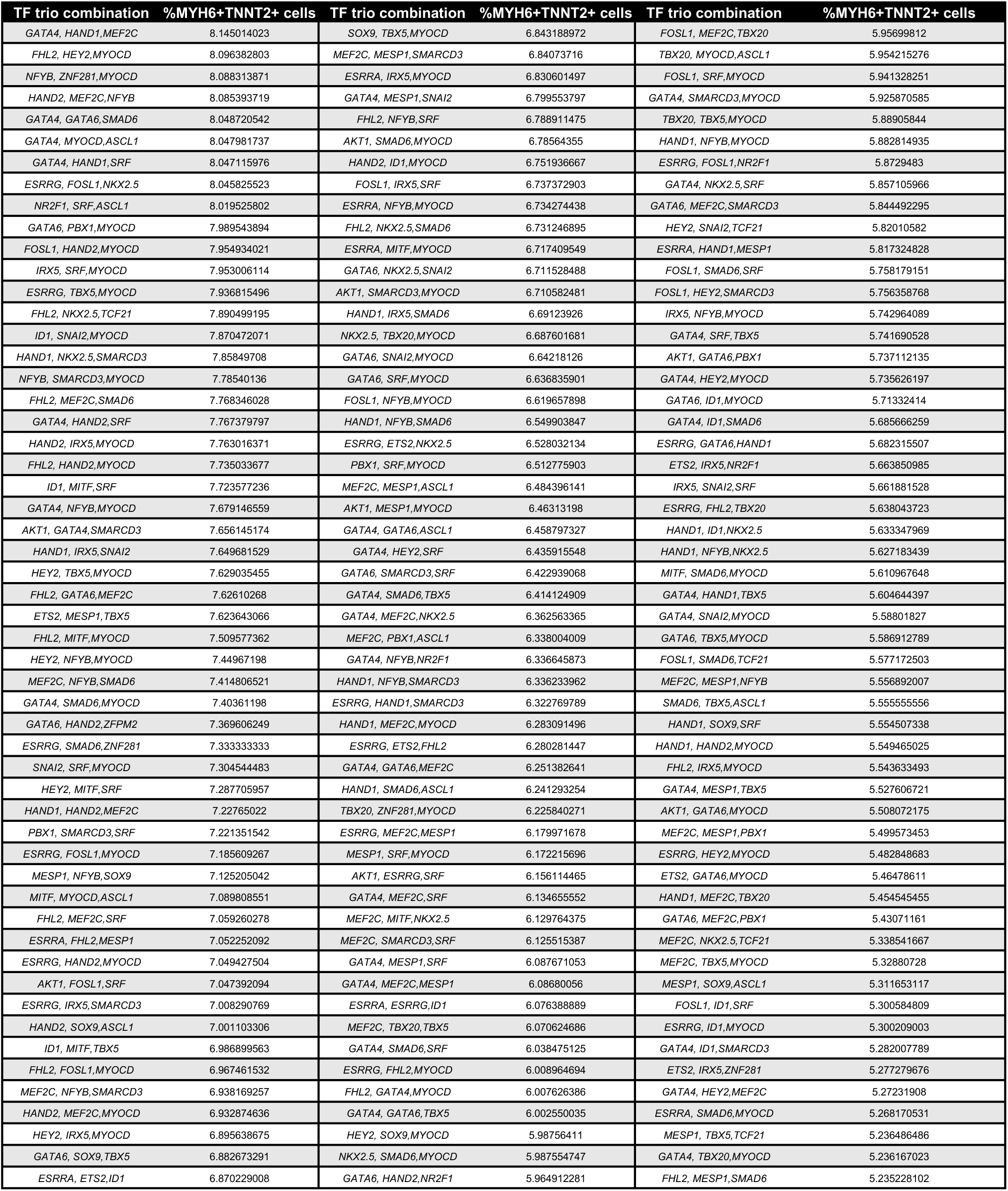
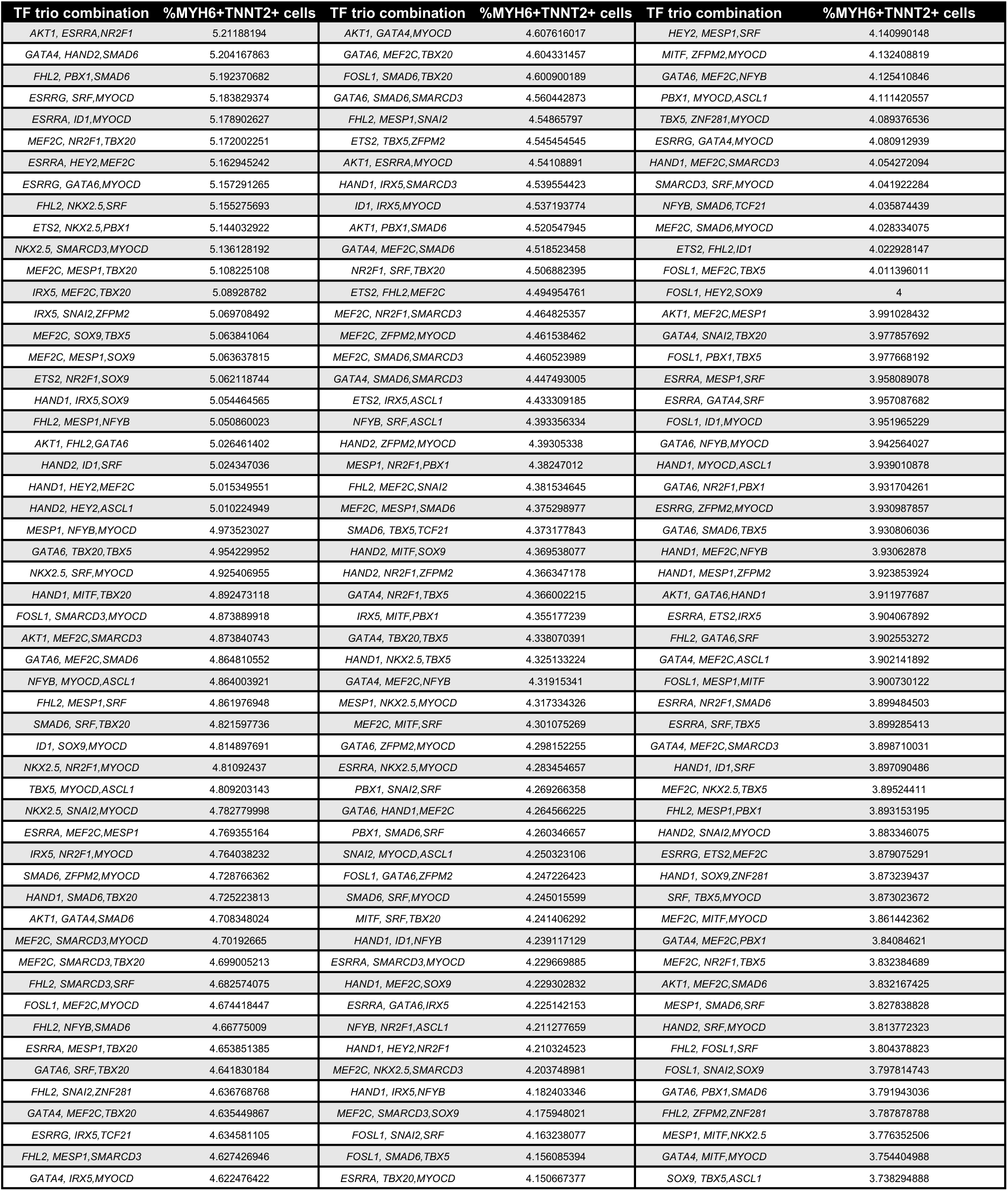
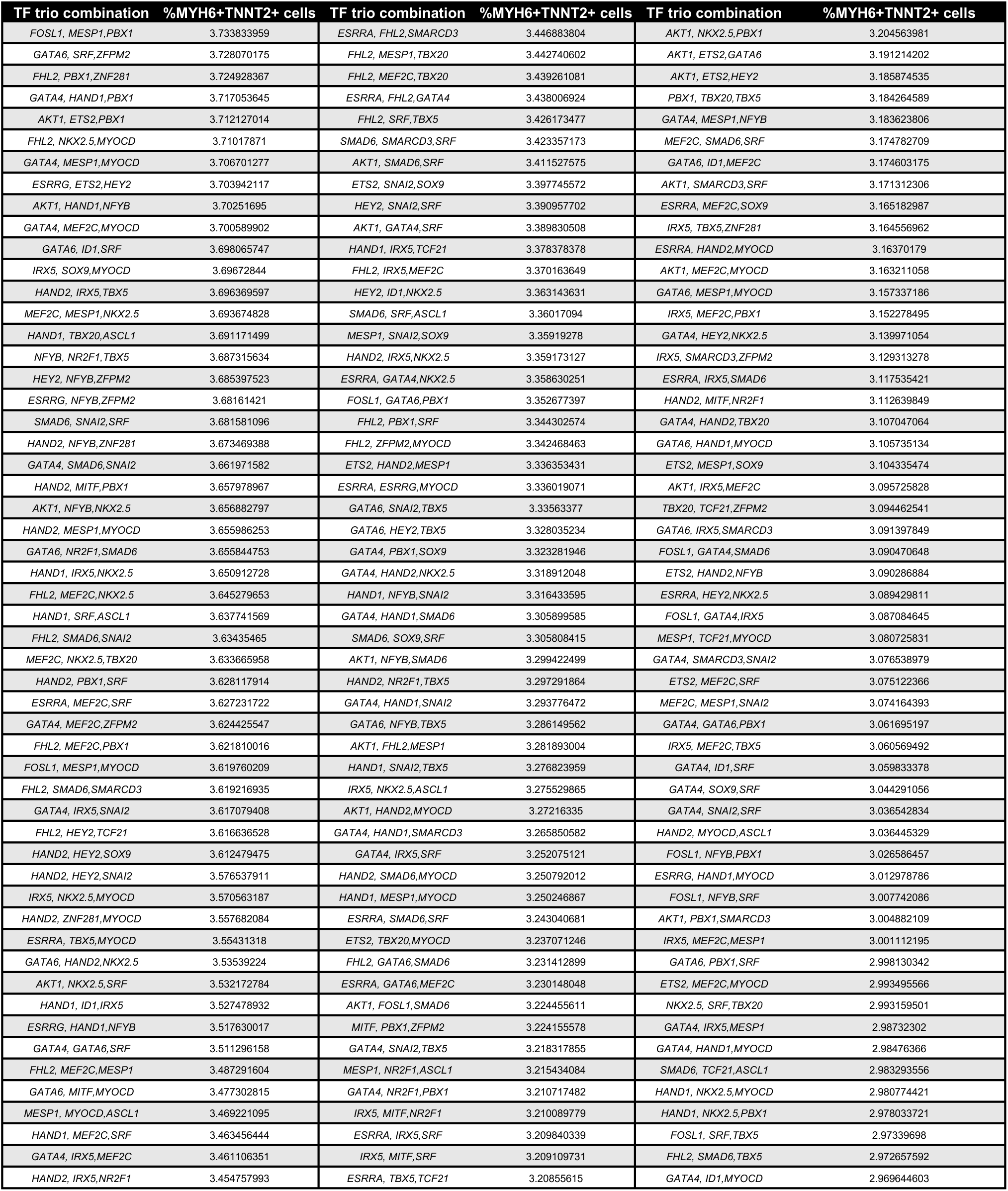
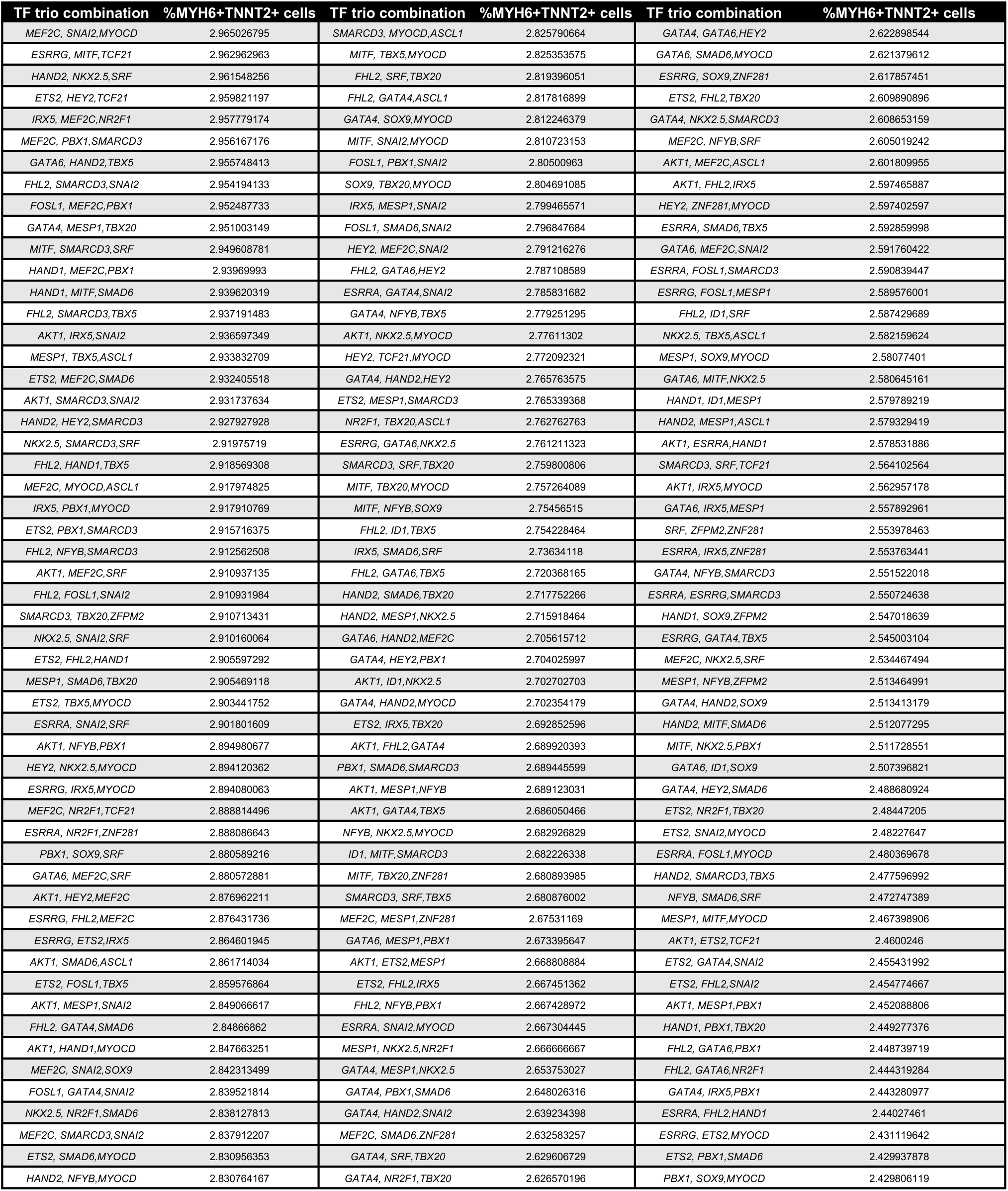
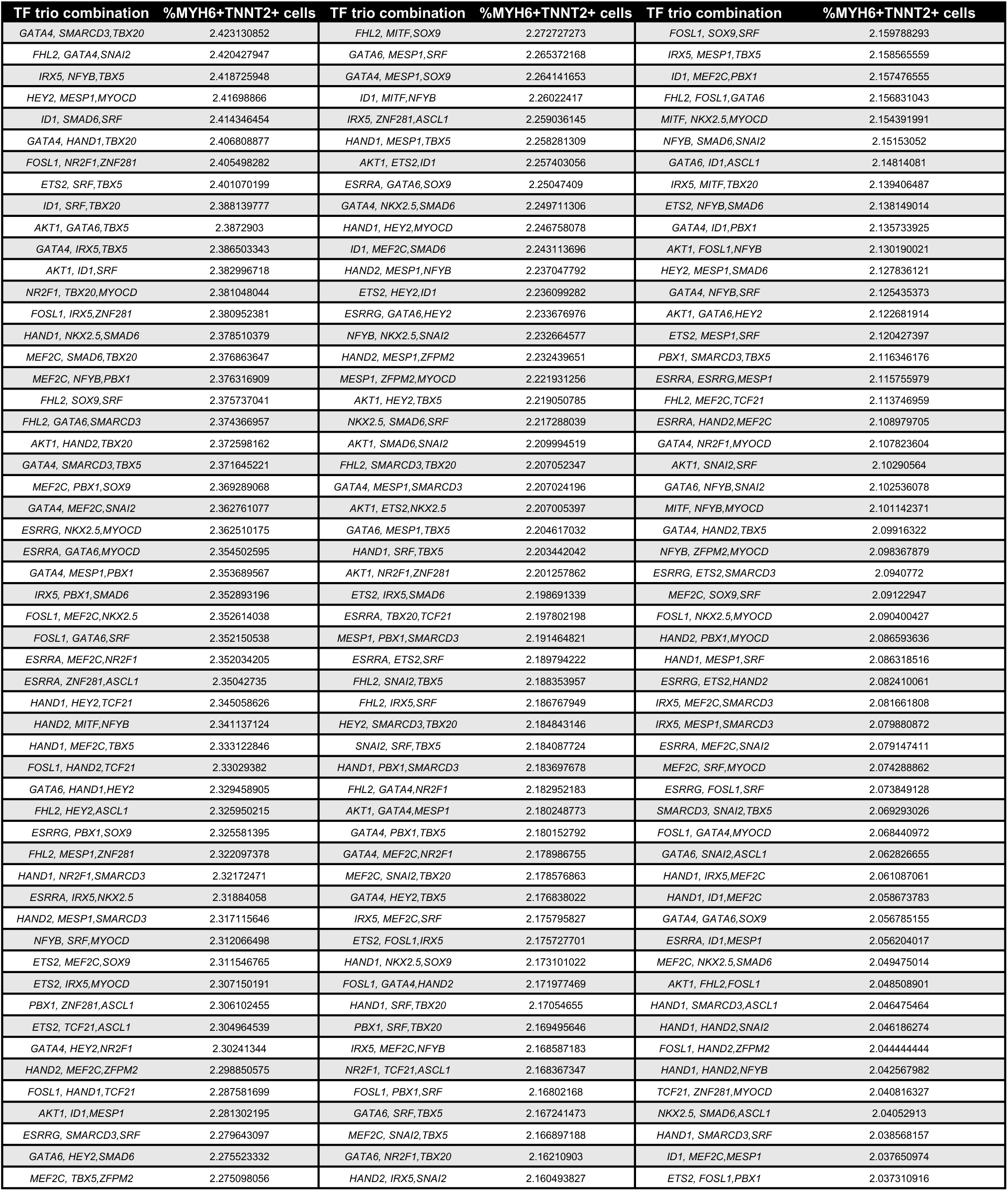
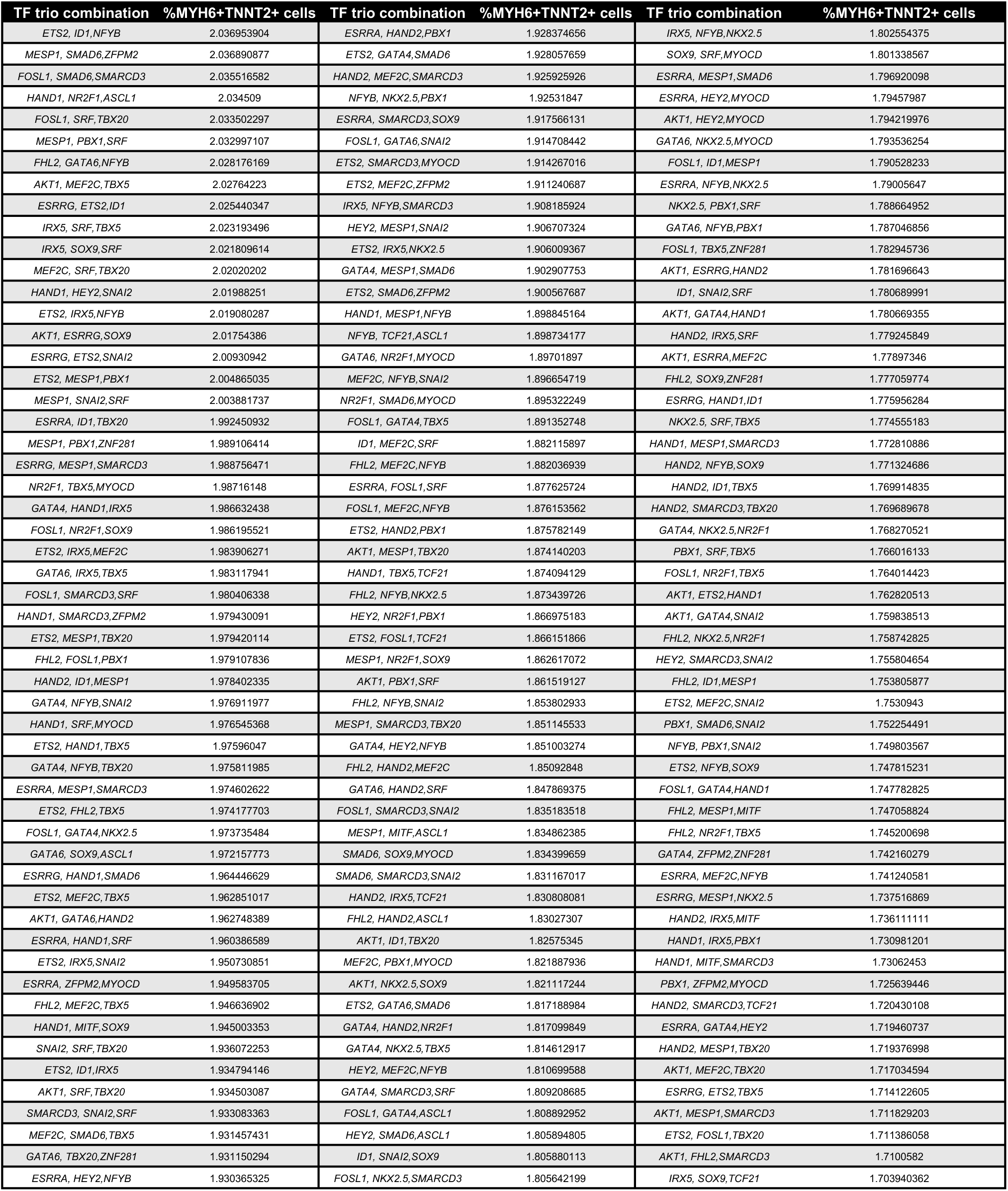
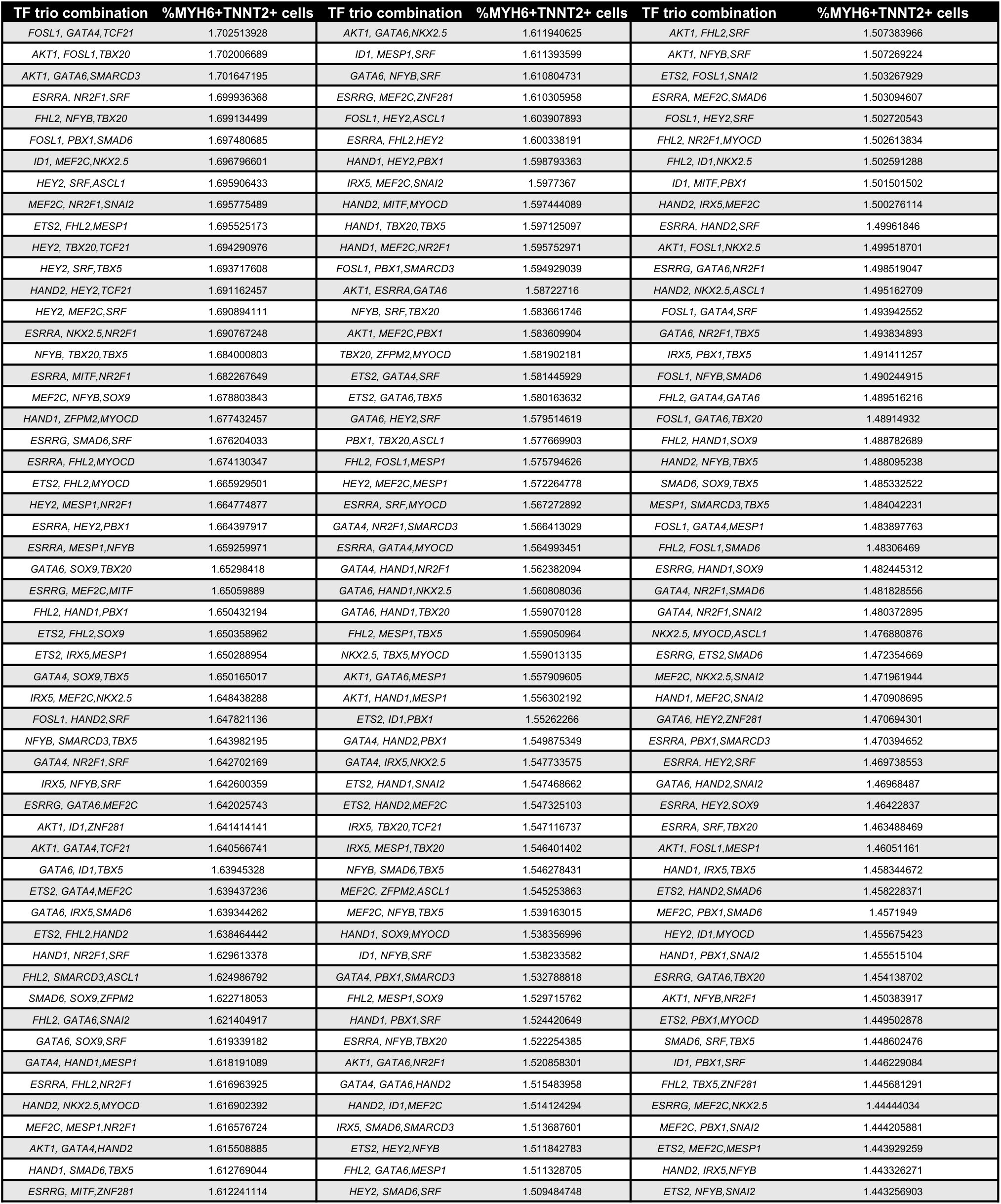
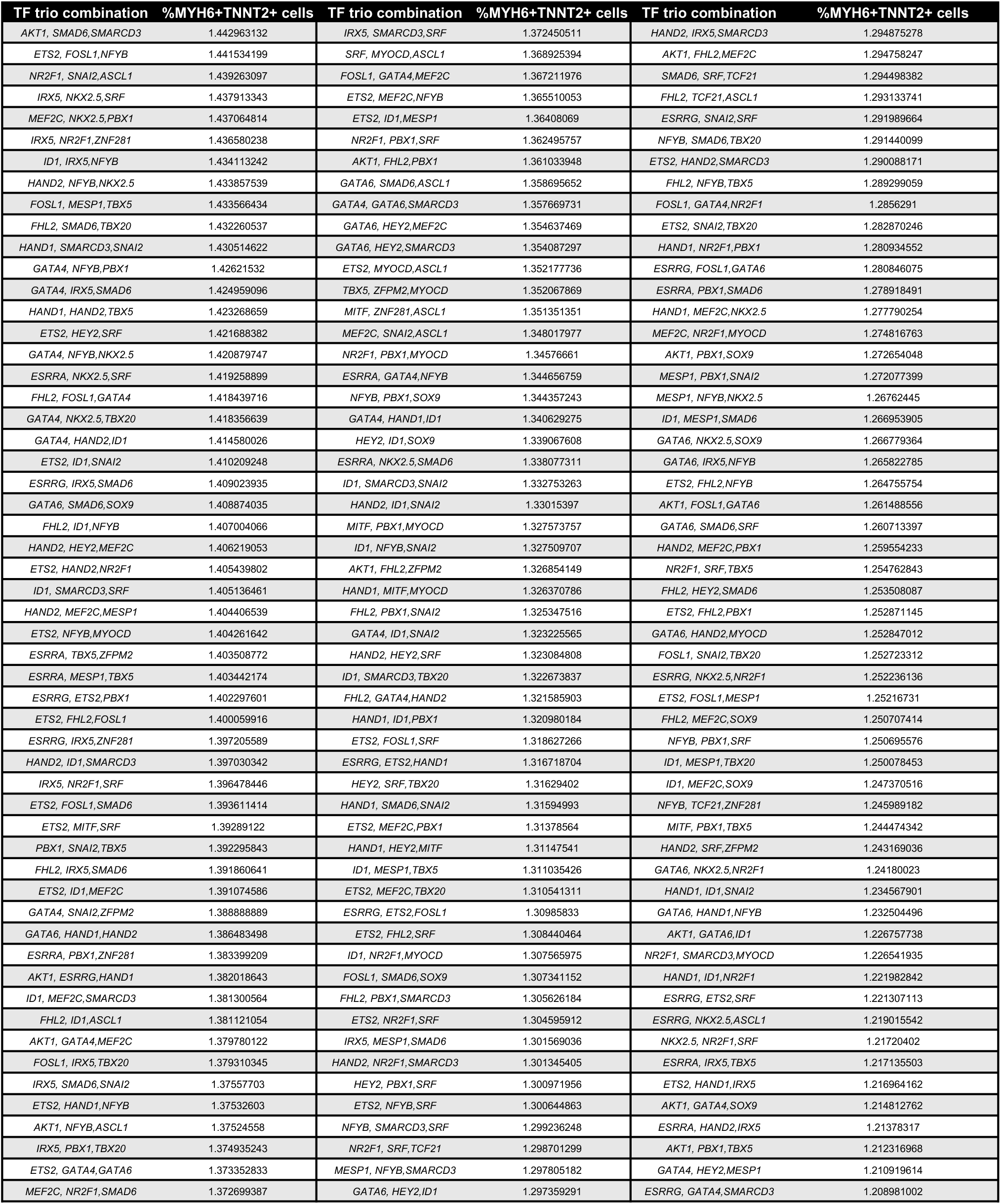
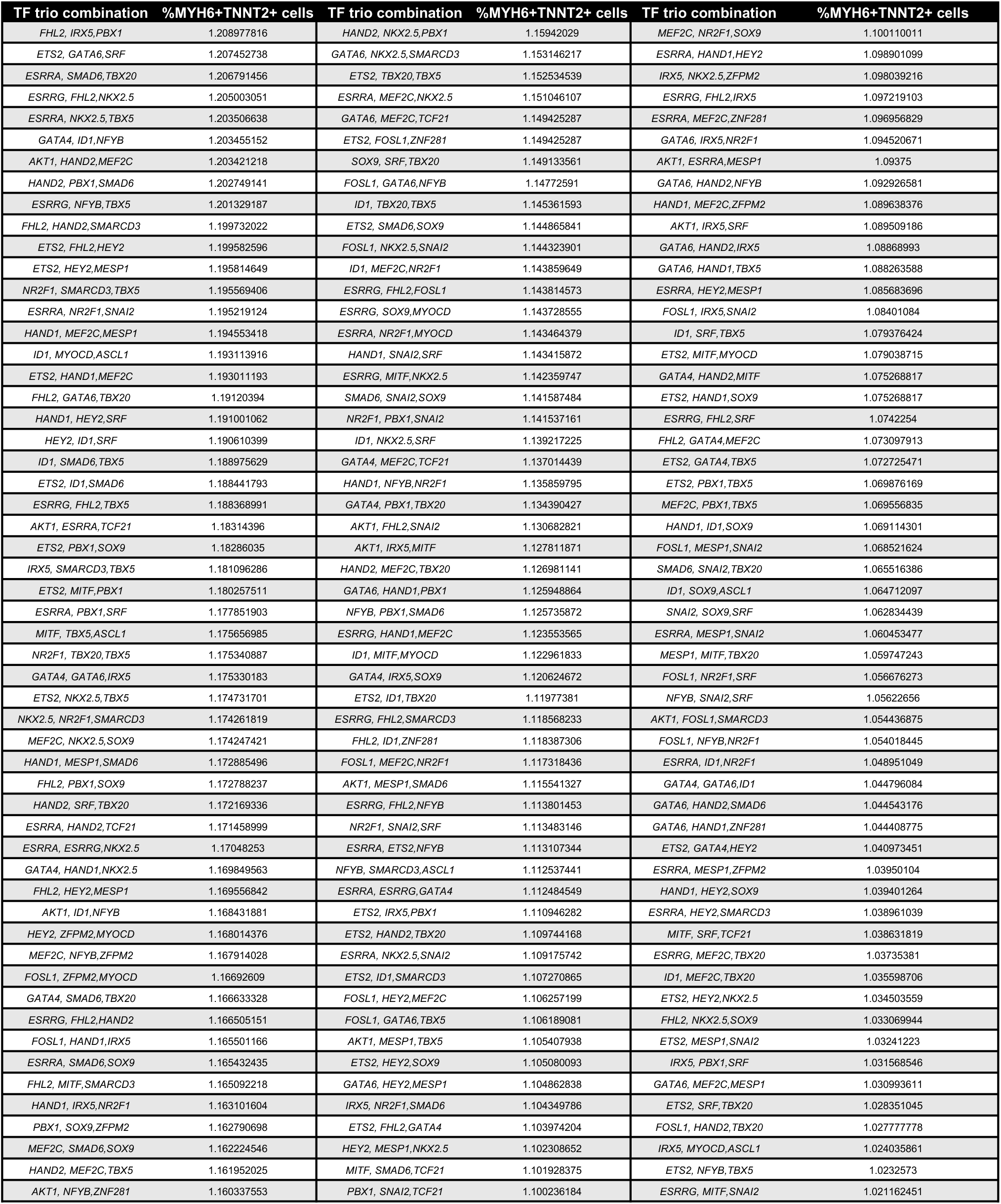
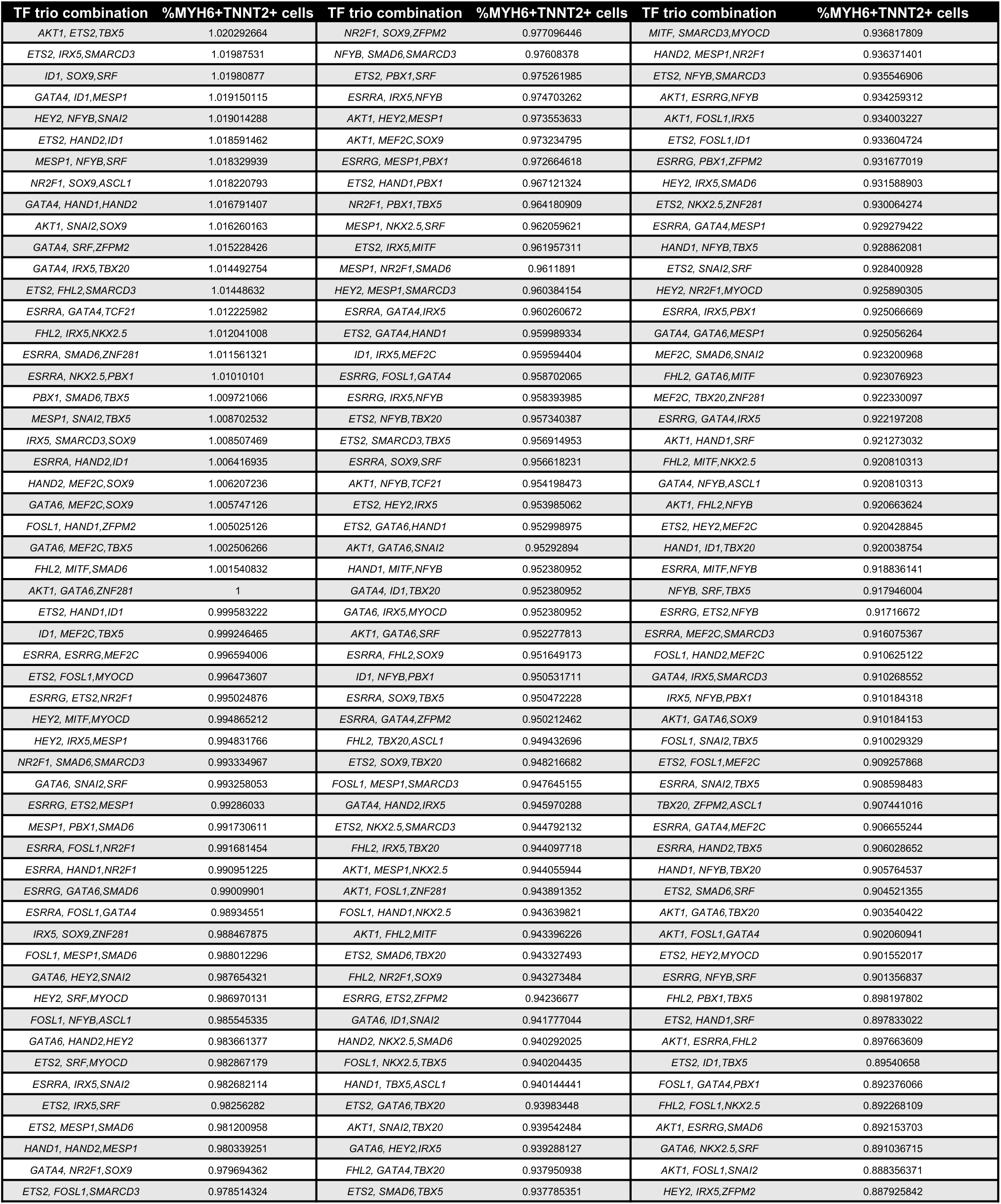
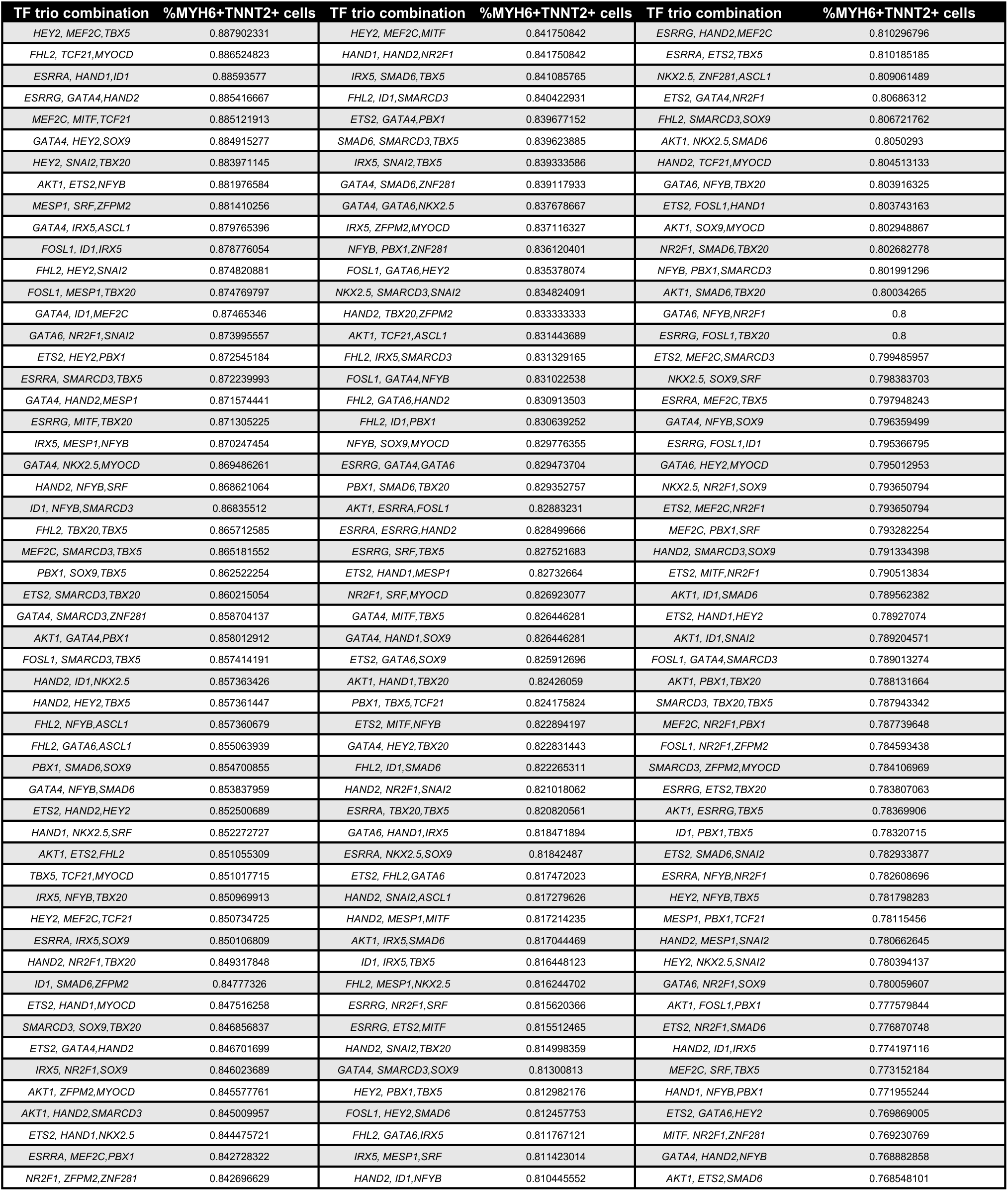
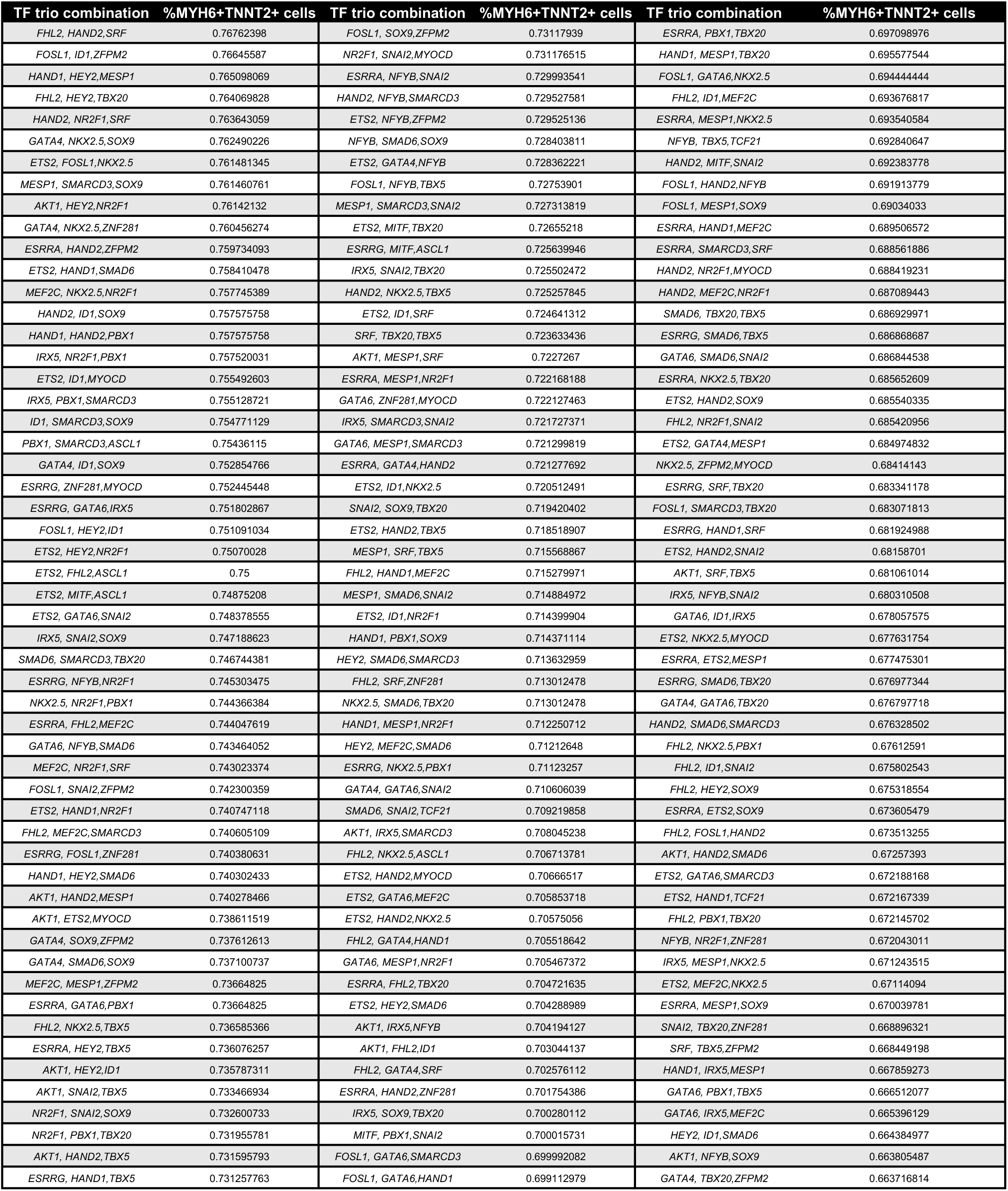
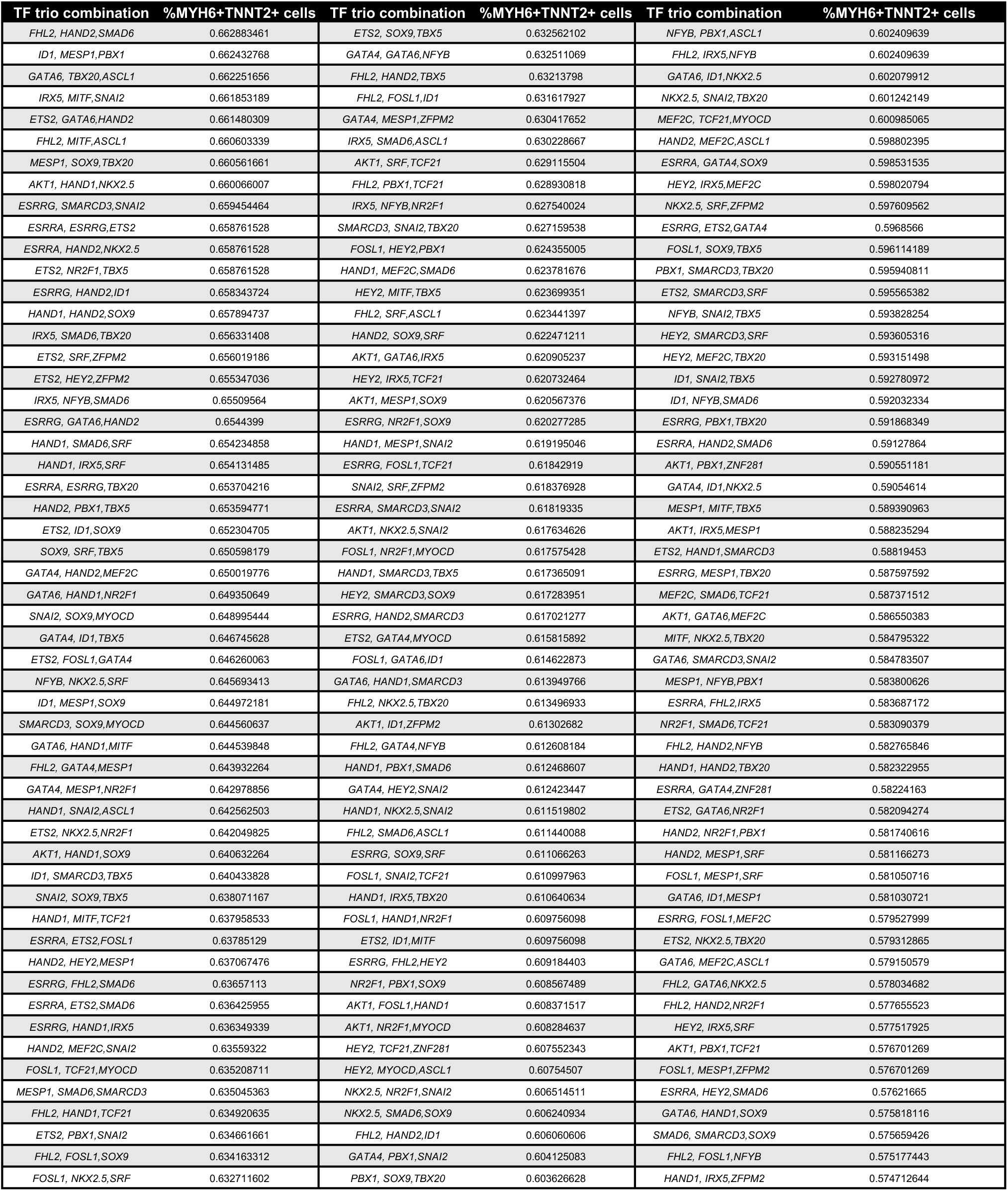
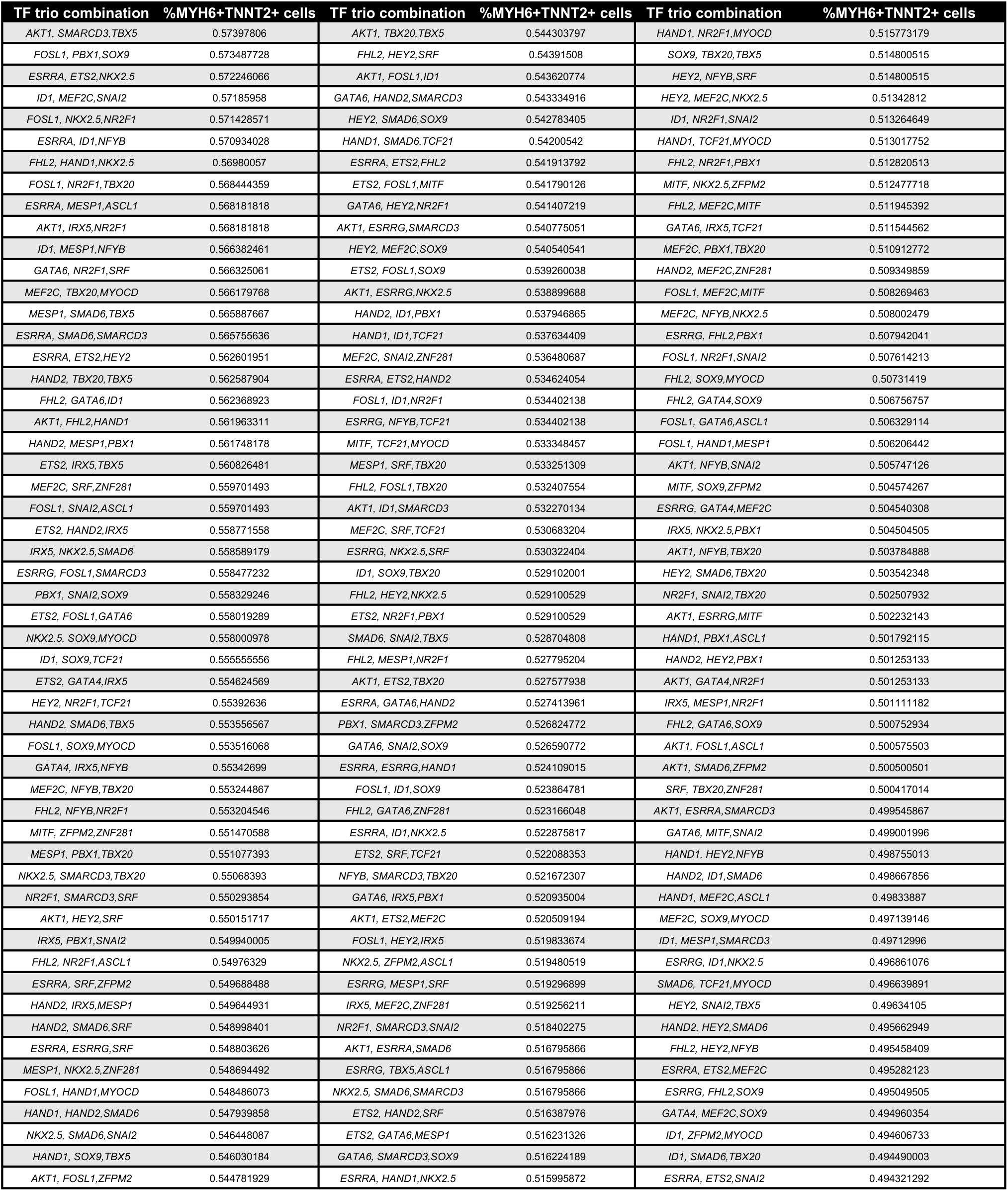
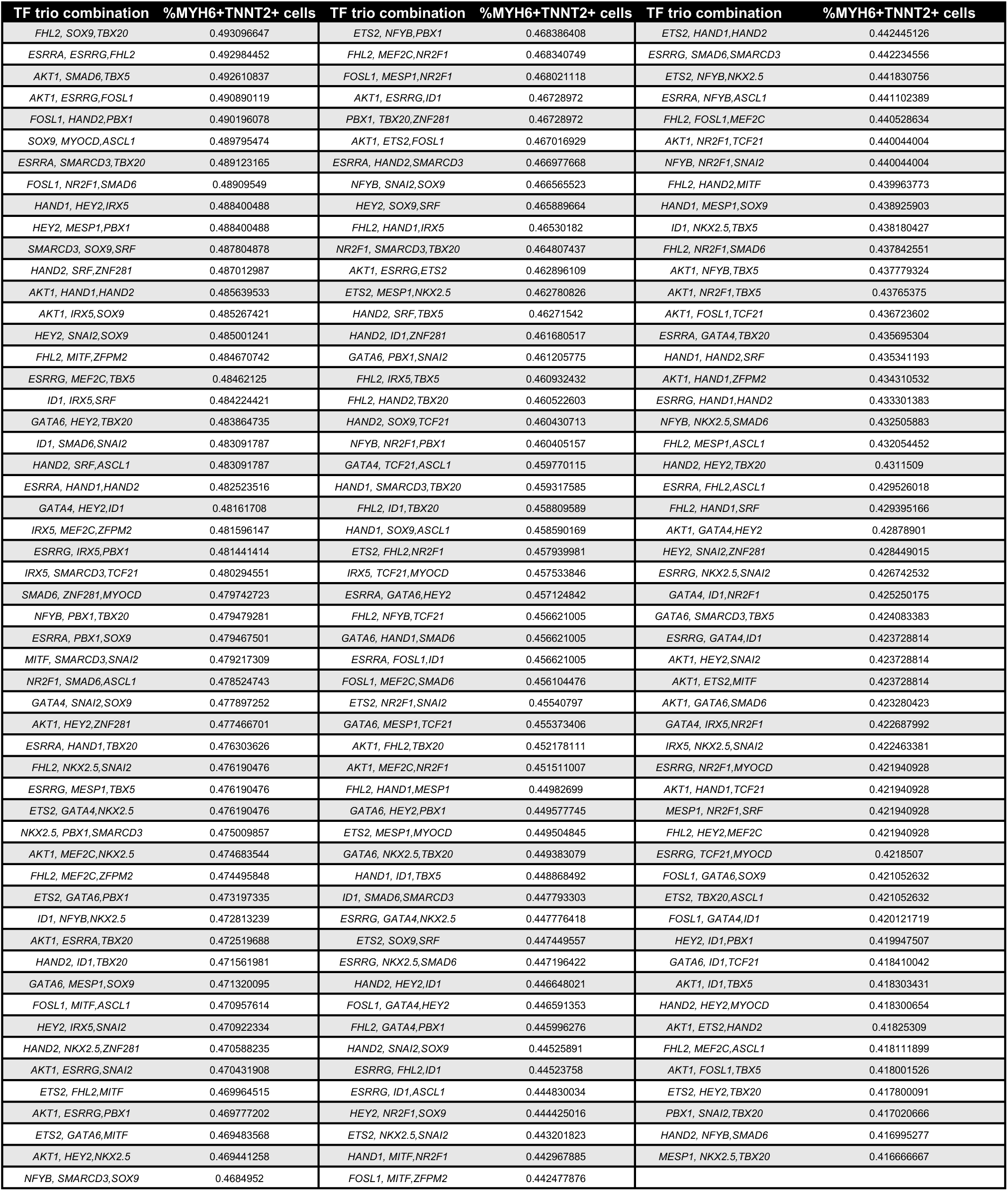
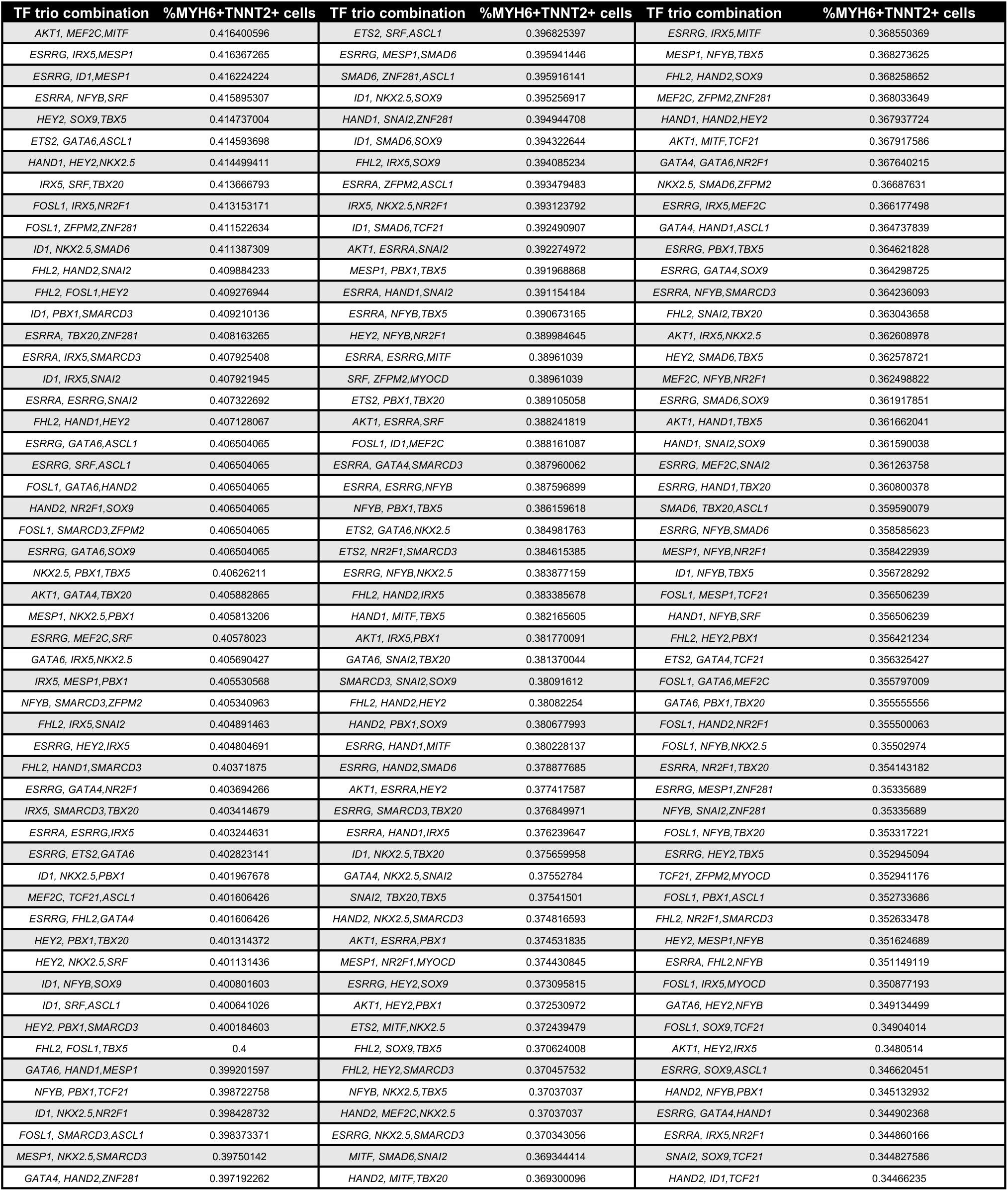
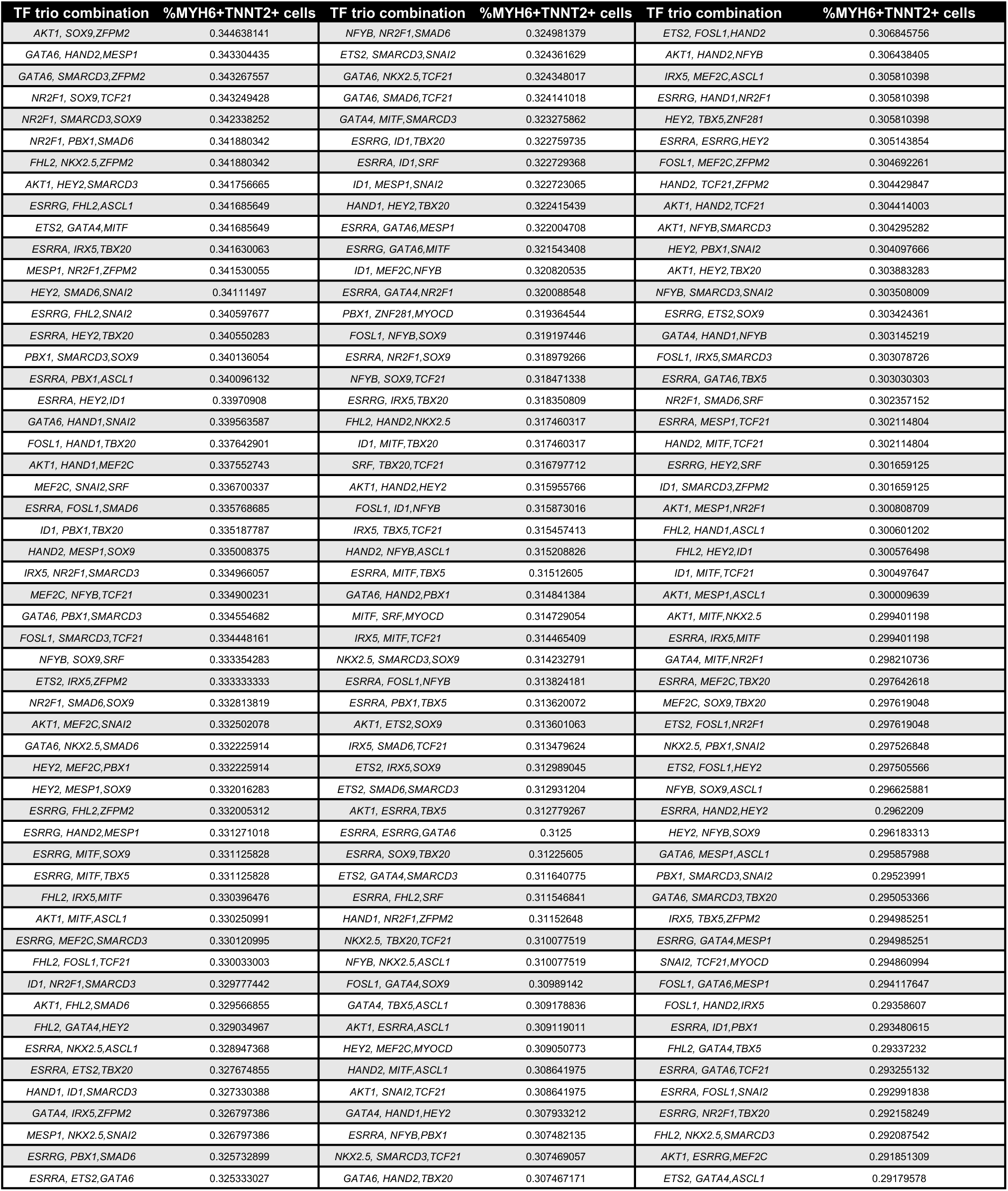
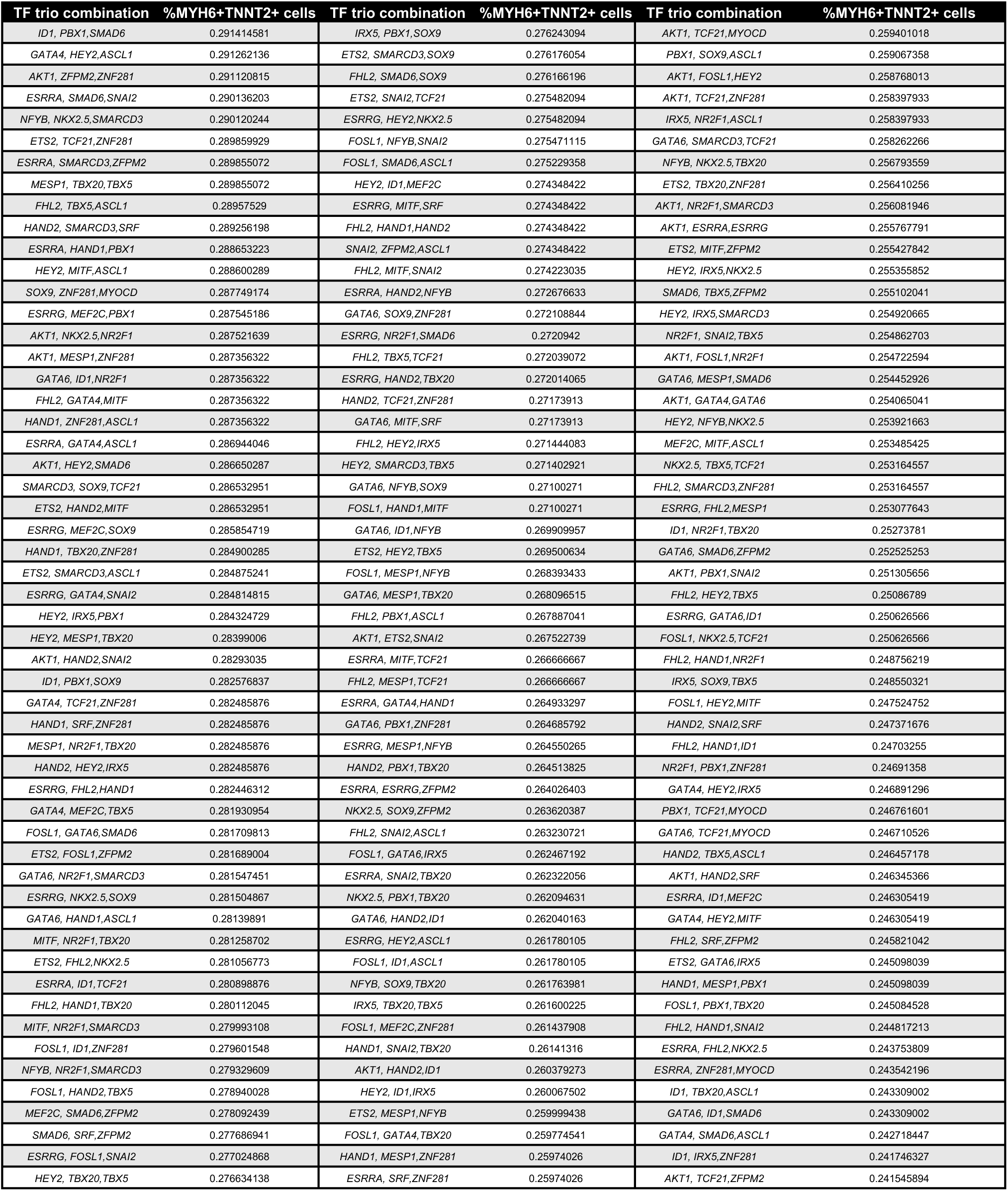
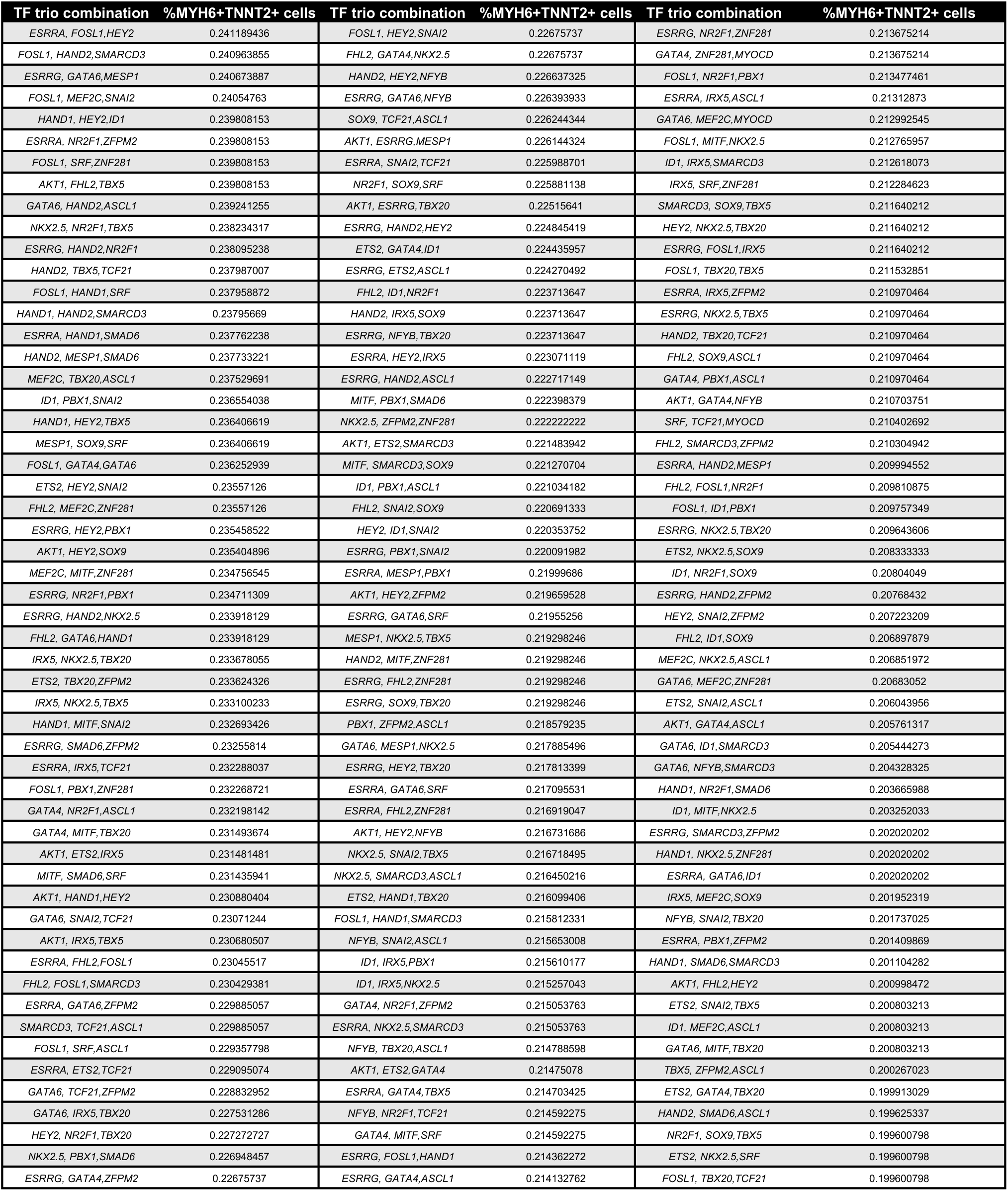
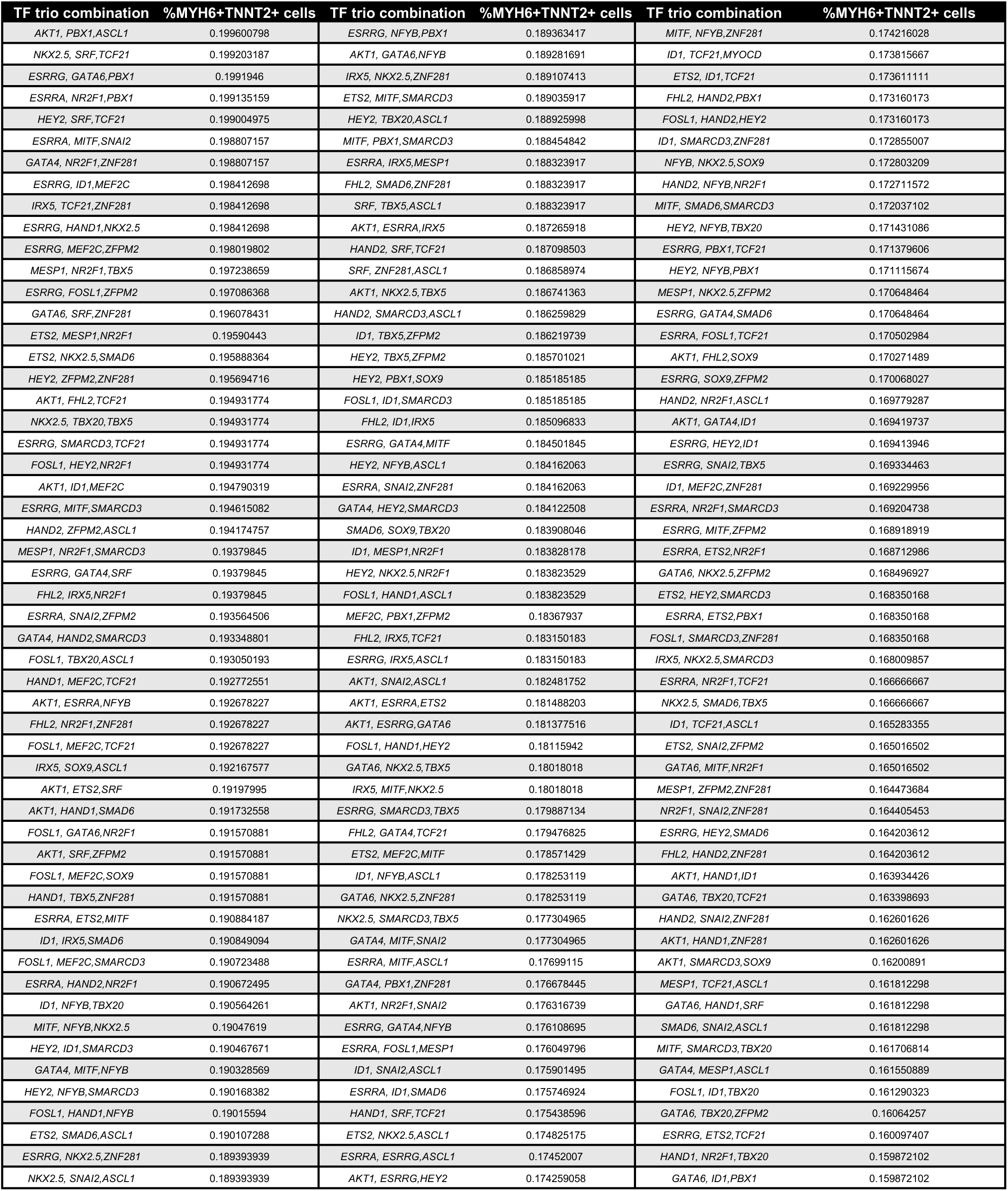
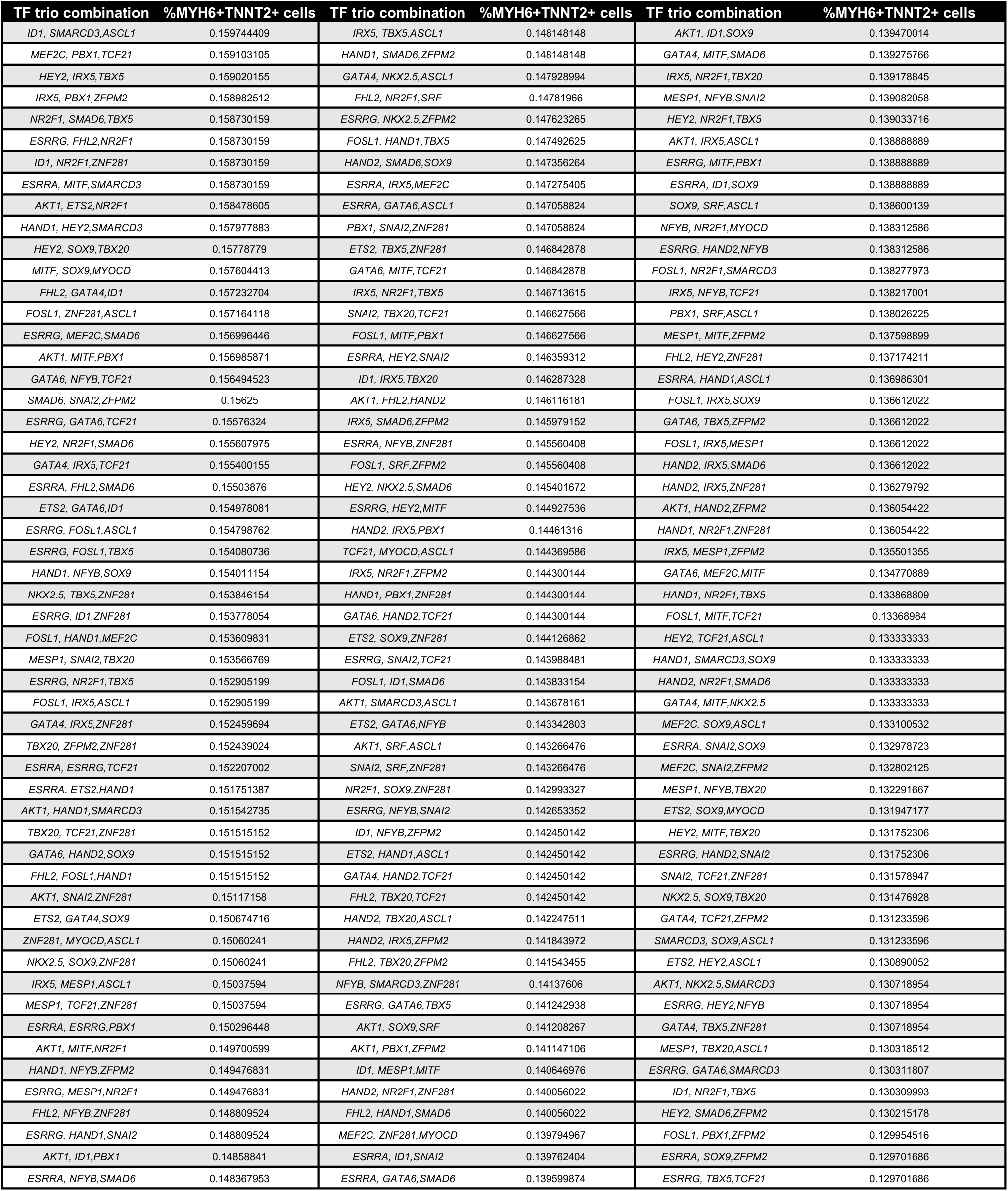
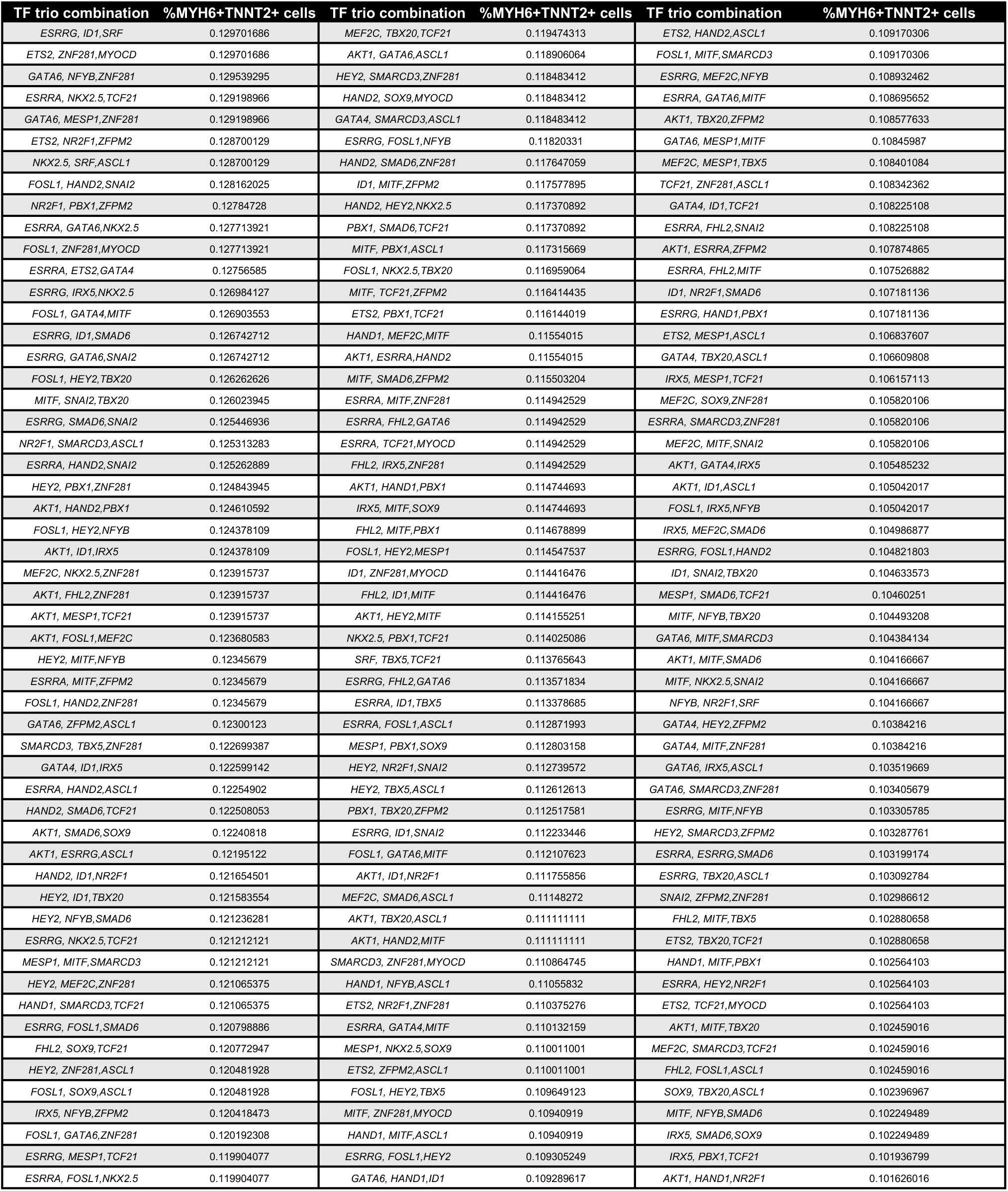
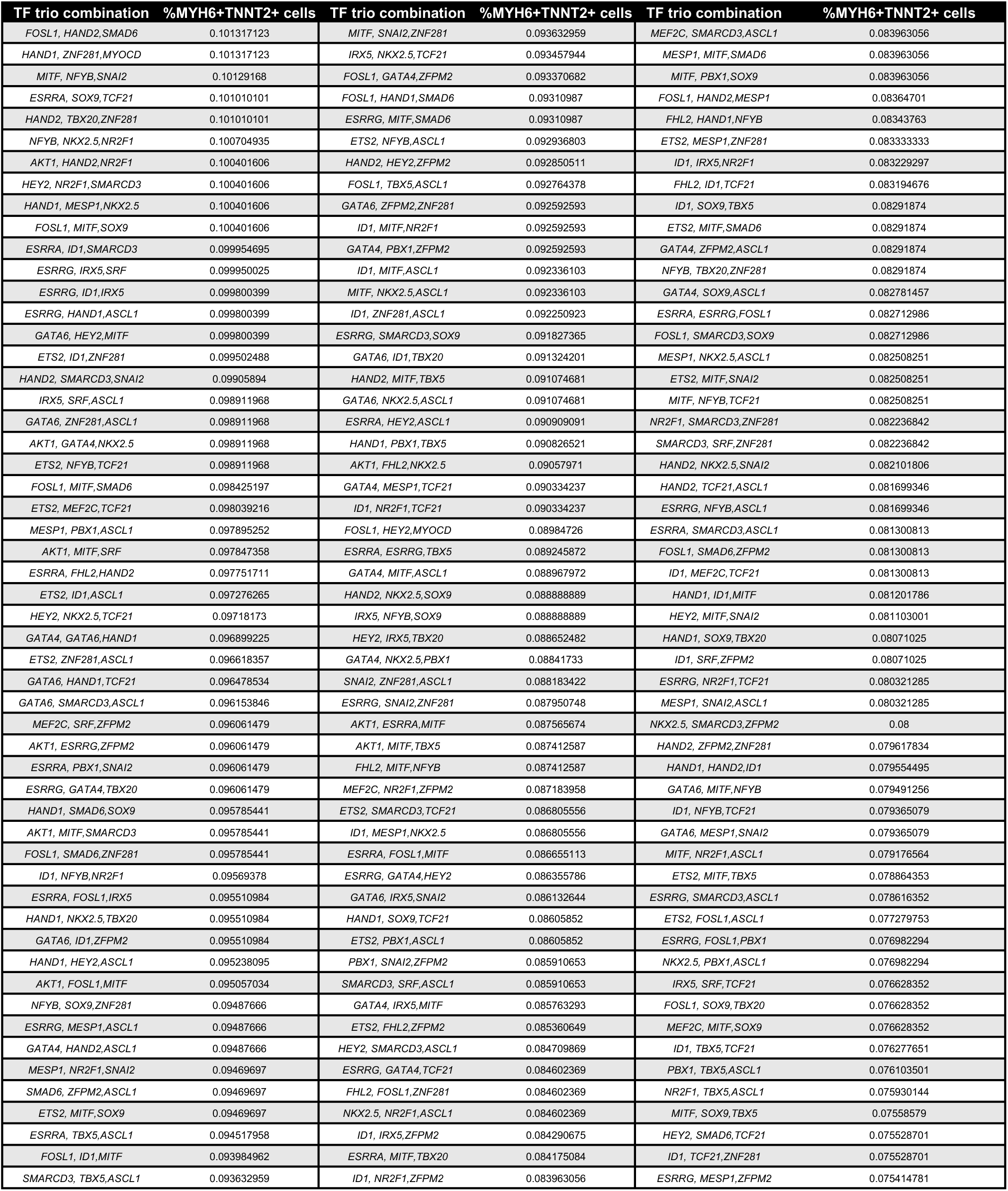
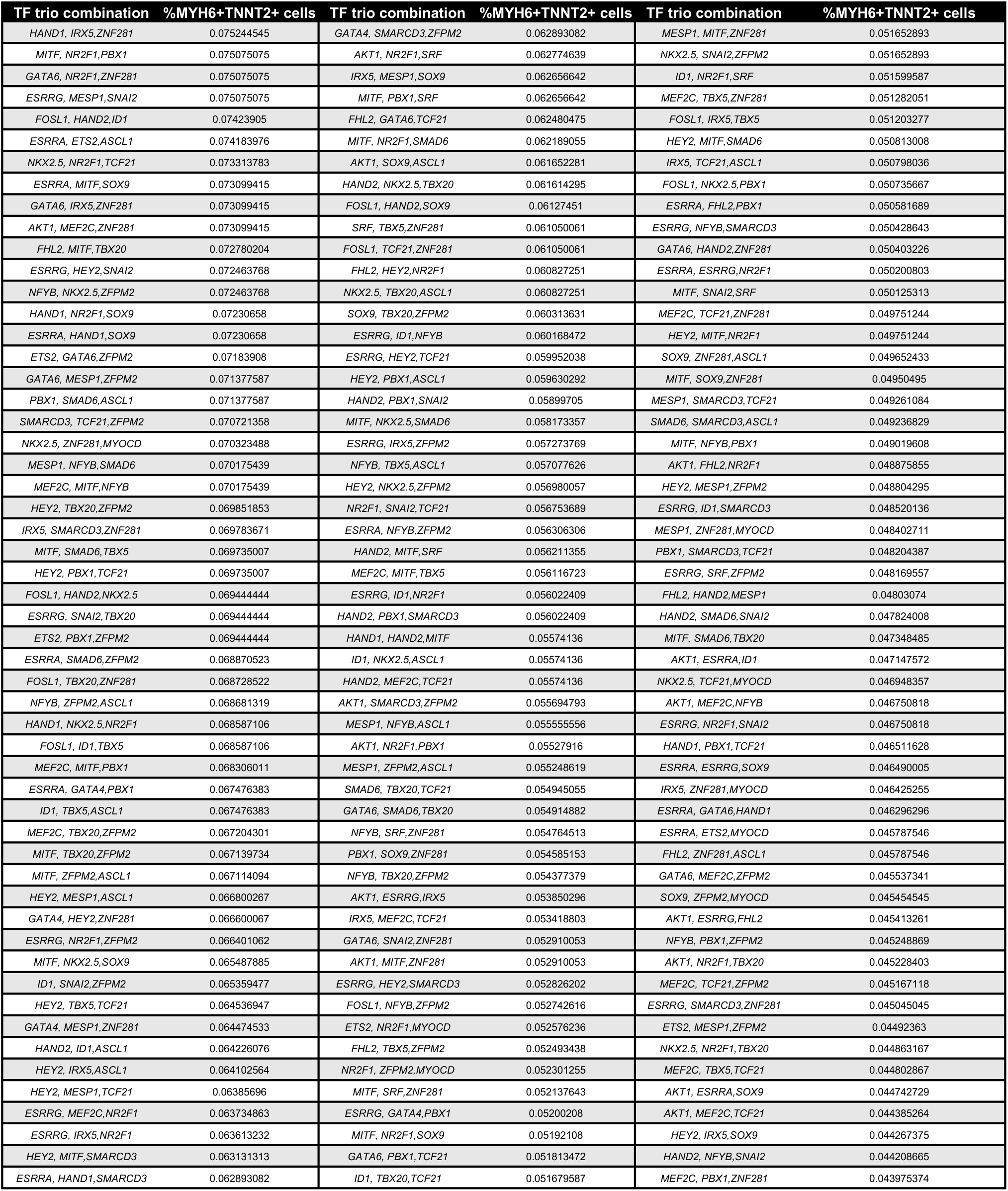
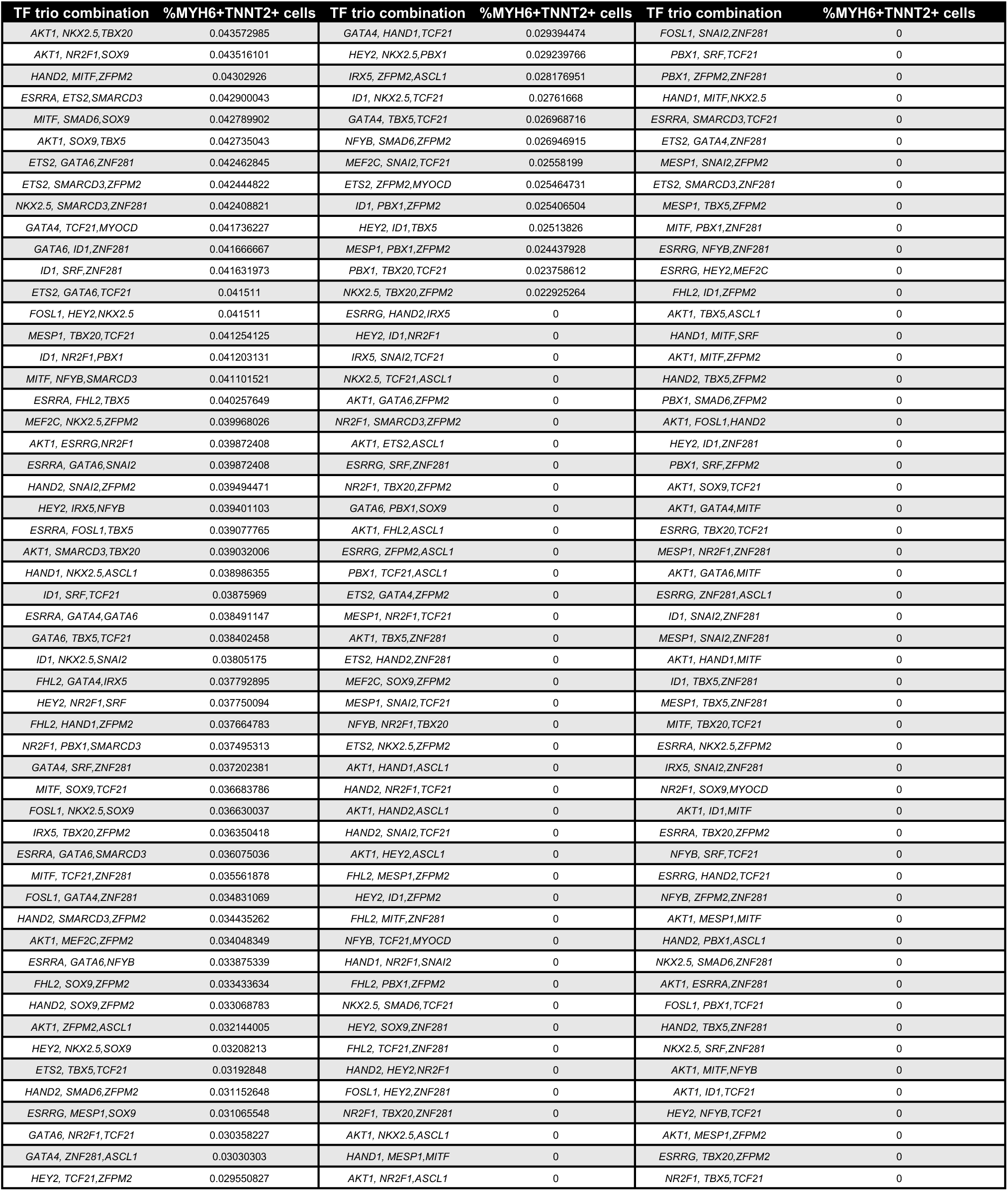
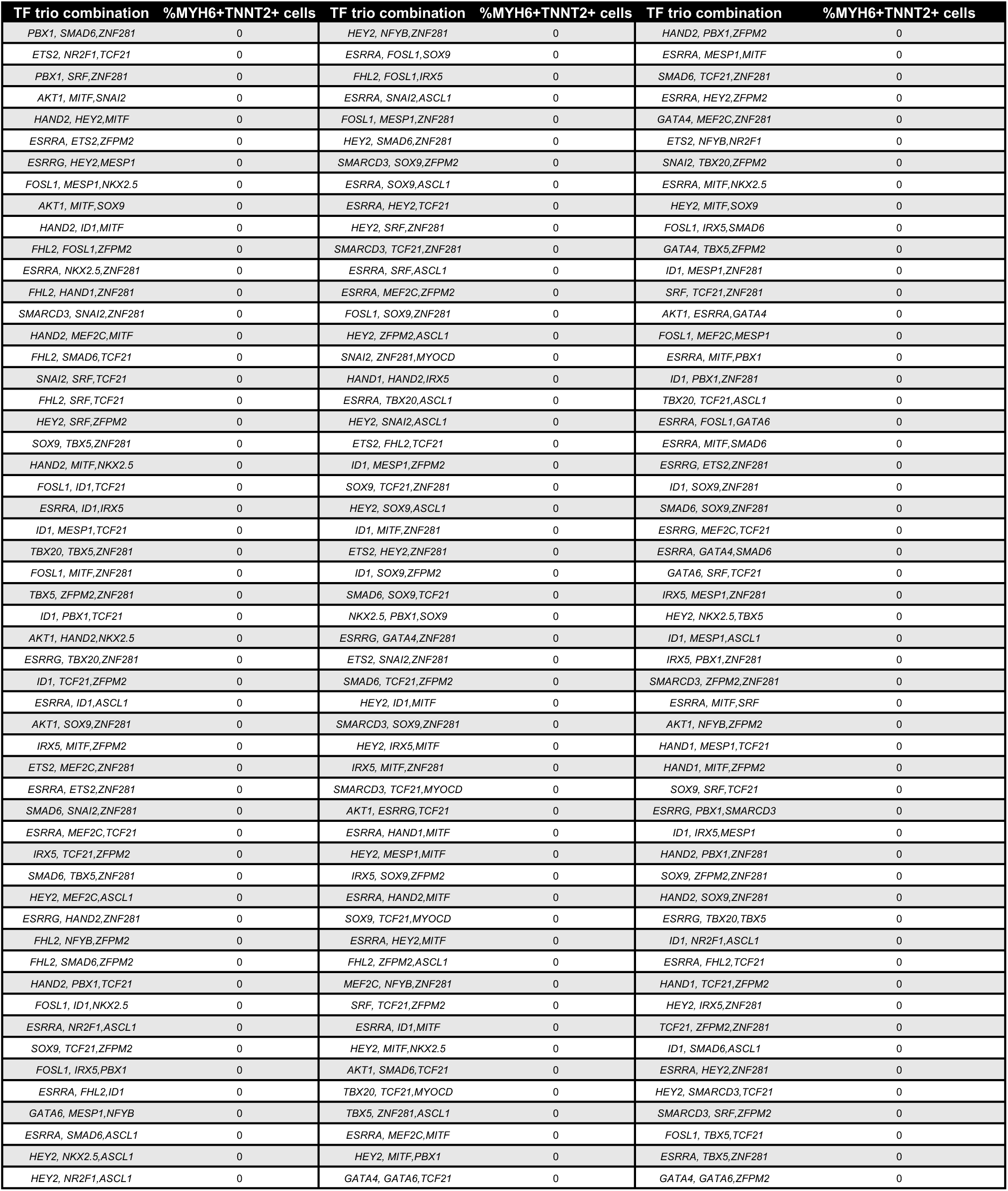
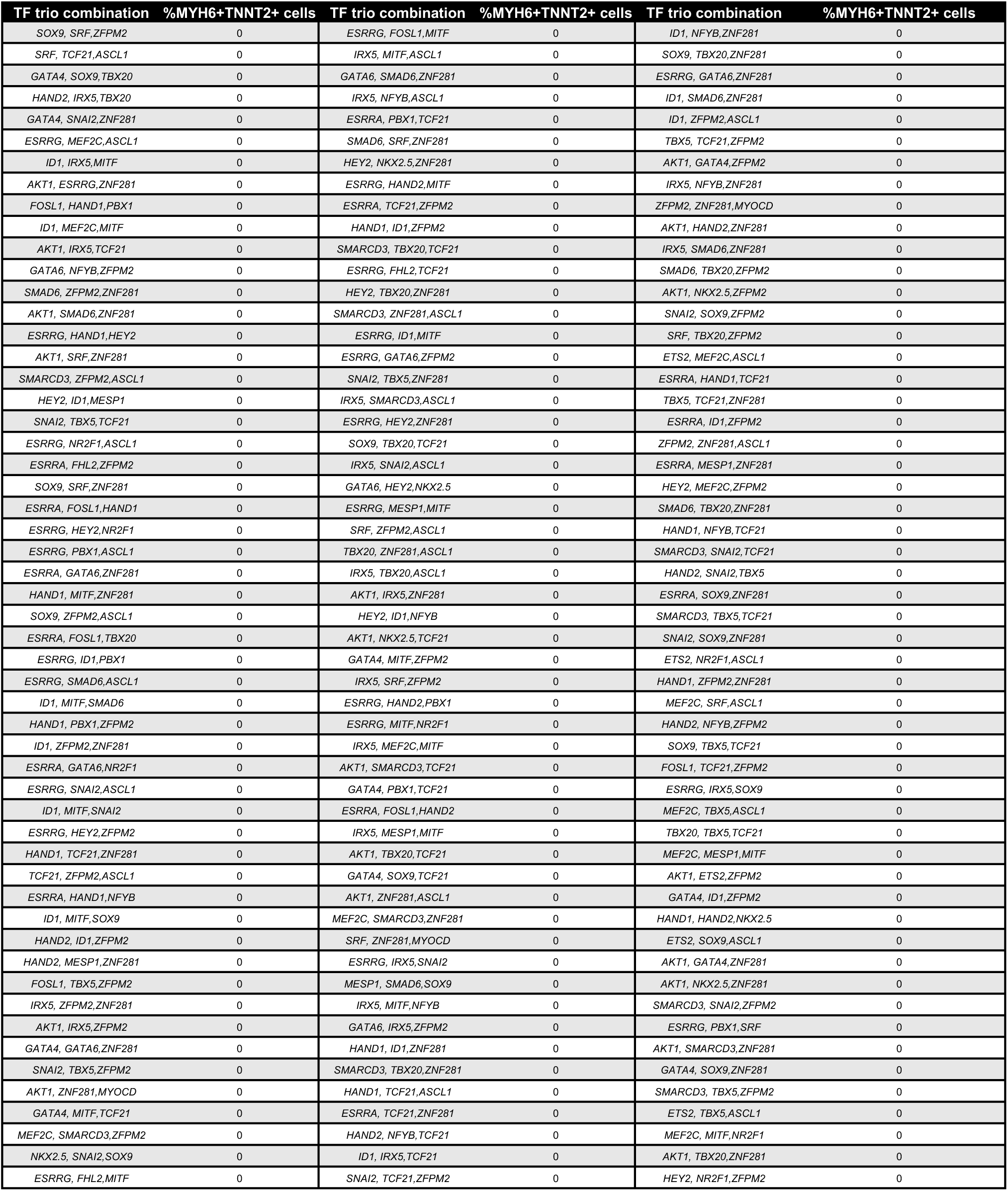
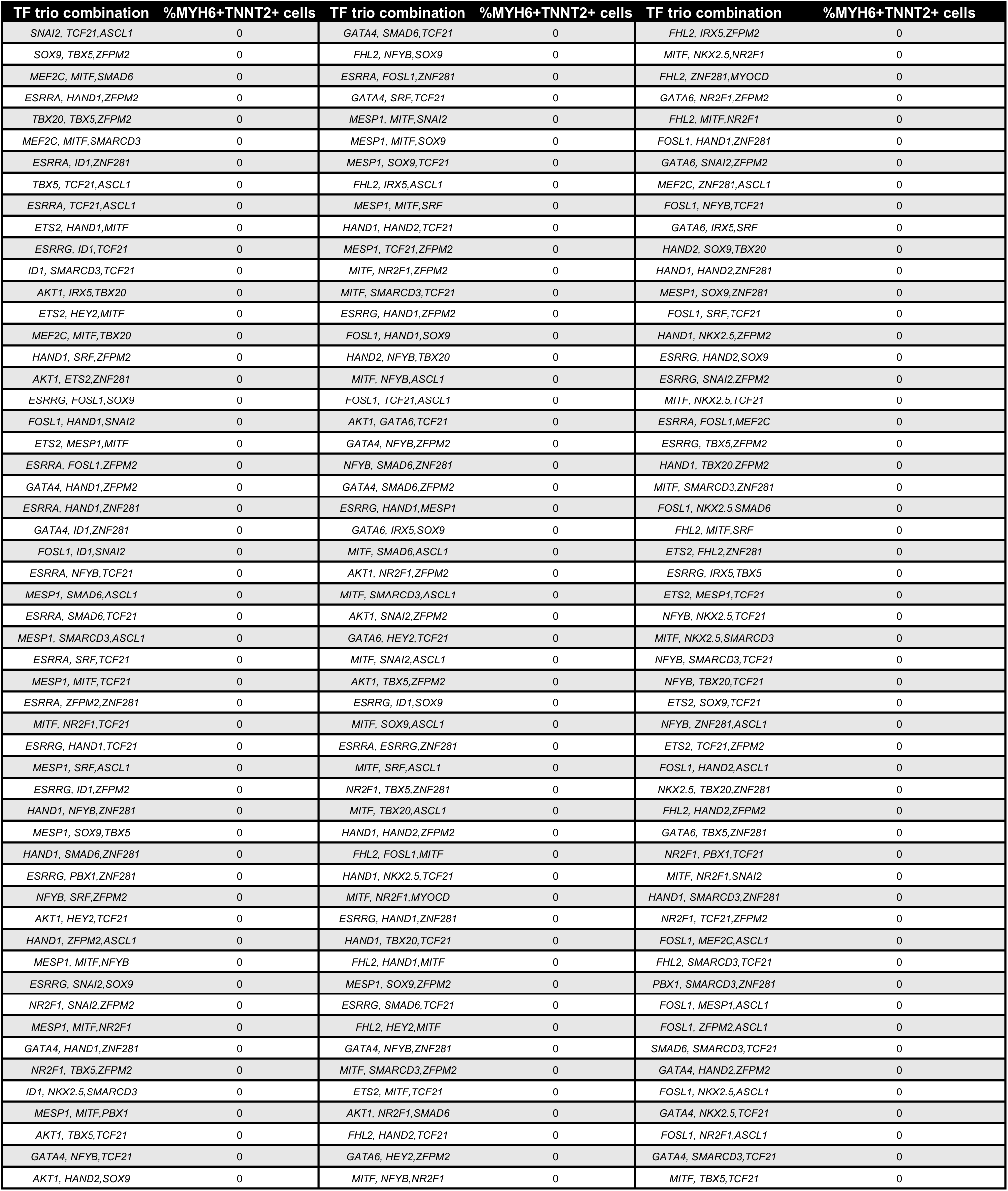
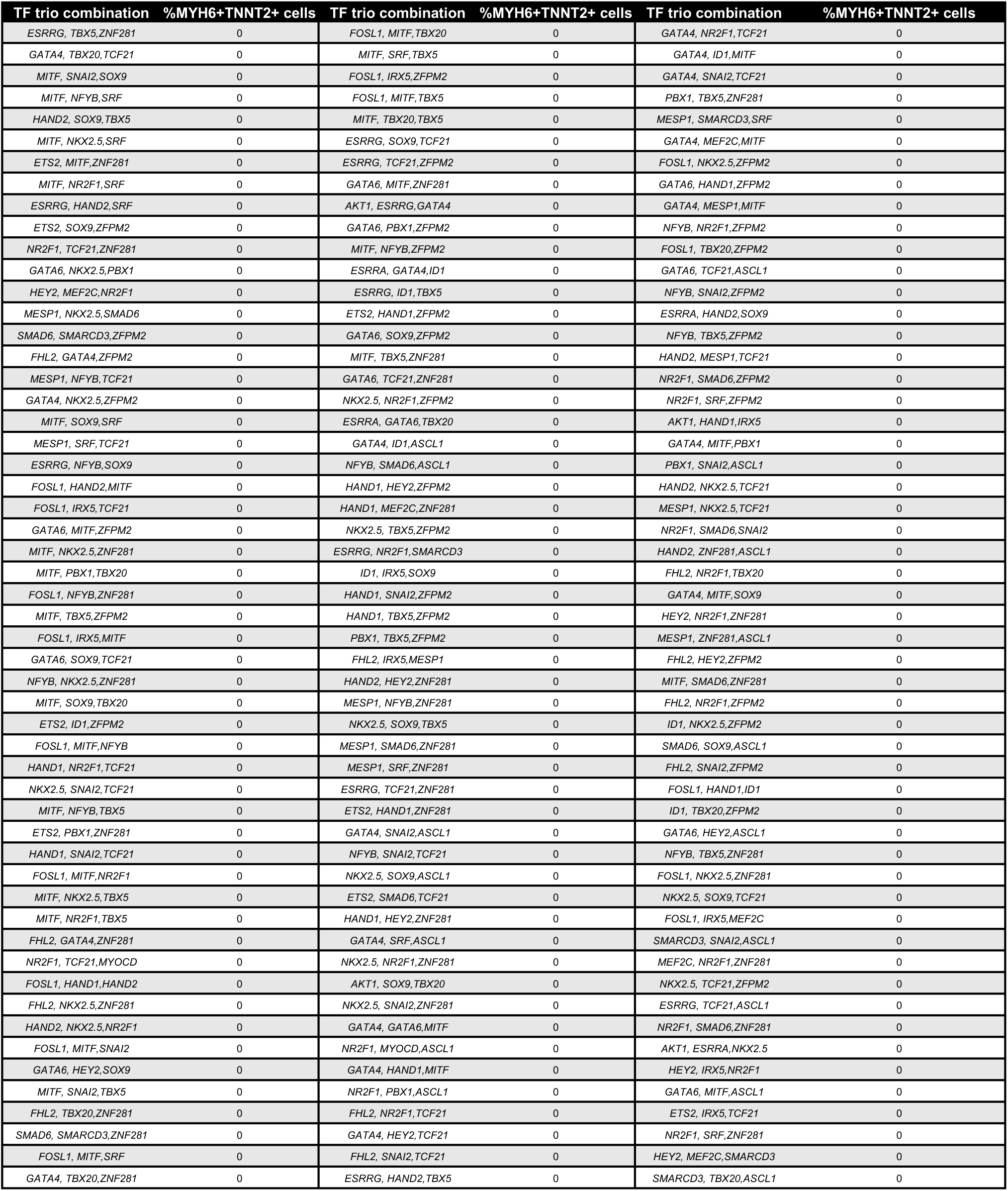
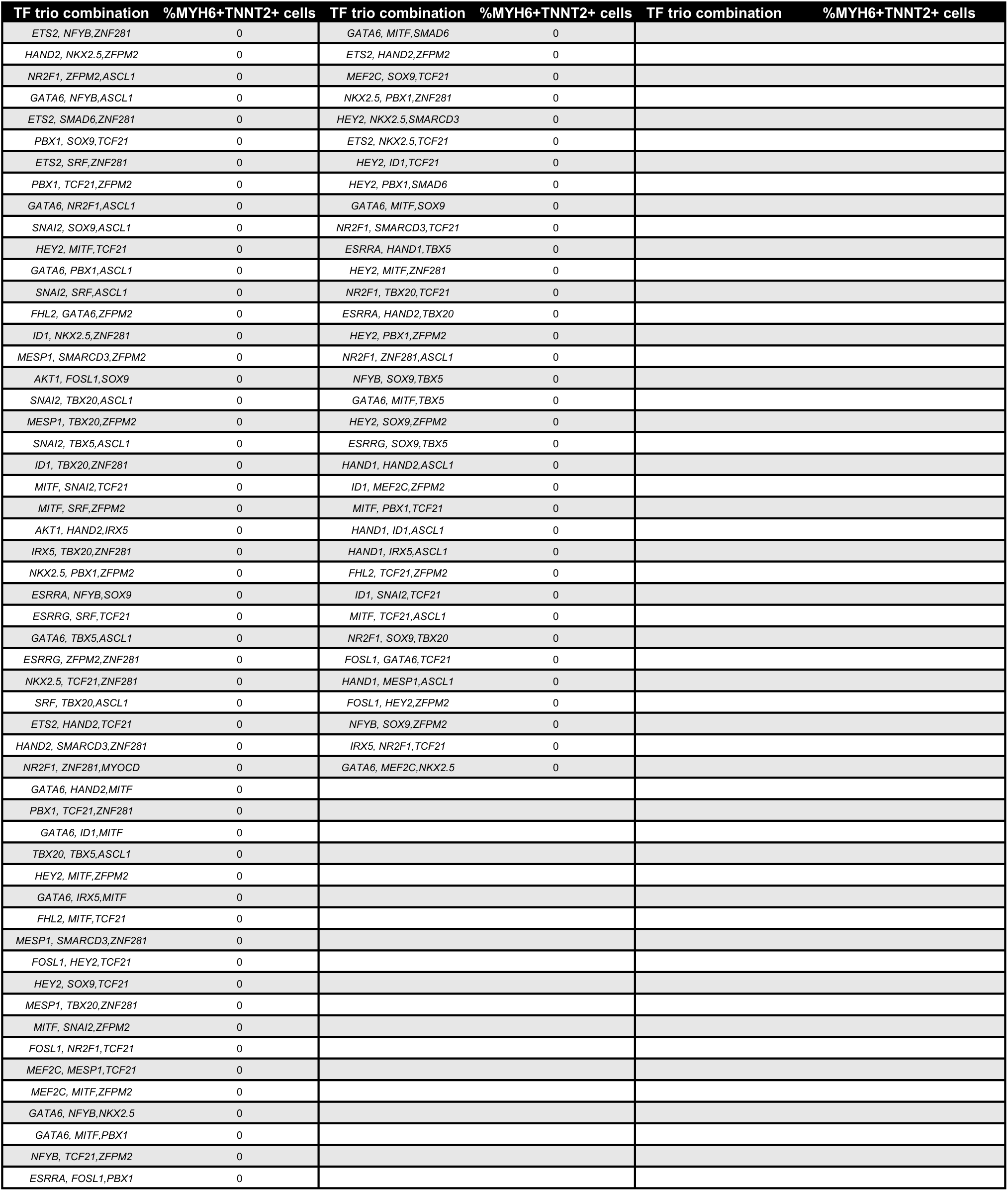
List of all 4,960 combinations tested and their respective reprogramming efficiency (MYH6^+^TNNT2^+^)

